# 340 dipteran genomes reveal the origin of Muller elements and sex chromosomes in Diptera

**DOI:** 10.64898/2026.06.01.729285

**Authors:** Julia Gries, Sam Ebdon, Aleksandra Bliznina, Joanna Collins, Christina N. Hodson, Thomas C. Mathers, Arif Maulana, Shane A. McCarthy, Michael Paulini, Dominic E. Absolon, Ksenia Krasheninnikova, Darwin Tree of Life Consortium, Mara K. N. Lawniczak, Richard Durbin, Kamil S. Jaron

## Abstract

Diptera genome evolution is traditionally viewed through the lens of *Drosophila melanogaster*. To provide an unbiased picture of chromosome evolution in Diptera, we reconstructed six ancestral linkage groups (ALGs) and their rearrangements using 340 chromosome-level genome assemblies from 59 dipteran Families and two outgroups. Notably, Diptera ALGs are conserved in many Nematoceran lineages, while emergence of Brachycera coincided with four chromosomal fissions, which subsequently fused in various combinations. In Schizophora and relatives, a stable karyotype of five metacentric chromosomes and a chromosome homologous to the dot emerged that is remarkably stable with the notable exception of Drosophila, where the metacentric chromosomes became acrocentric Muller elements. Furthermore, our reconstruction reveals an ancient sex chromosome system associated with a small and gene-poor ALG that frequently fused to other chromosomes.

## Introduction

The insect order Diptera, known as the true flies, consists of more than 152,000 species in 157 families with a last common ancestor estimated to have lived 240 million years ago (Mya, *1, 2)*. Our understanding of dipteran genome organization and evolution has traditionally been anchored to the model species *Drosophila melanogaster* (Drosophilidae, vinegar fly), which serves as the reference for many comparative studies (*3*–*5*). The chromosome arms of *D. melanogaster* are referred to as Muller elements A-F (*6*), and their gene content is conserved across many species of Drosophilidae (*7, 8*). Two important dipterans, *D. melanogaster* and *Anopheles gambiae* (Culicidae, mosquitoes), had reference genomes sequenced more than twenty years ago enabling the comparison of these species that diverged 240 Mya. The similarity of chromosomal gene content between *D. melanogaster* and *An. gambiae* led to the hypothesis of a highly conserved dipteran karyotype (*9*). This hypothesis was subsequently supported by the near-perfect conservation of gene content across 60-70 My of divergence between chromosomes in two schizophoran families, Drosophilidae and Tephritidae (fruit flies, *5*). Since then, Muller elements have been considered the ancestral karyotype of Diptera (*4, 10*–*12*). With hundreds of new dipteran chromosomal genome sequences, it is now possible to reevaluate whether Muller Elements are truly the ancestral dipteran chromosomes.

As *D. melanogaster* is a key model organism in biological and medical research and *An. gambiae* is a major malaria vector, studies of dipteran evolutionary genomics have typically focussed on Drosophilidae and Culicidae (*3*). In both lineages, intrachromosomal rearrangements are common but are largely restricted to individual chromosome arms. Additionally, metacentric chromosomes frequently exchange arms typically near the centromere via fission and subsequent fusion events (*8, 9*).

Muller element F, also known cytologically as the “dot” chromosome, has been proposed to correspond to the ancestral dipteran X chromosome (*4, 12, 13*), and even the ancestral sex chromosome of all insects (*14*). Multiple sex chromosome transitions from this assumed ancestral state have been observed across Diptera, using Muller elements as a reference framework (*4, 13*). However, the ancestral state of the dipteran sex chromosome has not been reconstructed explicitly and therefore its identity remains an open question.

In this study, we reconstruct the evolution of interchromosomal rearrangements in Diptera to define ancestral karyotypes as reference points for investigating chromosome evolution comparatively and describe the dynamics of both autosome and sex chromosome evolution at the order-scale.

### Genomic diversity of dipteran chromosome-level genome assemblies

We reconstructed Diptera ancestral linkage groups (ALGs) using chromosome-level genome assemblies of 340 dipteran species covering 59 families (**Fig. 1A**) the majority of which were assembled by The Darwin Tree of Life project (*15*). We inferred the phylogeny of these species using 5,067 Benchmarking Universal Single-Copy Orthologs (BUSCOs, diptera_odb12) present in a single-copy in at least 90% of the assemblies (**Fig. 1A)**. The resulting phylogeny was largely consistent with a previously published large-scale phylogeny (*16*, **Fig. S1**). Genome size varied ∼28-fold, from 71.55 Mb in *Smittia sp*. (Chironomidae), to 2.02 Gb in *Odontomyia ornata* (Stratiomyidae, **Fig. 1B**). Species representation varied across families (**Fig. 1C**), with some families being well sampled, while several species-rich families are represented by only a few genome assemblies. The haploid chromosome numbers ranged from n = 3 in *An. gambiae* (Culicidae) to n = 10 in *Villa cingulata* (Bombyliidae, bee flies, *17)*, with n = 6 being the most frequently observed haploid chromosome number (45%; **Fig. 1D**).

**Fig. 1:**
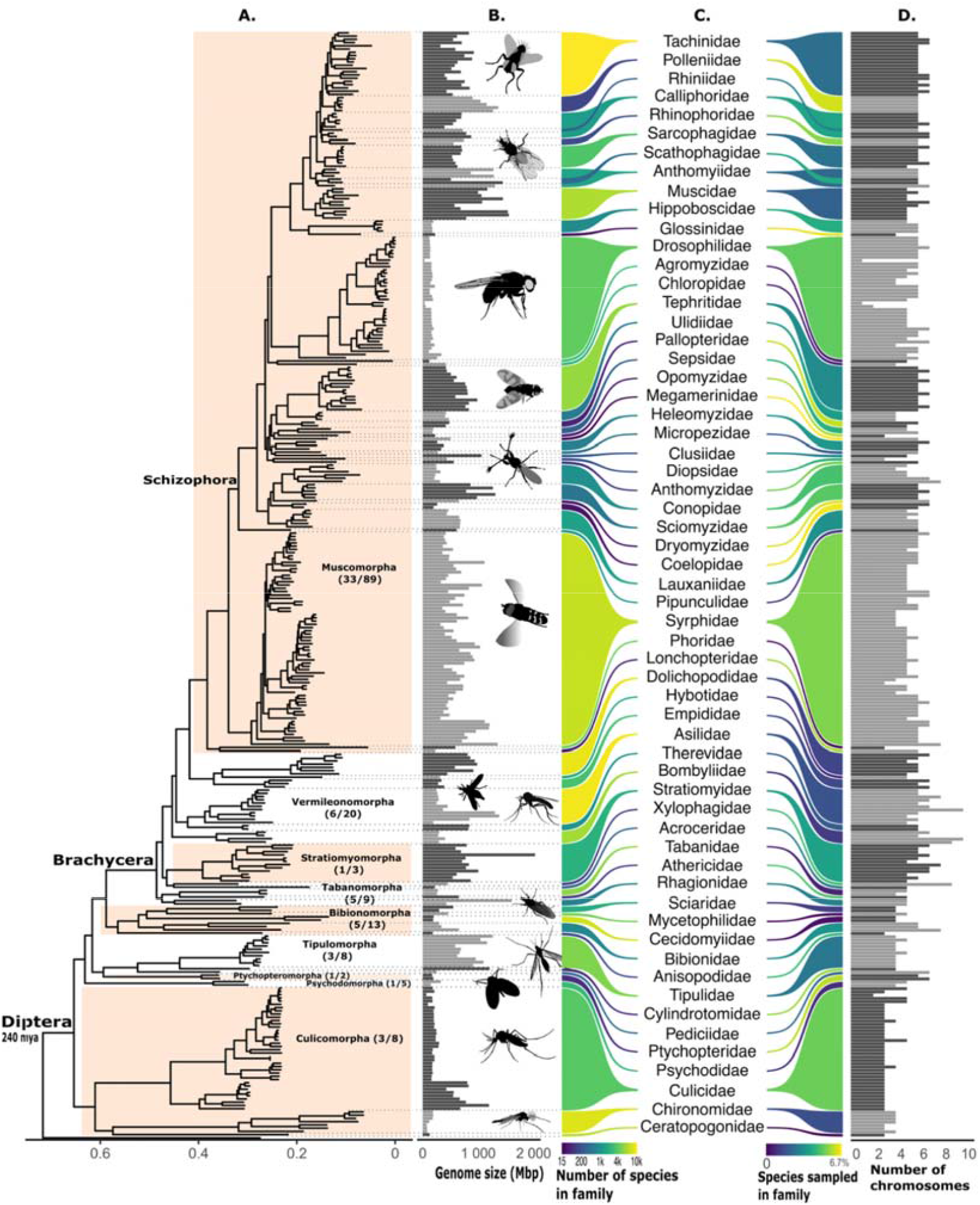
Phylogenetic relationships of the analysed genome assemblies. **A**. Species tree based on 5,067 BUSCO genes aligned with MAFFT and reconstructed with IQ-tree (Q.insect+I+G4) using *Panorpa germanica* (Mecoptera) as an outgroup. **B**. Genome sizes ranged from 71.55 Mb to 2.02 Gb. Shades of grey alternate between adjacent families. **C**. Sampled and described family diversity. The size of brackets represents the number of species for each family in our dataset. The colour of the left bracket represents the total number of described species per family and the colour of the right bracket indicates the percentage of described species sampled per family. **D**. Haploid chromosome counts range from 3 (excl. four assemblies with missing chromosomes) to 10. Shades of grey alternate between adjacent families. Tree annotations in panel A and species diversity data in panel C are from diptera.org.

### Three major karyotype transitions characterise early dipteran genome evolution

Before inferring Diptera ALGs, we excluded incomplete and highly rearranged genomes, including the outgroup species, that hindered reliable inference. This filtering resulted in a final dataset of 320 dipteran genome assemblies (**Supplementary Methods, Table S1, Text S1**). We inferred ALGs using syngraph (*18, 19*), an adjacency-based approach that infers linkage groups and interchromosomal rearrangements by leveraging the co-occurrence of loci on the same chromosome, irrespective of their order. With an initial syngraph analysis, we reconstructed Diptera ALG 1-5. As dipteran genomes show extensive chromosomal rearrangements (**Data S1)**, we applied a stringent marker threshold in the initial syngraph analysis to facilitate the inference of ancestral states. Specifically, at least 165 BUSCO genes (on average ≈ 18% of a chromosome) were required to move together for an event to be classified as a fission or fusion event (see Supplementary Methods). Many dipteran genomes contain a short chromosome, similar to the “dot” chromosome of *D. melanogaster* (*20, 21*). As expected, these chromosomes were not captured in our initial reconstruction due to the stringent marker threshold. To address this, we inferred ALG 6 separately. We reran our analysis exclusively on chromosomes that contained at least one BUSCO gene from Muller element F of *D. melanogaster* and had fewer than 150 BUSCOs in total. This includes many chromosomes that were not captured in the initial analysis, due to their low BUSCO counts. The inferred ALG 6 contains far fewer BUSCOs than the other ALGs and falls well below the previous detection threshold. A bootstrap analysis validated the high-confidence assignment of BUSCOs to the six inferred Diptera ALGs, hereafter referred to as ancestral karyotype α (**Fig. S2-S4**).

In some lineages, Diptera ALGs remained remarkably conserved even at the species level despite 240 My (*22*) of divergence, as exemplified by the genomes of *Bibio marci* (Bibionidae) and *Ptychoptera contaminata* (Ptychopteridae) in which the ancestral dipteran karyotype (α) is essentially unchanged (**Fig. 2A**). A major phylogenetically early transition was to the karyotype of the last common ancestor of all non-tabanomorphan Brachycera (190 Mya, *22*), which we designate as ancestral karyotype β. Finally, we reconstructed an ancestral karyotype in the last common ancestor of Asilidae, Therevidae, and Eremoneura (flies with three larval instars) that closely resembles Muller elements (160 Mya, *22*), referred to as ancestral karyotype γ. Strikingly, these three ancestral karyotypes (α, β, and γ) remained highly conserved in the ancestors of numerous dipteran families (**Fig. 2B, C**, see Fig. **S5** for a schematic overview). Below, we examine the extent of this conservation by focusing on changes shared across multiple families.

**Fig. 2:**
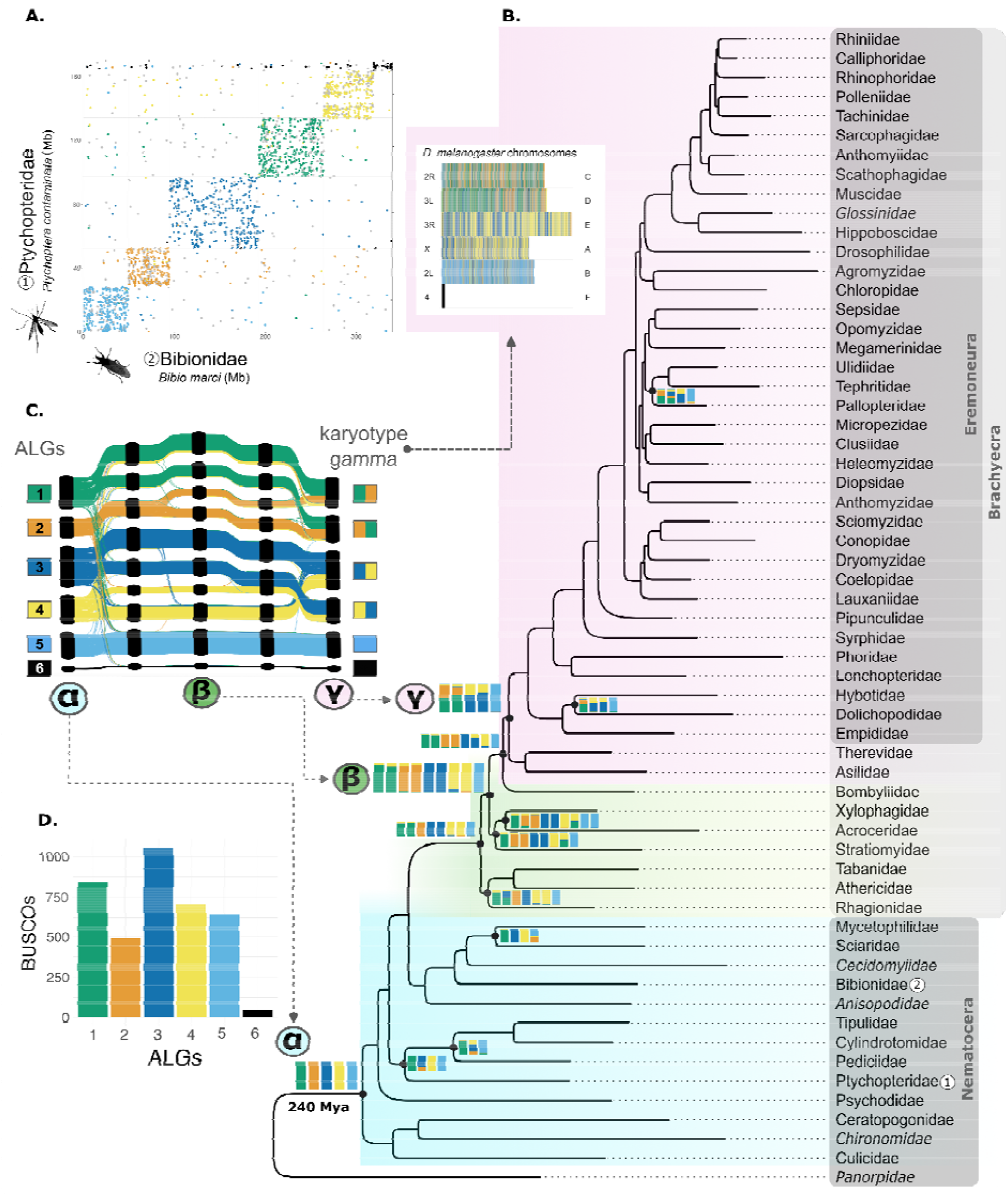
Reconstruction of genome evolution for 59 dipteran families. **A**. Conservation of Diptera ALGs in extant genomes. Dot plot of the genome assemblies of *P. contaminata* (Ptychopteridae) and *B. marci* (Bibionidae); exhibiting remarkable conservation of ALGs despite 240 million years of divergence (*16, 22*). Numbers indicate families in panel B. BUSCOs are coloured based on their assignment to Diptera ALGs. Unassigned BUSCOs are coloured in grey. **B**. Family level tree showing fissions and fusions reconstructed above the family level. Linkage groups at internal nodes are schematically represented as coloured rectangles of normalized height, with colours indicating assignment to Diptera ALGs; the corresponding numbers of BUSCO genes per ALG are shown in panel D. Fission events result in rectangles representing these additional chromosomes, whereas fusion events result in multi-coloured rectangles. Ancestral linkage groups in internal nodes without chromosomal rearrangements are not shown as they are well represented by the parental node. ALG 6 is not shown on the tree as it was reconstructed separately. Linkage groups were not inferred for families marked as italic, as they were excluded from the inference analysis due to poor assembly quality or extensive chromosomal rearrangements. (inset) *D. melanogaster* chromosomes are shown with BUSCOs coloured according to their assignment to ALGs. Left: chromosome arms; right: Muller elements. For further details on phylogenetic and linkage group reconstruction, see (**Text S3, Fig. S6**). **C**. Evolution of ancestral karyotypes. Ribbon plot depicting rearrangements of ALGs in the ancestral Dipteran lineage, tracing fission and fusion events from Diptera ALGs (ancestral karyotype α) through ancestral karyotype β to the last common ancestor of Asilidae, Therevidae and Eremoneura (ancestral karyotype γ), which closely resembles Muller elements. Nodes corresponding to these three ancestral karyotypes are indicated on the phylogeny. **D**. Numbers of BUSCO genes per inferred ALG.

Diptera ALGs (ancestral karyotype α) are conserved in the common ancestors of most nematoceran families. However, some nematoceran branches deviate from this karyotype. In the lineage leading to the common ancestor of Ptychopteromorpha and Tipulomorpha, two fusion events took place: first, ALGs 2 and 3 fused, followed by the fusion of this product with ALG 4, with the latter fusion being exclusive to Tipulomorpha **(Fig. 2B)**. Another exception is found in the common ancestor of Sciaridae and Mycetophilidae, where ALGs 2 and 5 fused. The highly rearranged genomes observed in Chironomidae and Cecidomyiidae prevented reliable inference and thus these were excluded from ancestral karyotype reconstruction (**Table S1; Data S1; Text S1**). We hypothesize that their ancestors likely also possessed extensively rearranged karyotypes.

Three chromosome fission events in ALGs 1, 3, and 4 led to the evolution of the karyotype in the last common ancestor of Brachycera (**Fig. 2C**). Brachycera initially bifurcates into the Tabanomorpha (represented by three families in our dataset) and all remaining brachyceran infraorders. The three tabanomorphan families (Rhagionidae, Athericidae, Tabanidae) share an additional fusion between parts of ALGs 1 and 3. Along the branch leading to the remaining Brachycera, a fission of ALG 2 gave rise to ancestral karyotype β of 10 linkage groups (**Fig 2B**) that subsequently fused in various combinations in descending lineages. The lineage leading from this node to Stratiomyidae, Acroceridae, and Xylophagidae is characterised by a fusion between parts of ALGs 1 and 4. In addition, Acroceridae and Xylophagidae both harbour a fission of ALG 5. It is intriguing that the initial rearrangement events in the ancestral dipteran lineage (α → β) coincide with the emergence of Brachycera, given their morphological distinctness from the paraphyletic Nematocera. Specifically, Brachycera are characterized by robust bodies and short antennae, whereas Nematocera typically exhibit slender bodies and long antennae (*23*). Whether these morphological differences relate to the underlying chromosomal rearrangements warrants further investigation in future studies.

Finally, four chromosome fusion events resulted in a third major karyotype (ancestral karyotype γ) in all muscomorphan families except Bombyliidae and Acroceridae. This group contains Asilidae, Therevidae, and the diverse clade Eremoneura, which includes well studied species such as *Musca domestica* (Muscidae) and *D. melanogaster* (**Fig 2B, C**). First, parts of ALGs 3 and 4 fused, a state conserved in the common ancestor of Bombyliidae. This was followed by a fusion of the remaining parts of ALGs 3 and 4, as well as two independent fusions of parts of ALGs 1 and 2 (**Fig. 2C**). The resulting karyotype (γ1-6, see **Fig. S5**) remained remarkably conserved across the majority of descendant family ancestors, including the common ancestor of Drosophilidae and Calyptratae. Consequently, karyotype γ closely resembles Muller elements, the chromosome arms of *D. melanogaster* (**Fig. 2B**). However, our findings indicate that Muller elements do not represent the ancestral karyotype of all Diptera, contradicting a widely accepted hypothesis. In Eremoneura, we identified two instances of deviation from ancestral karyotype γ. In the last common ancestor of Hybotidae and Dolichopodidae, the two linkage groups carrying parts of ALGs 1 and 2 fused. Additionally, in the last common ancestor of Pallopteridae, Tephritidae, and Ulidiidae, one linkage group carrying parts of ALGs 1 and 2 fused with a linkage group carrying parts of ALGs 3 and 4.

One ancestral linkage group, ALG 5, was conserved in all three ancestral karyotypes and across most common ancestors of dipteran families (**Fig. 2C**). Its conservation even extends to the species level. This is exemplified by the BUSCO gene content of Muller element B, which highly resembles that of ALG 5 (**Fig. 2B inset**). There are only two identified changes to the conservation of ALG 5 above the family level, a fusion of ALG 5 with ALG 2 in the common ancestor of Sciaridae and Mycetophelidae, and a fission of ALG 5 in the common ancestor of Acroceridae and Xylophagidae.

The reconstruction of Diptera ALGs 1–6 provides, for the first time, a framework for studying genome conservation and rearrangement across the order Diptera. Extensive intrachromosomal rearrangements, known from well studied dipteran species (*5, 8, 24, 25*), preclude reliable inference of gene order within ALGs.

### Ancestral karyotype γ is highly conserved at the species level

Whilst numerous new karyotypes have evolved from ancestral karyotype γ, we identify 91 species across 26 families that have retained the ancestral karyotype γ, and its linkage groups γ1-6 (**Fig. 3A, 5S**). We selected a representative species from each of these families to further investigate the extent of conservation using synteny. In all 26 species examined, four chromosomes are derived from Diptera ALGs 1-4, each formed by two discrete blocks corresponding to different ALGs in a bipartite organization (**Fig. 3A, 5S**), whereas the remaining two chromosomes each correspond to ALGs 5 and 6. In seven species with centromere annotations (*24, 25*), chromosomes with multiple ALGs are divided by the centromere into arms with distinct ALG identities, supporting the established role of centromeres as barriers to intra-arm rearrangements (*8, 26, 27*). An exception is presented by chromosomes derived from ALG 5, which contain BUSCOs from only a single ALG. In contrast, we observe that rearrangements within chromosome arms are frequent across selected species (**Fig. 3A**).

**Fig. 3:**
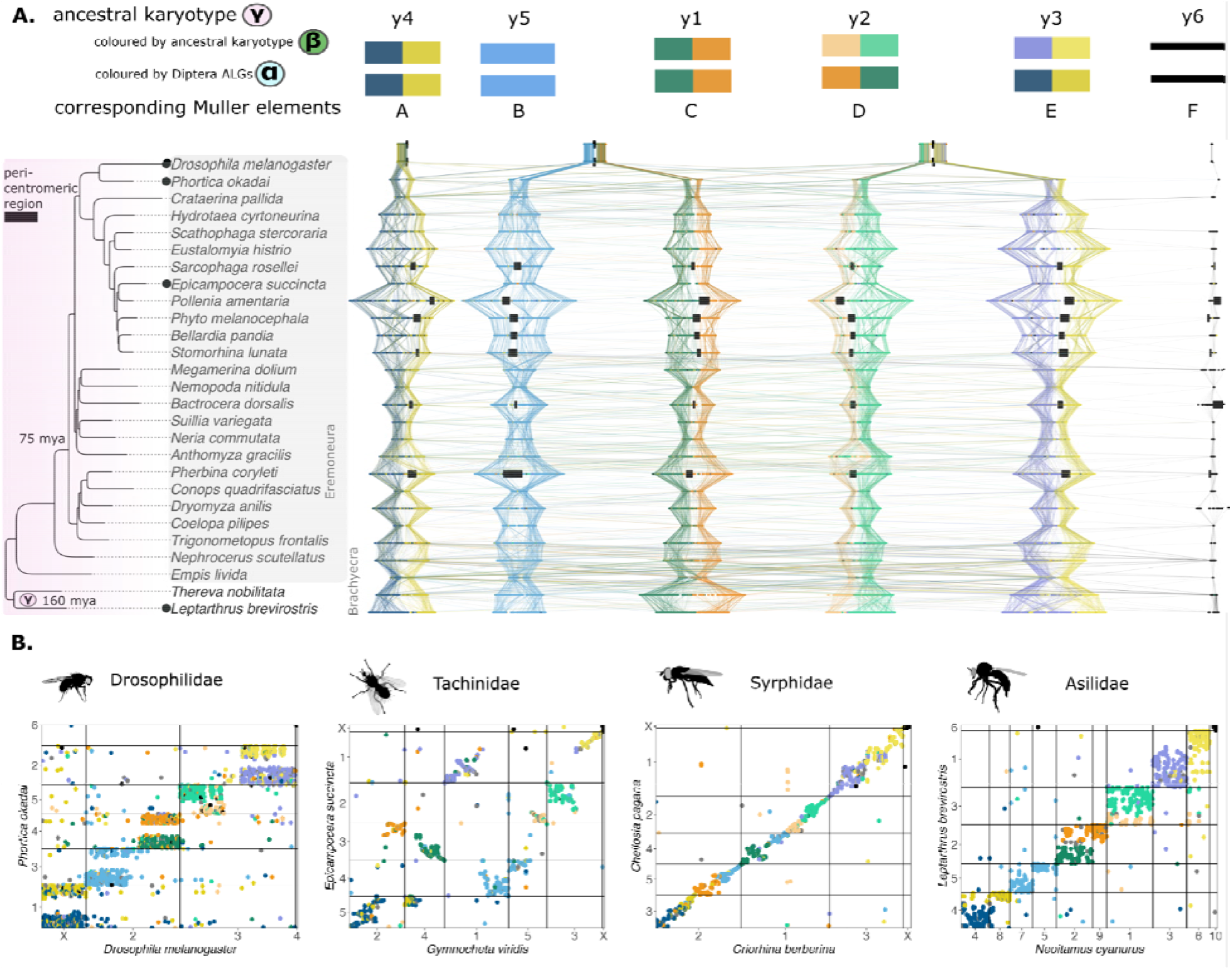
Conservation and rearrangements of ancestral karyotype γ across Asilidae, Therevidae and Eremoneura. **A**. Between family conservation of ancestral karyotype γ. The phylogeny shows a subset of dipteran species from 26 families, with genomes derived from ancestral karyotype γ, including the Therevidae, Asilidae, and Eremoneura. Each species represents a distinct family, except for *D. melanogaster* and *P. okadai*, which both belong to Drosophilidae. The genome of each species is displayed, with columns representing homologous chromosomes. The genome of *D. melanogaster* is stretched to enhance the visibility of its gene content. Genomes were selected based on their similarity to ancestral karyotype γ. Each genome contains 6 homologous chromosomes, with the exception of Muscidae, where a chromosome derived from ALG 6 is absent (*29*). BUSCO genes are coloured according to their assignment to ancestral karyotype β. In karyotype β, four ALGs underwent fission, hence different shades of color are used to highlight their distinct parts. Centromere annotations are available for seven genomes that harbor the ancestral karyotype γ and are indicated by black boxes (Rhiniidae: *Stomorhina lunata*, Calliphoridae: *Bellardia pandia*, Rhinophoridae: *Phyto melanocephala*, Pollenidae: *Pollenia amentaria*, Sarcophagidae: *Sarcophaga rosellei*, Tephritidae: *Bactrocera dorsalis*, Sciomyzidae: *Pherbina coryleti*). Additionally, the positions of centromeres in the genome of *D. melanogaster* are shown. **B**. Within family divergence of ancestral karyotype γ. Dot plots for four pairs of dipteran species are shown, with each pair comprising genomes of members of the same family. The chromosomes on the y-axis are conserved with respect to the ancestral karyotype γ and are indicated by black dots in the phylogeny in **A**. Syrphidae are not shown in **A**, as all members of this family share a fission of γ1 and γ5 followed by a reciprocal fusion of the resulting segments (**Data S1**). On the x-axis of the dot plots, the corresponding close relatives are shown, which possess derived karyotypes. Of these, only *D. melanogaster* is shown in the phylogeny in **A**.

To quantify how well centromeres divide chromosomes with two ALGs into distinct groups, we developed the ancestry division score (ADS), which scores chromosomes as fully intermixed to perfectly separated ALGs (0 to 1 respectively) (**Fig. S7**). Across the selected 26 species that retain the ancestral karyotype γ, ALGs remain consistently separated into distinct chromosome arms by the centromere despite more than 75 My of divergence (*16*). This effect is particularly strong for chromosomes homologous to γ1 and γ3, and weaker for those homologous γ2 and γ4.

As the chromosomes of all selected species in **Fig. 3A** are highly syntenic, we can assume that the remaining chromosomes without centromere annotations (except those derived from ALG 6) are also likely to be metacentric. Thus, parsimoniously, we hypothesise that ancestral karyotype γ consisted of five metacentric ALGs, and ALG 6 with an unknown centromere position. In the metacentric ALGs, the centromere likely separated chromosome arms with distinct ALG identities, reconstructed in ancestral karyotype β. This hypothesis is supported by karyotype data compiled in the Diptera karyotype database (*20*), where the majority of species with chromosome morphology annotations from Eremoneura, Therevidae, and Asilidae possess six chromosomes, with at least five being metacentric (**Fig. S8**).

*D. melanogaster* has long served as a model for comparative studies of chromosome evolution in Diptera, however, among the 38 Drosophilidae species in our dataset, only *Phortica okadai*, which represents the outgroup to the remaining Drosophilidae species, has retained the ancestral karyotype γ (**Fig. 3B**). In contrast to *P. okadai*, the majority of Drosophilidae species, including *D. melanogaster*, have different karyotypes (*8*). The correspondence of the linkage groups γ1-6 and Muller elements A-F is explicitly depicted in **Fig. S5**. Muller elements, the chromosome arms of *D. melanogaster*, are typically acrocentric or telocentric (centromeres near or at one chromosome end) across Drosphilidae, a feature previously hypothesized to reflect the ancestral state of the family (*8, 26, 27*). In some species, two acrocentric Muller elements are fused, and form large metacentric chromosomes. For example, in *D. melanogaster*, the haploid chromosome number is n = 4. Its genome contains two metacentric chromosomes (chr 2 = Muller element B and C, chr 3 = Muller element D and E), one acrocentric chromosome (Muller element A), and the small “dot” chromosome (Muller element F, **Fig. 2B, inset, 5S**).

Comparison of the chromosomes of *D. melanogaster* to those of *P. okadai*, a drosophilid which retained ancestral karyotype γ, reveals that each of the two large, metacentric *Drosophila* chromosomes evolved from the fusion of two independent pairs of ancestrally metacentric chromosomes (**Fig. 3B, S5**). The ancestral centromere structures separating chromosome arms were lost and replaced by a single new metacentric structure in the fused region. As a result, the genomic content of these new chromosome arms rearranged extensively relative to ancestral karyotype γ. Similarly, the acrocentric chromosome (Muller element A) exhibits a different organization, with intermixing of markers from two ALGs, likely reflecting disruption of the ancestral metacentric architecture (**Fig. S5**). These findings contradict the hypothesis that the six chromosome arms of Drosophilidae were ancestrally acrocentric (*8, 26*), and illustrate that using *D. melanogaster* as a point of reference fails to reveal the evolutionary history of chromosomes.

Alternative karyotypes derived from ancestral karyotype γ are also found outside of Drosophilidae. In total, 91 species from 26 families retained ancestral karyotype γ, while 152 species across 23 families show derived karyotypes. It remains unclear why closely related species sometimes maintain highly similar genomes, while others show extensive chromosomal rearrangements. Examples of rearranged genomes derived from ancestral karyotype γ, including the aforementioned *D. melanogaster*, are shown in **Figure 3B**. In *Gymnocheta viridis* (Tachinidae), we inferred rearrangements between all arms of the five ancestrally metacentric chromosomes. In Syrphidae, all species share a fission of Muller element homologs B and C and a reciprocal fusion of the resulting fragments (**Data S1**), maintaining a haploid chromosome number of n = 6 in most species. In *Criorhina berberina* (Syrphidae), however, the haploid number is reduced to n = 4 as a result of two additional fusion events. Unlike in *D. melanogaster*, the genomic content of these fused chromosomes does not appear to be intermixed, and the fusion may therefore be recent. In the Asilidae species *Neoitamus cyanurus*, nearly all metacentric chromosomes have undergone fission, resulting in a haploid karyotype of n = 10. Notably, even the typically conserved ALG 5 derived chromosome is fragmented in this species, with two distinct chromosomes each retaining a subset of ALG 5 associated BUSCO genes. Interestingly, ALG 5 is the only large linkage group of karyotype γ (γ5) that has been retained as a distinct chromosome. Overall, observed rearrangement patterns suggest that centromere position plays a role in shaping genome stability and rearrangement. Consistent with previous Drosophila work (*7, 8*), the centromere may act as a barrier that preserves the separation of ancestral chromosome arms, while intra-arm rearrangements are frequent (**Fig. 3A**). At the same time, centromeres of metacentric centromeres may serve as fission breakpoints enabling subsequent fusions into new combinations of ancestral segments, as exemplified by *G. viridis* (**Fig. 3B**). Conversely, loss or repositioning of centromeres may facilitate previously separated genomic regions to become extensively rearranged, as observed in *D. melanogaster*. However, the genome of *C. berberina* challenges this model, as it contains chromosomes with multiple well-separated ancestral segments. This either suggests relatively young fusions, or another mechanism preventing rearrangements.

These observations are consistent with the elementary algebraic operations proposed to underlie metazoan chromosome evolution (*28*). Conserved synteny without preservation of collinearity corresponds to the maintenance of homologous chromosome arms observed across species that retained ancestral karyotype γ (**Fig. 3A**). Robertsonian translocations, in which chromosomes or chromosome arms reversibly fuse without intermixing, resemble the rearrangements inferred in Tachinidae (**Fig. 3B**). Likewise, centric insertions, where one chromosome is inserted into another while retaining separation of genomic content, resemble the patterns observed in Syrphidae (see **Supplementary genome paintings**). In contrast, fusion-with-mixing, in which the genes of formerly separate chromosomes become irreversibly intermixed through extensive intrachromosomal rearrangement, characterize the genomic organization observed in Drosophilidae (**Fig. 3B, S5**). Together, these results suggest that centromere dynamics both constrain and facilitate chromosomal evolution, thereby contributing to the contrasting patterns of strong conservation and extensive rearrangement observed across Diptera.

### ALG 6 is the ancestral sex chromosome of Diptera

The gene content of Muller element F, the smallest chromosome of *D. melanogaster*, is conserved on homologous chromosomes across many dipteran families (*4, 14, 30*). Indeed, comparative genomic studies have suggested that Muller element F may be ancestral to the entire order Diptera (*4, 12*). Given that these homologous chromosomes are frequently X-linked outside of the Drosophilidae *(4, 14)*, it is therefore plausible that Muller element F is a descendant of the ancestral dipteran X chromosome. However, early cytological surveys across the dipteran phylogeny produced conflicting results regarding the presence of a “dot” chromosome and remained inconclusive (*21, 30*).

Consistent with previous genomic work, we found that the majority of sampled dipteran species possesses a chromosome homologous to Muller element F, which is a subset of ALG 6. With only 71 BUSCOs, ALG 6 has approximately 7-fold fewer BUSCOs than the next smallest linkage group, ALG 2, which contains 492 BUSCO genes (**Fig. 2D**). Likewise, ALG 6 derived chromosomes (i.e. chromosomes with >50% of ALG 6 genes) are typically shorter (Mann-Whitney U, p < 0.0001) and have fewer BUSCO genes (Mann-Whitney U, p < 0.0001) compared to other chromosomes (**Fig. 4A**). Chromosomes derived from ALG 6 vary dramatically in size; the smallest is 0.99 Mb (in *Drosophila busckii* (Drosophilidae)) and the largest is 140.14 Mb in *Merzomyia westermanni* (Tephritidae), a 140-fold difference (**Fig. 4B**). Nevertheless, they both carry only a handful of BUSCO genes (19 and 6 respectively). We estimated the ancestral length of ALG 6 to be 15.39 Mb using phylogenetic independent contrasts. Although the size varies, ALG 6 derived chromosomes remain conserved across the whole order, with minimal admixture between ALG 6 derived chromosomes and other ALGs (**Fig. 4C**).

**Figure 4:**
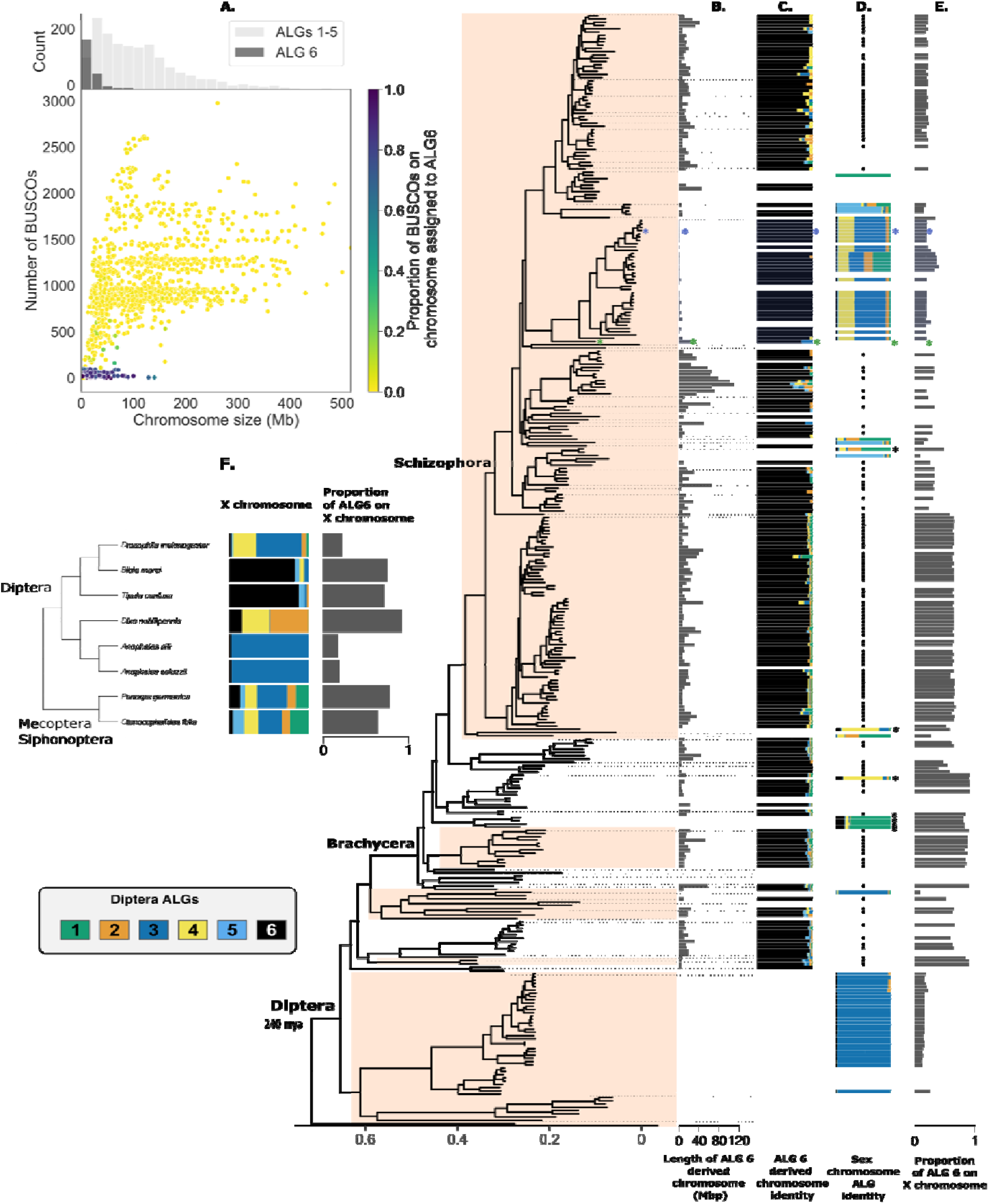
The evolution of ALG 6 derived chromosomes and sex chromosome transitions across the dipteran phylogeny. **A**. Scatter plot of BUSCO gene count per chromosome versus chromosome size. Dots ar coloured by the proportion of ALG 6 BUSCOs per chromosome. The histogram above the plot shows the distribution of chromosome sizes and the proportion of ALG 6 derived chromosome across size classes. ALG 6 derived chromosomes are the shortest chromosomes in dipteran genome assemblies and have the lowest BUSCO gene content. **B**. Sizes of ALG 6 derived chromosomes across dipteran genome assemblies. Empty positions indicate assemblies lacking an ALG 6 derived chromosome. Drosophilidae marked in blue. The reduced size and BUSCO gene content of Muller element F in Drosophilidae represents a derived condition specific to this family (blue asterisks). Notably, this reduction coincides with a sex chromosome turnover. The first species branching within Drosophilidae in this tree is *P. okadai* (green asterisk). **C**. BUSCO genes located on ALG 6 derived chromosomes, coloured by their assignment to ALGs. Chromosome sizes are normalized. **D**. BUSCO gene content of annotated sex chromosomes. Black dots indicate where ALG 6 derived chromosomes (shown in **C**.) were annotated as the sex chromosome. Empty positions indicate species in which the sex chromosome was not identified. Coloured bars indicate sex chromosomes other than ALG 6 derived chromosomes, with lengths normalized and colours reflecting BUSCO gene assignment to ALGs. Sex chromosomes with ALG 6 fusions are labeled with black asterisks. **E**. The proportion of all ALG 6 BUSCO genes on the X chromosome, for genomes with annotated sex chromosomes. **F**. Cladogram showing overrepresentation of ALG 6 BUSCOs on the X chromosomes across Diptera and in the outgroup species *P. germanica* (Mecoptera) and *C. felis* (Siphonaptera). *Dixa nubilipennis* is not included in the main phylogeny as its branching location was inferred later by fixing the core topology.

To characterise sex chromosome dynamics and identify sex-linked ALGs in Diptera we analyzed 222 genome assemblies from 41 families with annotated sex chromosomes (**Table S1**). We validated sex chromosome assignments in 50 species with reads available from male samples (see **Supplementary Methods, Table S5**) as male heterogamety is the typical sex chromosome configuration in Diptera (*20*). In all cases, the sex chromosome assignments reported in databases were consistent with coverage-based expectations (see **Supplementary Methods**). We therefore relied on database annotations for the remaining species.

To infer the ancestral sex chromosome of Diptera, we examined BUSCO genes located on the X chromosome in species with sex chromosome annotations. In 143 species (64.4% of species with annotated sex chromosomes), more than 50% of X-linked BUSCOs are from ALG 6 (**Fig. 4D**, indicated by black dots and shown in **Fig. 4C**). Additionally, in 7 species, the sex chromosome represents a fusion between ALG 6 and another chromosome (**Fig 4D**, labelled with black asterisks). In these cases, at least 50% of the genomes ALG 6 BUSCOs are still located on the sex chromosome (with six containing greater than 85%), even though the majority of BUSCOs on the sex chromosome are assigned to other ALGs. Overall, the majority of ALG 6 BUSCOs are concentrated on the X chromosomes in most species outside of Schizophora and Culicomorpha, although *Dixa nubilipennis* is a culicumorphan with most of ALG 6 BUSCOs located on the X chromosome (**Fig. 4E, F**). In Schizophora, we see a reduction of the X-linked ALG 6 BUSCOs to <40%. In these species, many ALG 6 BUSCOs have been redistributed to autosomes, despite the retention of an ALG 6 derived X chromosome in most lineages outside the Drosophilidae (**Fig. 4E**).

ALG 6 derived X chromosomes are found in numerous distantly related species indicating X-linkage in the common ancestor of Tipulomorpha, Ptychopteromorpha, and Psychodomorpha (**Fig. 4F**). Two additional observations suggest that ALG 6 was ancestrally sex-linked in Diptera. Firstly, within Culicomorpha, ALG 6 markers are co-located with different ALGs on the sex chromosomes in Dixidae and Culicidae, which suggests that ALG 6 likely existed as a sex-linked element in the ancestor of this group (**Fig. 4F**), and is further supported by independent fusion of ALG 6 in the third culicomorphan family Ceratopogonidae (**Fig. S9**). Finally, evidence that this conservation extends to the last common ancestor of Diptera is indirectly supported by two outgroup species. In both *Panorpa germanica* (Mecoptera, scorpionflies) and *Ctenocephalides felis* (Siphonaptera, fleas), the X chromosomes are substantially larger than those observed in Diptera. Yet, in both species the X chromosomes also contain the majority of ALG 6 BUSCO genes (**Fig. 4F**) which, in addition, are clustered toward one end of the chromosome, indicating a relatively recent fusion event (**Fig. S10**). Importantly, neither outgroup shows conservation of any other dipteran ALG besides ALG 6. These observations are consistent with previous reports of homology between Muller element F and the X chromosome of a related scorpionfly, *P. cognata* (Mecoptera) (*31*). Furthermore, cytological studies have shown three additional mecopteran species without reference genomes that possess a “dot”-shaped X chromosome, resembling the cytology of Muller element F in *D. melanogaster* (*32*).

Taken together, these results indicate that ALG 6 was the ancestral sex chromosome of Diptera, as suggested by previous studies, given Muller element F is derived from ALG 6 (*4, 12, 15*), similar to other systems with fewer sex chromosome turnovers such as cephalopods (*33*) and lepidoptera (*34*). Additionally, we hypothesize that ALG 6 may have already been sex-linked in the most recent common ancestor of Mecoptera, Siphonaptera, and Diptera, potentially as a relatively short, BUSCO-poor chromosome. However, definitive conclusions regarding the conservation and characteristics of this linkage group in deeper insect ancestors will require additional genome sequencing and robust sex chromosome identification across the class Insecta.

### The role of ALG 6 in sex chromosome dynamics

Across the analyzed dipteran genome assemblies, we identified 32 sex chromosome transitions (See **Text S4, Table S4**), which largely overlap with recently reported dipteran sex chromosome transitions (*12*). In 17 of these cases, the standalone ALG 6 derived chromosome was lost, either via loss of the sex determining function on the chromosome prior to the transition, presumably due to relaxed selection on maintaining the chromosome, or via chromosomal fusion (**Table S4**). The retention of sets of ALG 6 genes on most of these neo sex chromosomes suggests the latter is more likely (**Fig. 4D**).

The loss of the standalone ALG 6 derived chromosome is sometimes a clear case of chromosomal fusion as found in *Microdon myrmicae* (Syrphidae), *Atylotus latistriatus* (Tabanidae), or the common ancestor of Bombyliidae. In some cases, all chromosomes carry some of the ALG 6 genes, as found in Culicidae, Muscidae, or Chironomidae. Examples of both dispersed ALG 6 genes and chromosomal fusions are found in Culicomorpha, despite none of the species in our dataset featuring a distinct ALG 6 derived chromosome. In Culicidae and Chironomidae, ALG 6 genes are dispersed throughout the genome, while in Dixidae and Ceratopogonidae the ALG 6 genes are largely found on one end of a single chromosome derived from ALGs 2 and 3 respectively **(Fig. S9)**. This pattern indicates that ALG 6 was a standalone chromosome in the common ancestor of Culicomorpha that fused to different linkage groups in different lineages (**Fig. S11**), and was possibly followed by further BUSCO gene translocations in both Culicidae and Chironomidae. Therefore, the dispersal of ALG 6 genes across genomes might be the consequence of ancient fusion and subsequent rearrangements rather than the mechanism of chromosomal loss directly.

We identified three lineages with neo sex chromosomes that retained an ALG 6 derived autosome: In *Thecophora atra* (Conopidae) and all members of both Hippoboscidae and Drosophilidae. In the case of the Drosophilidae, the chromosome exhibits a strong reduction in content and size leading to the ‘dot’ morphology of the chromosome. Diptera ALG 6 contains approximately three times more BUSCO genes and the reconstructed size is approximately elevenfold longer than Muller element F in *D. melanogaster* (**Fig. S11**), demonstrating that the ‘dot’ morphology of Muller Element F is a derived condition (**Fig. 4B**). While this reduction is associated with sex chromosome turnover, the dispersal of the remaining ALG 6 BUSCO genes across the genome makes reconstructing the order of events difficult (**Fig. S12**). This may be ameliorated by denser sampling within Drosophilidae.

Finally, despite the fact that some species and families do not have annotated sex chromosomes, as they do not feature the ancestral sex chromosome (a distinct ALG 6 derived chromosome **Table S1**), we can infer there must have been a sex chromosome transition, even though in these cases we are unable to characterise the type of transition (fusion or turnover). This logic reveals 13 additional sex chromosome transitions, four of which are in Tabanomorpha, an infraorder without any previously reported sex chromosome transitions. Combining our newly found transitions with previously reported sex chromosome transitions we compiled the most up-to-date list of 45 transitions in 32 families (**Table S4**). The numbers of sex chromosome transition events found in Schizophora (19), non-Schizophoran Brachycera (14) and Nematocera (11) are roughly similar, challenging the previously suggested concentration of the number of sex chromosome transition events in Schizophora (*12*).

## Conclusions

Our analysis of the evolutionary history of 340 dipteran genome assemblies indicates a pattern of karyotype evolution. Large-scale rearrangements in the ancestral dipteran lineage occurred within a relatively short evolutionary window and transformed the stable ancestral karyotype α (Diptera ALGs) into the likewise stable ancestral karyotype γ (**Fig. 2B, C, 3A**). Notably, the occurrence of these ancestral rearrangements coincides with the emergence of Brachycera. Furthermore, the ancestral karyotype γ represents a stable combination of chromosome arms on metacentric chromosomes, maintained for 75 my (**Fig. 3A**). This finding contrasts the previous view, based largely on Culicidae and Drosophilidae, that chromosome arms frequently exchange between chromosomes rather than forming a preferential, stable constellation (*8, 9*).

Our detailed investigation of genome rearrangements in Diptera contradicts the hypothesis that the six chromosome arms of Drosophilidae were ancestrally acrocentric (*8, 26, 27*). Our findings illustrate that Muller elements are neither ancestral to the entire order Diptera, nor to Schizophora, and using them as a reference for synteny obscures important aspects of chromosome evolution, namely the deep preservation of nine chromosomal arms paired in a canonical set of metacentric chromosomes (**Fig. 3, S5**). In parallel, our analysis of Schizophoran chromosomal organization implicates centromeres as a major factor in retaining the identities of chromosomal arms; systematic characterization of centromeres across clades may reveal more unrecognized facets of genomic organization.

Sex chromosomes are typically the most conserved chromosomes within clades (*10, 33*) as well, the ancestral sex chromosome ALG 6 shows deep conservation, possibly even beyond the common ancestor of Diptera. However, in strong contrast to other systems, ALG 6 derived chromosomes dramatically vary in size and gene content, which might be linked to frequent sex chromosome turnovers (**Fig. 4, Table S4**). In addition to 13 known sex chromosome transitions, we identified 32 additional sex chromosome transitions, although for the majority of them the sex chromosome identity remains to be characterized. These transitions appear to frequently, but not always involve fusions with ALG 6, and in other instances more complex rearrangements have left genes canonically associated with ALG 6 dispersed across the rest of the genome.

By reconstructing major karyotype transitions over the evolutionary history of Diptera, we now have a framework to link patterns of conservation and change within and between families to family- or species-specific traits, which will be key towards revealing the underlying drivers of chromosomal evolution. Expanding genomic sampling across dipteran diversity, particularly within Nematocera, which is diverse and contains many species that have highly rearranged genomes, will be critical to refine this framework. This model of Diptera ALGs also provides a means to address a central question in genome evolution: why closely related species sometimes retain highly conserved karyotypes, while others undergo extensive chromosomal rearrangements.

## Glossary

ancestral karyotype α, β, and γ: Extant dipteran genomes derive from three ancestral karyotypes (α, β, and γ), which arose through large-scale chromosomal rearrangements early in dipteran evolution
ancestral linkage group (ALGs): Sets of genes that were located together on the same chromosome in a common ancestor
ALG derived chromosome: Chromosome of extant dipteran species that carries most BUSCOs of a distinct ALG
Muller elements: Chromosome arms of *D. melanogaster*
“dot” chromosome: Cytological feature of Muller element F in Drosophilidae and beyond

## Acknowledgements

We thank the many colleagues who have collected and identified specimens; assisted with access to sites; offered advice on legal, ethical, policy, and engagement issues; provided advice and support in extraction and sequencing; and worked with us in developing and improving our genome informatics. We would like to thank Beatriz Vicoso, Melissa Toups and Lorena Layana for presubmission manuscript sharing and helpful discussions. We would also like to thank Brian Charlesworth, Alex Mackintosh, Joana Meier, Mark Blaxter, Charlotte Wright, Alex Makunin, and the rest of the Tree of Life programme for helpful discussions and constructive criticism. This research was funded in whole, or in part, by the Wellcome Trust 220540/Z/20/A. For the purpose of Open Access, the author has applied a CC BY public copyright licence to any Author Accepted Manuscript version arising from this submission. The Darwin Tree of Life Project is funded by the Wellcome Trust through a Discretionary Award to the partnership (218328) and core funding to the Sanger Institute (206194), and by in-kind support from the partner institutions.

