## Supplementary materials for "340 dipteran genomes reveal the origin of Muller elements and sex chromosomes in Diptera"

†Collective authorship: <https://zenodo.org/records/4783559>

**The PDF file includes:**

Materials and Methods

Supplementary Texts S1 - S4

Figs. S1 - S12

References

**Other Supplementary Materials for this manuscript include the following:**

Tables S1 - S5

Data S1

### Materials and Methods

#### Reference genomes

We used GoaT (*35*) and the NCBI web interface to compile a list of available dipteran and mecopteran (as an outgroup) chromosome-level genome assemblies. As of June 2025, this list consisted of 341 dipteran genomes and one mecopteran genome. Most of the reference genomes (269) were sequenced by the Darwin Tree of Life Initiative (The Darwin Tree of Life Project Consortium 2022; See **Table S1** for the full list of assembly accessions and references); We downloaded the genomes using the NCBI datasets v16.13.0 tool (https://www.ncbi.nlm.nih.gov/datasets/). We included only one (the latest) assembly per species leaving us with 342 genomes used for the phylogenetic reconstruction. For each assembly, we used the primary haplotype only. Some genomes were excluded during the reconstruction of the ancestral karyotype (see below).

#### Phylogenetic Tree

We identified benchmarking single-copy orthologs in all the genomes (busco v5.8.2 using metaeuk and diptera_odb12 set of orthologs; see **Tables S2** and **S3** for a summary of BUSCO results) (*36*). We used a custom script busco2fasta.py (<https://github.com/lstevens17/busco2fasta>) to identify 5067 BUSCOs in the genome assemblies. We aligned sequences with MAFFT v7.525 (*37*) and trimmed them using trimAl v1.4.rev15 (*38*) with parameters -gt 0.8 -st 0.001 -resoverlap 0.75 -seqoverlap 80. A total of 4298 alignments passed the trimming step. We concatenated alignments into a supermatrix using a script catfasta2phyml.py (https://github.com/nylander/catfasta2phyml). We inferred the phylogenetic tree using IQ-TREE v2.4.0 (*39*) with autoselection of the substitution model (Q.insect+I+G4 substitution model with gamma rate variation) and 1000 ultrafast bootstrap replicates (*40*).

#### Reconstructing Diptera Ancestral Linkage Groups 1 - 5

We inferred the ancestral linkage groups of Diptera using a parsimony-based method syngraph (*19,* <https://github.com/Obscuromics/syngraph>). Syngraph progressively reconstructs ancestral linkage groups across a phylogenetic tree by minimizing the syntenic distance between two descendant nodes/tips and an outgroup node/tip (triplets of chromosomal states). We used a fissions and fusions (-r 2) model that defines the syntenic distance as the number of chromosomal rearrangements affecting more than a user-specified minimum number of gene markers (-m). It is possible to reconstruct the ancestral state when the number of chromosomal rearrangements does not exceed the number of chromosomes (*18*). The version of syngraph we used changes the order of resolving nodes where the topology does not impose a natural order of evaluation, i.e. pairs of unevaluated dichotomous splits (Fig S3). The current version evaluates all four possible combinations and first reconstructs the triplet with the minimal syntenic distance to the reconstructed ancestor and then proceeds in reconstructing the remaining triplets (Fig. S3). Given a choice of equally parsimonious reconstructions the first in the list is used. Given the limitation of the number of rearrangements per branch, we were unable to use the triplet approach to reconstruct the ancestral node of Diptera. That is because both of the available closest relatives have extremely rearranged genomes compared to Diptera (**Fig. S10**). Instead, we take the consensus of the last two reconstructed nodes (the common ancestor of Culicomorpha, and all remaining Diptera).

Before the inference we excluded Y chromosomes from all assemblies (See **Text S2** for genome modifications**)**. As Syngraph accommodates for missing data, we controlled for quality by manually checking the reconstruction node-by-node and iteratively excluding genomes which introduced considerable dropouts in marker-set propagation or collapses in inferred number of ALGs. Genomes were primarily excluded due to: 1) chromosomes missing from assemblies, or 2) insufficient marker counts and/or linkage group numbers per rearrangement (i.e. inadequate synteny, see **Text S1** and **Table S1** for a genome-by-genome outline of exclusions). While this led to the exclusion of some BUSCO-poor genomes, in numerous instances nodes were successfully reconstructed despite relatively low BUSCO counts (e.g. *Anopheles longipalpis* with only 73.2% BUSCOs in a single copy) providing the other genomes in the triplet were sufficiently marker-rich and syntenic. This round of filtering was used to explore the minimum number of markers required for a chromosomal rearrangement (parameter -m). Following authors’ recommendations we ran the analysis with a small -m, starting at 5, and gradually increased until the reconstructed number of ALGs stabilised with maximum marker propagation, which was for all values greater than 165 (tested up to 180). The number of ancestral linkage groups stabilised from approximately -m 140. This value corresponds to a threshold below which the number of rearrangements exceeds the number of chromosomes for a substantial portion of the tree, making ancestral state inferences impossible. Given the average marker density and genome size, -m 165 is equivalent to filtering for rearrangements larger on average than ≈ 18Mb, which is approximately ⅕ of the average chromosome length.

Using the optimal parameter setting (-r 2 -m 165), we further assessed the quality of reconstruction by manually inspecting individual nodes for unexpected ALG changes such as a single ALG, or rearranged karyotypes at nodes ancestral to conserved karyotypes (e.g. the common ancestor of Bibionomorpha given the remarkable preservation of dipteran ALGs in *Bibio marci*). The identified branches and individual genomes causing collapses of reconstruction were of two types. Firstly, issues of assemblies. In these cases we issued a correction when possible (See **Table S1** for details). Secondly, branches with a high number of rearrangements and sparse sampling, hindering inference of ancestral states. We initially iteratively excluded 20 genomes but corrected one of them, reducing the initial dataset of 340 to 321 dipteran species for the final inference (**Table S1**).

The final set of ALGs were based on 1000 bootstrap replicates of syngraph. In each replicate we resampled 5067 BUSCO markers with replacement and rerun syngraph reconstruction using the same finalized parameters (-r 2 -m 165) and 321 genomes. On average (and by definition) 63.21% of BUSCO markers were selected in each replicate and therefore the probability of co-occurence of any two markers in a bootstrap replicate is 39.96%. Using binomial distribution we calculated expected co-occurance of markers if they will have full consistency in reconstructions. Given the high number of theoretical pairwise comparisons (square of the number of markers), we chose 1e6 quantile of binomial distribution B(0.3996, 1000) as the threshold for the definition of ALGsTherefore, ALGs were then defined as sets of BUSCO genes that co-occured in the same linkage groups at least 327 times. This strategy resulted in a clear-cut definition of the file ALGs (Fig. S2).

#### Reconstructing Diptera ALG6

Given the well known ‘dot chromosome’ (Muller Element F) of *Drosophila melanogaster*, we considered that a smaller additional ALG was filtered out by Syngraph due to ignoring rearrangements smaller than 165 BUSCO genes (parameter -m). We expected chromosomes syntenic to a putative ALG 6 to be conserved and contain relatively few BUSCOs, like Muller Element F of *D. melanogaster*. We therefore searched for all chromosomes across the assemblies that: contained at least one of 21 BUSCO genes found on the Muller element F ( GCF_000001215.4, NC_004353.4) and contained fewer than 150 BUSCO genes. These criteria identified exactly one chromosome in each of 237 species and two chromosomes in one species (*Dolichopus virgultorum*). In all cases the chromosome had no clear ancestry related to the 5 ALGs previously identified by syngraph.

We retained this set of chromosomes, and reconstructed their evolution using the same approach as outlined above for ALGs 1-5 (using -m 1). This inference resulted in an ancestral reconstruction of a single chromosome with 78 genes. Seven of these (162912at7147, 164810at7147, 60829at7147, 75965at7147, 61998at7147, 173520at7147, 245796at7147) had conflicting assignments to ALGs 1-5. We conservatively chose to discard all seven genes, resulting in 71 BUSCO genes in ALG 6.

To investigate the size of ALG 6 we examined the length distribution of 212 ALG 6 derived chromosomes. We define a subset of these as ALG 6 derived chromosomes where >50% of markers belong to ALG 6. We obtained the lengths of all the chromosomes by indexing the genome assemblies with samtools v1.20. Finally, we estimated the length of ALG 6 in the last common ancestor, based on the chromosome lengths of extant species. To obtain this length we used Phylogenetic Independent Contrasts via the ace() function from the ape package v5.8.1.

#### Sex chromosome verification

We gathered information on previously identified X chromosomes from NCBI. To verify some of the assignments, we used the depth-of-coverage method (*41*). This method is well suited for detecting sex-linked regions that exhibit substantial sequence divergence. Sequencing reads, ideally obtained from both sexes, are aligned to a reference genome. Then the coverage ratio between males and females along the assembled sequence is calculated. If sufficient divergence has accumulated between the X and Y chromosomes, X-linked sequences will show approximately half the sequencing depth in males compared to females, while Y-linked sequences will exhibit male-specific coverage. However, this method is not suited to detect homomorphic sex chromosomes and is only applicable if read data from the heterogametic sex is available. We collected available sequencing reads from ENA. We mapped male PacBio reads to the matching genome assemblies using minimap2 v2.28 with parameter -ax map-hifi and sorted and indexed the BAM files using samtools v1.20. We calculated the read coverage depth with mosdepth v0.3.8.

### Supplementary Text

**Text S1: Genomes excluded from ancestral linkage group reconstruction**

*Drosophila gunungcola, D. willistoni, D. nebulosa, D. athabasca*, *D. affinis,* and *Anopheles rivulorum* were excluded given the assemblies were clearly missing whole or substantial portions of chromosomes. *Anopheles parensis* was excluded due to a lack of synteny between its focal triplet.

We excluded all Chironomidae as they are highly rearranged relative to the nearest relative in the dataset which diverges close to the common ancestor of all Diptera. However, all samples in the dataset are from the Chironomini subfamily, and so this node may be reconstructable if other subfamilies are sampled in addition and they feature a subset of the rearrangements we see in the Chironomini.

Lack of sampling in certain families of Bibionomorpha (Sciaridae, Cecidomyiidae, Bibionidae, Anisopodidae, Pleciidae) and Brachycera (Glossinidae) left it impossible to correctly reconstruct all rearrangements.

*Culicoides sonorensis, Forcipomyia palustris, Atylotus latistriatus,* and *Dixa nubilipennis* were not included in the reconstruction as their genomes were released after our inference and data freeze, but we have incorporated them into our analysis by painting their genomes with ALGs.

#### Text S2: Manual adjustments of individual genomes

The complete list of 11 excluded Y chromosomes is OZ020531.1, OV277353.1, OZ018405.1, OY770266.1, OX451349.1, OZ017744.1, OY744479.1, OX608056.1, OX596015.1, OW569400.1, OW121789.1.

The assembly of *Tolmerus cingulatus* (GCA_959613345.1) contained a Y chromosome (OY390709.1) with 387 BUSCO genes unique to the chromosome. We find the genome unlikely to miss so many BUSCO genes in females, therefore we considered this to be likely a labelling mistake, we therefore kept this chromosome for the inference.

The assembly of *Anopheles vaneedeni* (GCA_016170025.1) had only two chromosomes labelled, that were representing two chromosomal arms of the same chromosome. However, some of the longest scaffolds contained chromosome names in the headers and had matching expected size, we concluded that this must have been a submission error and labelled them as chromosomes.

In *Sicus ferrugineus*, Conopidae (idSicFerr1) the X chromosome is misannotated as part of a Y. We used additional sequencing data of a female, to confirm that a part of the Y chromosome represents an X chromosome instead. The genomic data does not support a connection of the previously identified X chromosome (which also shows the expected X chromosome coverage pattern), we therefore propose the system to be X_1_X_2_Y, while the final confirmation should be done using cytology. The updated version of the assembly is GCA_922984085.2.

#### Text S3: Dipteran phylogenetic reconstruction in the light of ALGs

We used the phylogeny from Wiegmann et al. 2011 (*16*) as a reference comparison, as this phylogeny considers the relationship between dipteran families rather than specific subfamilies, and to our knowledge, incorporates the largest number of molecular markers. This phylogeny was constructed using a partitioned maximum likelihood approach, with 202 taxa with data from ribosomal, mitochondrial and up to 12 nuclear protein-coding genes. In contrast, our phylogeny was constructed using a maximum likelihood approach from the set of 4298 BUSCO genes (all nuclear protein coding genes) in 340 dipteran genomes from 59 families. The topology of our phylogeny differed from that in (*16***, Fig. S1**), but both families retained the taxonomic clades of Brachycera, Eremoneura, Cyclorrhapha, Schizophora, and Calyptratae as monophyletic clades. These taxonomic clades are associated with morphological or life history features thought to have evolved once in Diptera (*3*).

Notably, several families with different phylogenetic placements in our family compared to Weigmann et al. 2011 (i.e. Xylophagaidae, Acroceridae, Hybotidae, Megamerinidae, Sepsidae, Micropezidae, Clusiidae) are families for which we had only one genome, and as such placement may be incorrect (although note for all of these families the reference phylogeny also had limited data). We view these families as those that most likely have an incorrect position in our phylogeny. We painted the genomes of individual flies with Brachycera ALGs (karyotype β, **Fig. 2C, Data S1**) to assess if the tree topology is parsimonious in respect to the number of chromosomal rearrangements. This analysis revealed some nodes in our phylogeny, as well as some in the reference phylogeny, which are potentially incorrect. The logic behind this assessment is that a fission splitting the same set of genes into two linkage groups in the same way is unlikely to occur multiple times in the evolutionary history of diptera. Moreover, fusions combining two linkage groups into one could potentially happen more than once in different branches of the phylogeny, but parsimony would suggest that the minimal number of these events is the most likely. Given this logic, we were able to reconstruct (albeit with some uncertainty) what we view as the most likely topology to the dipteran phylogeny (i.e. the minimal number of fissions and fusions between the same linkage groups, **Fig S1**). This phylogeny subtly differs from **Fig. 1** and resulted in indication that the individual chromosomal fissions and fusions (α → β → γ) happened sequentially (**Fig S6**). However, the finding that multiple fissions and fusions of linkage groups occurred in the Brachyceran lineage after the split from Bibionomorpha, but before the common ancestor of all Eremoneura, holds for both phylogenies (as well as the Weigmann et al. (2011) phylogeny).

#### Text S4: Sex chromosome transitions

Sex chromosome transitions were recorded in Bombyliidae, Syrphidae and Asilidae. In the common ancestor of the four sampled Bombyliidae species, the ALG 6 derived chromosome fused with a chromosome carrying BUSCO genes of ALG 1, forming the neo-sex chromosome. In one species of Syrphidae and Asilidae each of the ALG 6 derived chromosomes fused with the ALG 4 derived chromosome, forming a neo-sex chromosome.

A transition is found in *Teleopsis dalmanni* (Diopsidae). The ALG 5 derived chromosome is the X chromosome in this species, while the ALG 6 derived chromosome fused to a different chromosome, which carries BUSCO genes of ALG 1, 2, 3 and 4.

Additional transitions are present in Drosophilidae and Hippoboscidae. In the common ancestor of Drosophilidae a chromosome carrying BUSCO genes of ALG 3 and 4 became the sex-chromosome. Similarly in *Ornithomya chloropus* (Hippoboscidae) the ALG 5 derived chromosome became the X chromosome. In both cases, the ALG 6 derived chromosome remained unchanged and became an autosome. Additionally, both families include species in which the already derived sex chromosome was turned over once more due to a fusion with an autosome. Multiple Drosophila species share this neo-sex chromosome which resulted through a fusion of the derived sex chromosome with an autosome carrying BUSCO genes of ALG 1 and 2. In Hippoboscidae, the derived sex chromosome fused to a chromosome carrying BUSCO genes of ALG 1 and 2. For the two families Drosophilidae and Hippoboscidae, these two consecutive sex chromosome transitions are the only ones in the present dataset.

Conopidae (thick-head flies) stand out as the family with the highest frequency of sex chromosome transitions. Among the four species with annotated sex chromosomes, each exhibits a distinct sex chromosome configuration. In *Conops quadrifasciatus*, the ALG 6 derived chromosome is retained as the sex chromosome. In *Myopa testacea*, the ALG 6 derived chromosome fused with a chromosome carrying BUSCO genes from ALG 1 and ALG 2, forming a neo-sex chromosome. This fusion is likely ancestral to both *M. testacea* and *Thecophora atra* as it is present in both species. In the latter, however, the derived chromosome carrying BUSCO genes of ALG 6, 1, and 2 subsequently fused with an additional chromosome carrying BUSCO genes from ALG 3 and ALG 4. However, this composite chromosome is not sex-linked in *T. atra*; instead, the ALG 5 derived chromosome is annotated as the sex chromosome in this species. In *Sicus ferrugineus*, two X chromosomes are identified: one derived ALG 6 and a second carrying BUSCO genes from ALG 3 and ALG 4. In addition, a single Y chromosome is annotated, potentially pairing with both X chromosomes. The remarkable sex chromosome diversity observed among this small number of sequenced species suggests that even greater, as yet undiscovered, diversity exists within Conopidae, warranting further investigation.

We observed several sex chromosome transitions in species with very small sex chromosomes. In a member of Muscidae, *Polietes domitor,* the X is extremely BUSCO poor, but it does not carry any ALG 6 BUSCO genes and only has a single BUSCO gene. Muscidae have documented sex chromosome system diversity including sex chromosomes homologous to Muller element F (*29*). This example might represent a more dramatic reduction of ALG 6 than Drosophilidae or a completely new dot-shaped chromosome.

In *Dolichopus virgultorum* (Dolichopodidae) ALG 6 split into two chromosomes, both of which are X chromosomes, representing a fission. However, *D. virgultorum* is the only member of Dolichopodidae with an identified sex chromosome; all other members of the family feature a single intact ALG 6 derived chromosome, and therefore this is a recent fission or possibly an assembly artifact.

The ALG 6 derived chromosome was lost in species of the following families, with its BUSCO genes dispersed across the genome and arisal of a neo-sex chromosome: Sciaridae, Culicidae, Clusiidae, Lonchopteridae and Glossinidae. In Sciaridae and Culicidae, the ALG 3 derived chromosome is the X chromosome. The observed transition in Sciaridae might be related to a number of other unusual reproductive features observed in this family (*42*). In some species of Culicidae, subfamily Culicinae (genera *Aedes* and *Culex*), the homologous ALG 3 derived chromosomes form a homomorphic sex chromosome pair (*43*). In the subfamily Anophelinae only a fraction of ALG 3 BUSCO genes are present on the sex chromosome. This is probably due to a fission of ALG 3, only ancestral to the subfamily Anophelinae.

In *Clusia tigrina* (Clusiidae) and *Lonchoptera lutea* (Lonchopteridae), a chromosome harbouring a fusion between parts of ALGs 1 and 2 (corresponding to Muller element D) became the sex chromosome and finally, in Glossinidae it is part ALGs 3 and 4 (Muller element A).

**Additional transitions to those outlined above:**

Simple fusions

- In *Lutzomyia longipalpis* ALG 6 is fused to ALG 1.
- In *Dixa nubilipennis* the X chromosome has a large portion of ALG 6 fused to ALG 2.
- In *Culicoides sonorensis* ALG 6 is fused to ALG 3 independently of the Culicidae, although the identity of the sex chromosome is unknown.
- In *Leptogaster cylindrica* ALG 6 is fused to part of ALG 3.
- In *Atylotus latistriatus* and *Atherix ibis* ALG 6 fused to part of ALG 4.
- *Holcocephala fusca* is the only example of ALG 6 fusing to part of ALG 5.

Complex fusions

- In Chloropidae ALG 6 is fused to ALGs 1 and 2.
- In *Liriomyza trifolii* ALG 6 is fused to parts of ALGs 1 and 2.
- In *Sylvicola cinctus* ALG 6 fused to ALGs 1 and 3.
- In *Xylophagus ater* ALG 6 is fused to small pieces of ALGs 2 and 3.
- In *Chrysops caecutiens* ALG 6 is fused to ALG 2 and part of ALG 4
- In *Acrocera orbiculus* ALG 6 is fused to parts of ALGs 2 and 4.
- In *Dioctria linearis* ALG 6 is fused to part of ALGs 3 and 4.
- In *Opomyza florum* ALG 6 is fused to parts of ALGs 1, 2, 3, and 4.

Recorded by Muller element

- In *Ephydra hians* and *Themira minor* X chromosome is Muller Element D.
- In *Tephritis californica* X chromosome is reportedly Muller Element F
- In *Musca domestica* X chromosome corresponds to Muller Elements A and F.
- In *Stomoxys calcitrans* X chromosome corresponds to Muller Elements D and F.

Unclear

- Cecidomyiids are typically monogynous with two X chromosomes with many ALGs.
- Chironomids sex chromosomes unknown?
- Simulid sex chromosomes are unknown but have turnovers.
- *Megaselia abdita* and *M. scalaris*

### Supplementary Figures

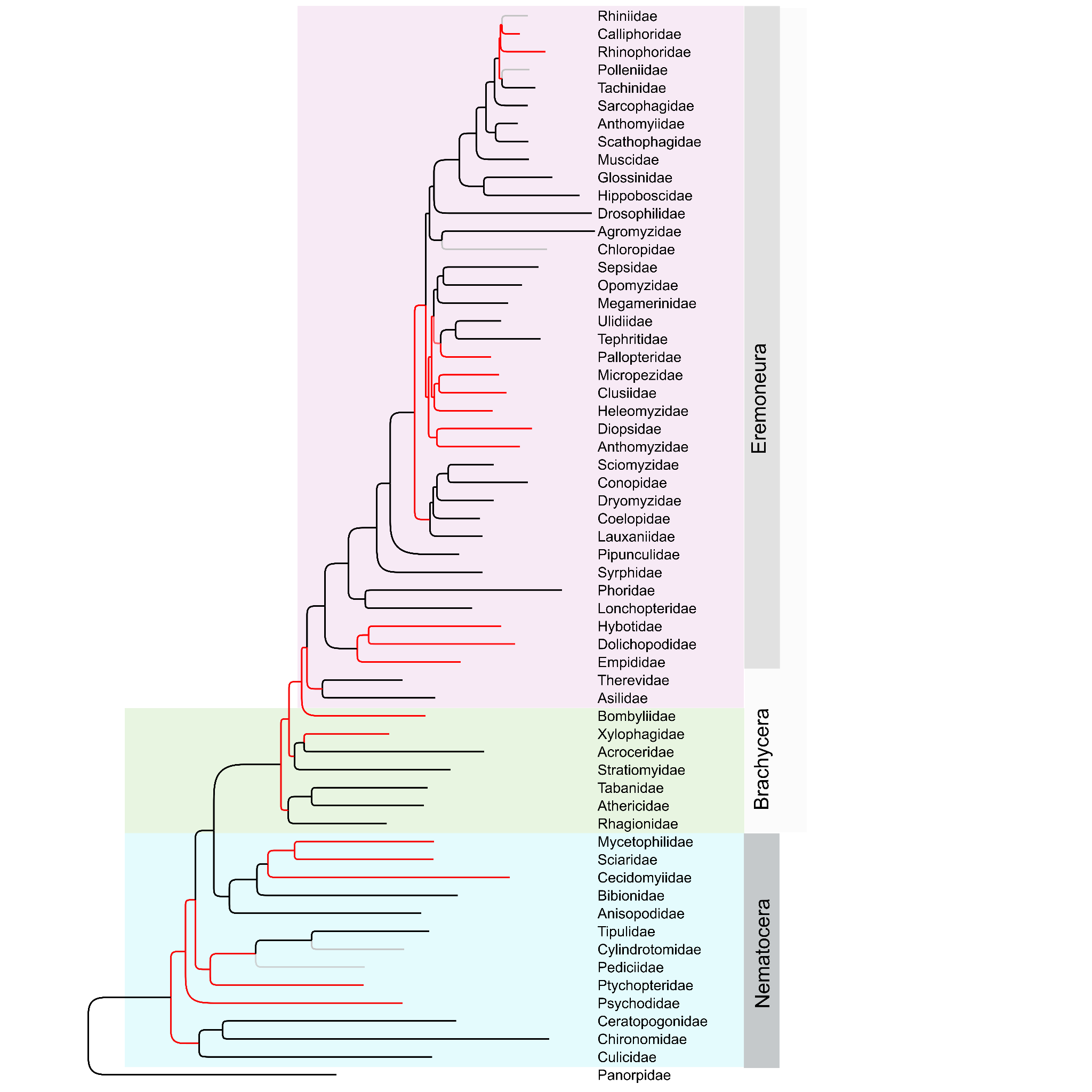

#### Fig. S1: Comparison of the topology of the phylogeny in this paper with a widely used published study.

Our phylogeny based on 4298 nuclear BUSCOs is compared to Wiegmann et al. 2011 (based on up to 12 nuclear protein coding genes, 18S and 28S, and the complete mitochondrial genome) (*16*). Branches coloured red have a different topology compared to Wiegmann et al. (2011), while branches in grey are associated with families absent from Wiegmann et al. (2011). Background colours are associated with the major karyotype changes in dipteran evolutionary history outlined in Figure 2.

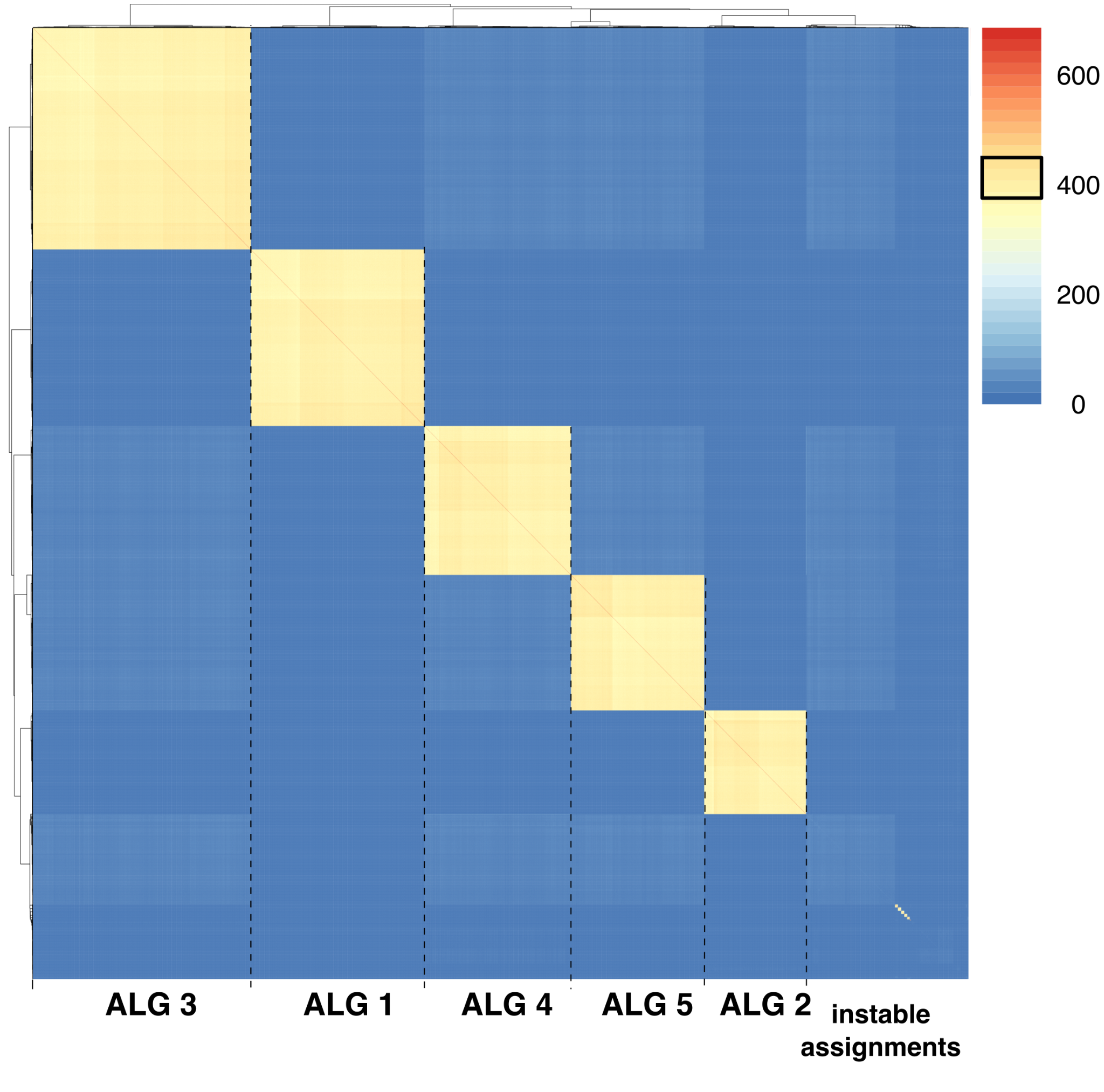

#### Fig. S2: Bootstrapping support of co-assignments of BUSCO genes in ancestral linkage groups.

Co-assignment support of 5067 BUSCO genes into inferred ALGs over 1000 bootstrap replicates of syngraph. The square on the legend corresponds to 0.01 - 0.99 quantiles of binomial distribution corresponding to expected sampling variation of bootstrap replicates in case of perfect match of markers (in all cases when selected BUSCOs were inferred as members of the same linkage group).

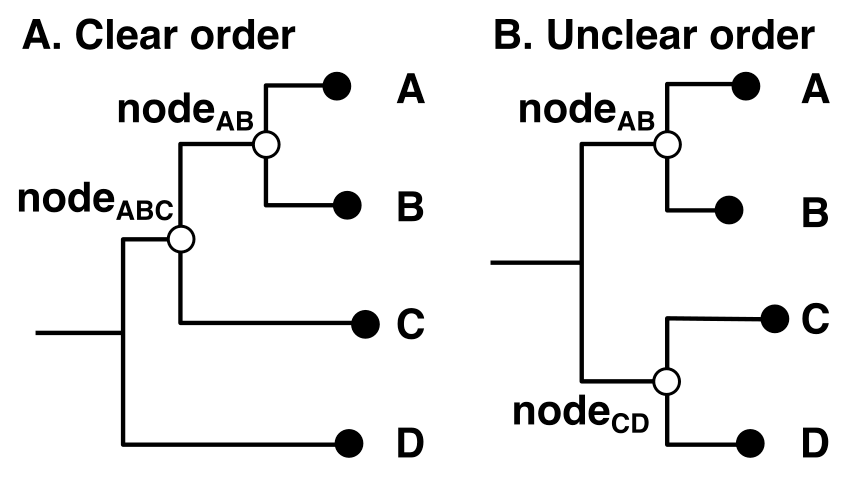

#### Fig. S3: An example phylogenetic trees with and without clear triplets for ancestral linkage group reconstruction

The tips (filled circles) represent four species with known genomes (gene markers assigned to chromosomes), the white circles represent the ancestral states to be reconstructed. **A.** The first node to be reconstructed is node_AB_ as a state that minimizes the syntenic distance of A, B and C to this node. The second reconstructed node will be node_ABC_ minimising distance to node_AB_, C and D. **B.** Representing a topology without clear resolution as node_AB_ would optimally use node_CD_ as an outgroup, and vice versa. In the original syngraph algorithm, the first node in the tree will get evaluated first and the tip with the shorter branch will be considered as an outgroup. Hence, in this example node_AB_ would be reconstructed first as the state that minimises syntenic distance to A, B, and D, and node_CD_ would proceed with a well defined tripled. Our adjusted version evaluates syntenic distances of all four possible triplets (A, B, C; A, B, D; C, D, A; C, D, B) and reconstructs first the node that minimises the syntenic distance. In case of two or more equal syntenic distances we use the first one.

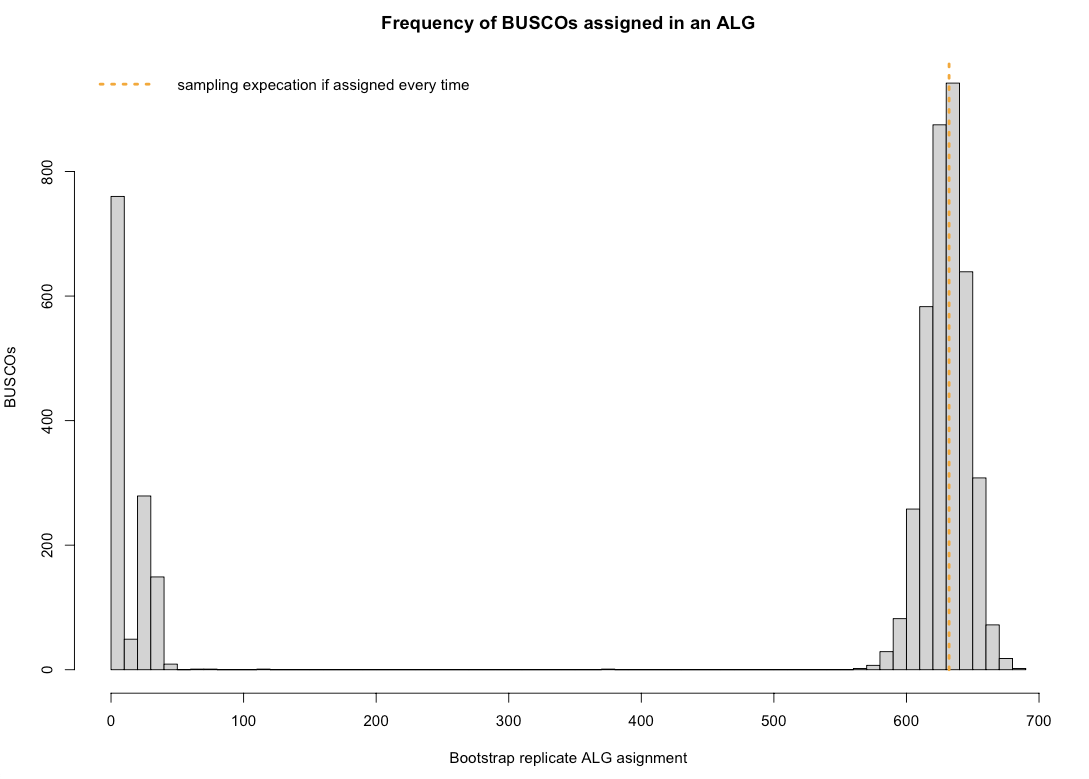

#### Fig. S4: The frequencies of BUSCOs assigned to ALGs.

Frequency of BUSCOs assigned into an ALG among 1000 bootstrap replicates. The majority of markers are within expectation given random sampling of bootstrap replicates, indicating that these markers were assigned to a linkage group in all, or nearly all, bootstrap replicates. Only genes assigned more than 327 times (1e-6 quantile of binomial distribution) were considered for assignment to ALGs.

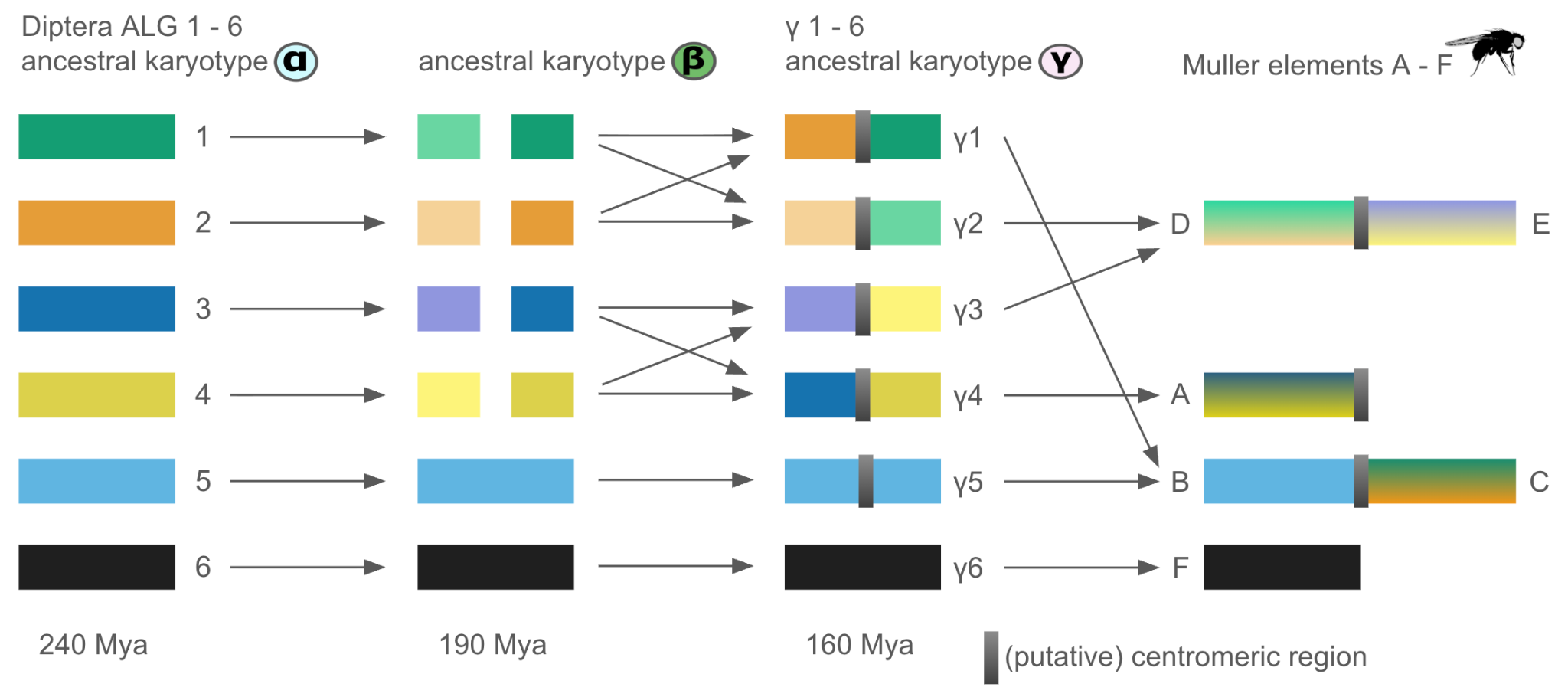

#### Fig. S5: Schematic illustration of the evolutionary transition from Diptera ALGs 1 – 6 to Muller elements A – F

We reconstructed Diptera ALGs 1–6 (ancestral karyotype α) in the last common ancestor of Diptera, which lived approximately 240 Mya. Subsequently, four of these linkage groups underwent fission in the ancestral dipteran lineage, giving rise to ancestral karyotype β, estimated to have existed 190 Mya. The resulting eight segments later fused in distinct combinations to form the metacentric linkage groups γ1–γ4 of ancestral karyotype γ, approximately 160 Mya. This karyotype remained conserved across many descendant dipteran families. A notable exception is observed in the Drosophilidae, including *D. melanogaster*. Linkage groups γ1 – γ4 lost their ancestral metacentric organization in all species in our dataset except *P. okadai*. Subsequent fusions between γ1 and γ5, as well as between γ2 and γ3, generated two novel metacentric chromosomes. During these fusion events, the ancestral centromeric structures separating chromosome arms were lost and replaced by newly formed centromeres at the fusion sites. As a consequence, the genomic content of the resulting chromosome arms became extensively rearranged relative to ancestral karyotype γ. Likewise, the acrocentric chromosome corresponding to Muller element A displays a distinct organization, characterized by intermixing of markers from two Diptera ALGs, likely reflecting disruption of the ancestral metacentric architecture. For a data-based version of this scheme capturing all gene movements, see **Fig 2C**.

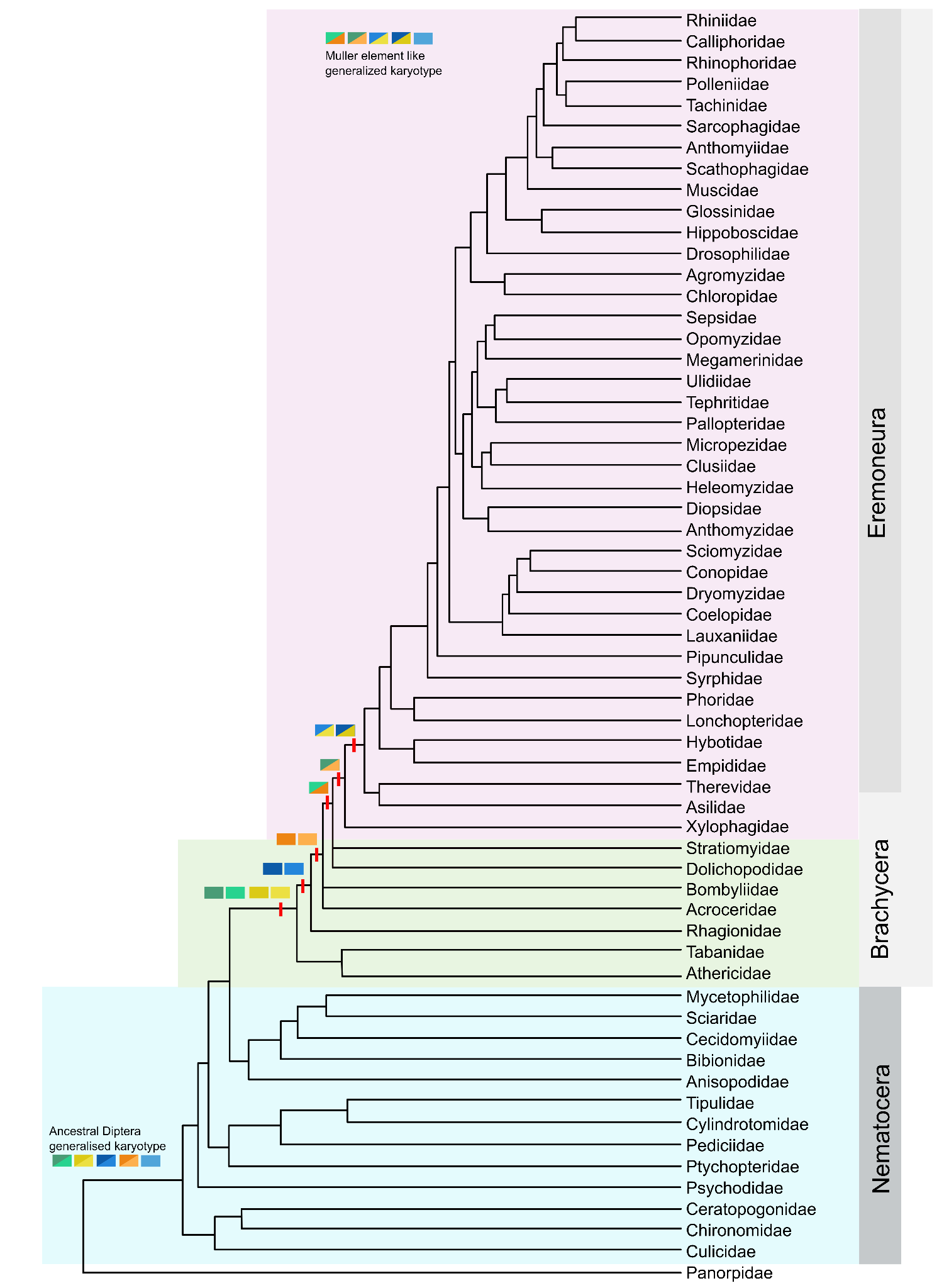

#### Fig. S6: Dipteran phylogeny reorganised to minimise the number of fissions and fusions

Specifically to minimise changes in the transition from the ancestral dipteran karyotype to that found in Eremoneua species (Muller element-like).

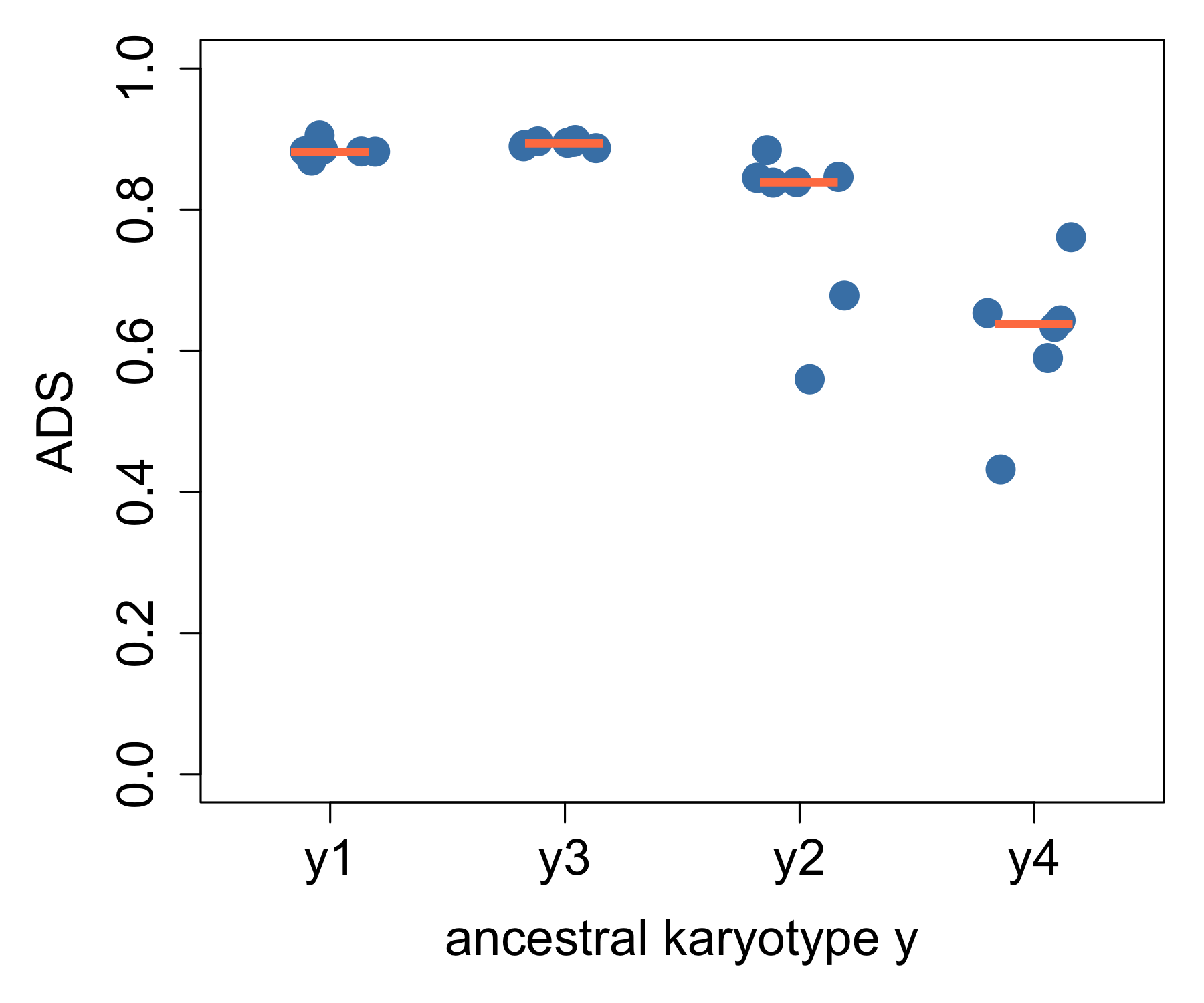

#### Fig. S7: Ancestry division scores (ADS) of chromosomes with centromere annotations.

ALGs are well separated on chromosome arms by the centromere across genomes descending from ancestral karyotype γ despite more than 75 my of divergence. Median: coral.

We have developed the ancestry division score (ADS) to quantify how effectively a centromere partitions a chromosome into co-ancestral segments, considering a chromosome composed of two ALGs, A and B.

The ADS is calculated as:

ADS = max(α_Aleft_, α_Bleft_) − min(α_Aleft_, α_Bleft_)

or equivalently

ADS = max(α_Aright_, α_Bright_) − min(α_Aright_, α_Bright_)

where α_Aleft_ denotes the proportion of markers from ALG A located on the left side of the centromere (and analogously for the other terms).

The ADS ranges from 0 to 1. A value of 1 indicates perfect separation, where each side of the centromere contains markers from only one ALG. A value of 0 indicates no separation, with equal proportions of A and B markers on both sides of the centromere.

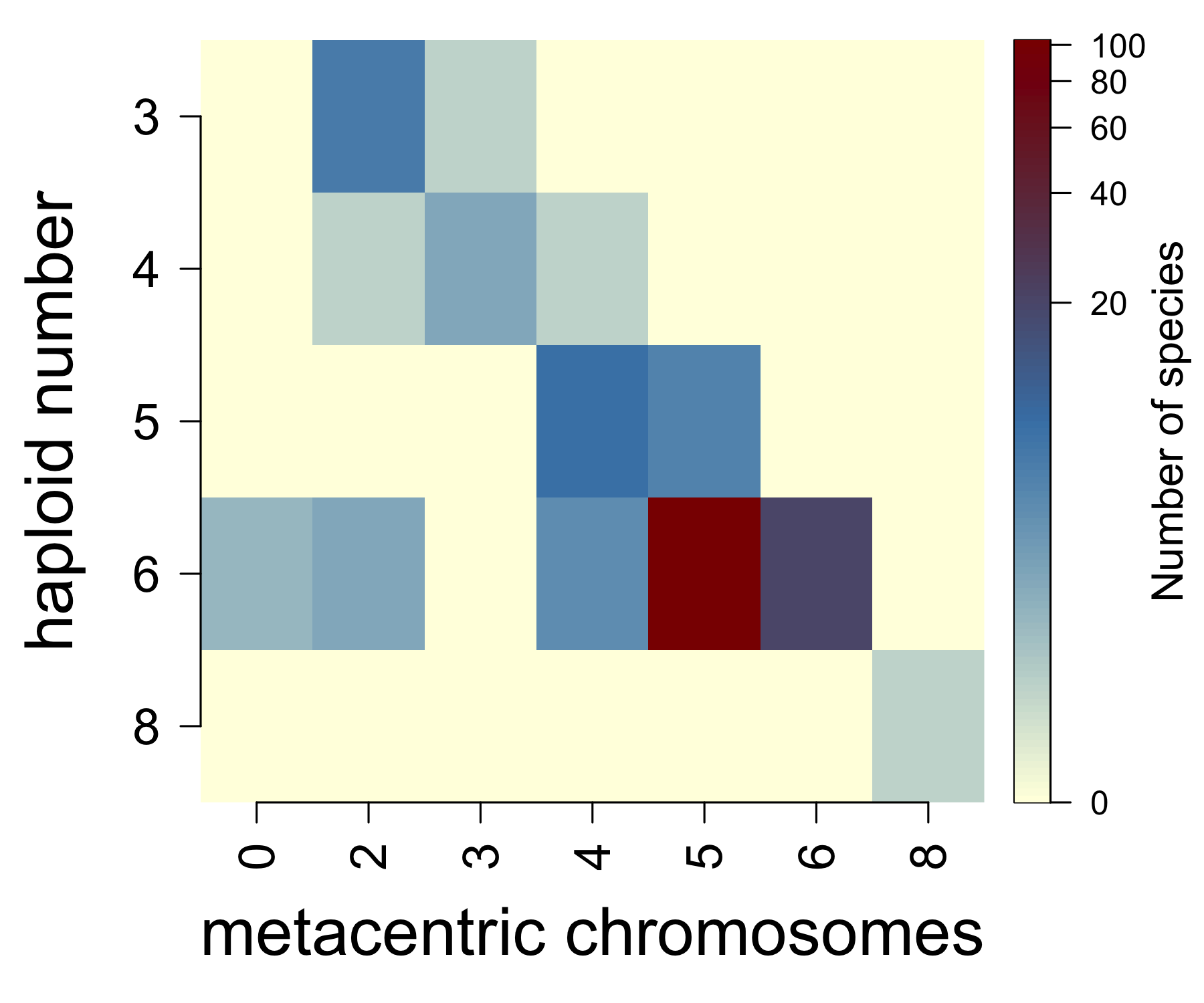

#### Fig. S8: Heat map summarizing metacentric chromosome counts per genome in species descending from ancestral karyotype γ in the Diptera karyotype database

We hypothesise that ancestral karyotype γ consisted of five metacentric ALGs, and ALG 6 with an unknown centromere position. In the metacentric ALGs, the centromere likely separated chromosome arms with distinct ALG identities, reconstructed in ancestral karyotype β. This hypothesis is supported by karyotype data compiled in the Diptera karyotype database (*20*), where the majority of species with chromosome shape annotations from Eremoneura, Therevidae, and Asilidae possess six chromosomes, with at least five being metacentric. The heat map shows the relationship between haploid chromosome number and the number of metacentric chromosomes; colours represent log10-transformed counts. Most species with annotated chromosome morphology exhibit a karyotype of n = 6 with five or six metacentric chromosomes.

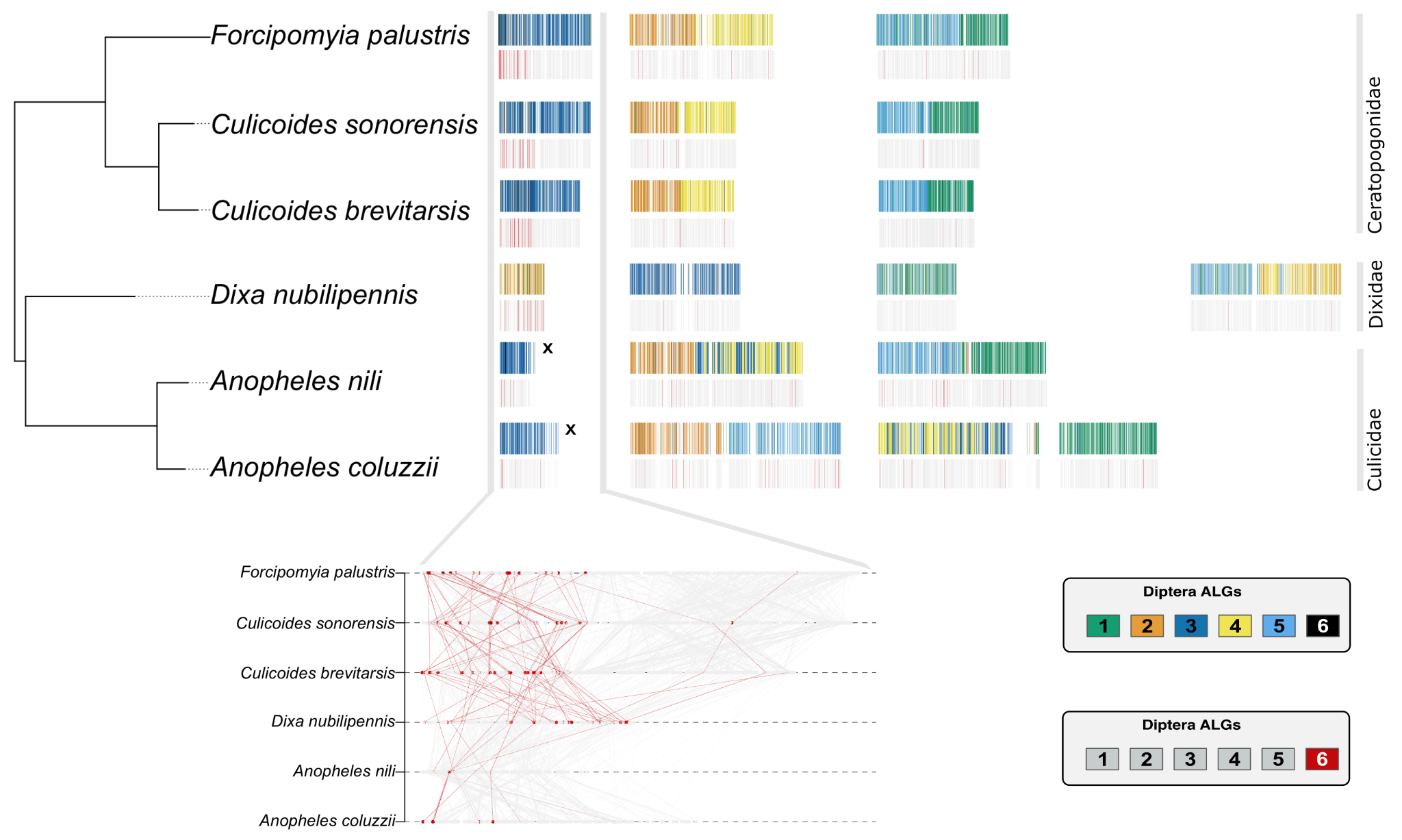

#### Fig. S9: ALG 6 likely constituted an independent linkage group in the last common ancestor of Culicomorpha

Schematic visualization of three Ceratopoginae (biting midges) genomes, one Dixidae (meniscus midges) genome and two representative Culicidae (Anophelinae) genomes. The last common ancestor of these three culicumorphan families lived 220 Mya (*22*). BUSCO genes are coloured according to their assignment to Diptera ALGs. The bottom rows highlight ALG 6 BUSCOs in red. Each row represents one genome and each element corresponds to a chromosome. *C. brevitarsis* was the only member of Ceratopogonidae included in our dataset. Since our analysis, two additional Ceratopogonidae genomes (*F. palustris* and *C. sonorensis*) as well as one Dixidae genome (*D. nubilipennis*) have become available. Ceratopogonidae and Dixidae genomes lack sex annotation of sex chromosomes. Notably, ~60% of ALG 6 BUSCOs cluster on one side of a chromosome in Ceratopogonidae, which is homologous to the X chromosome in Anophelinae. In contrast, ALG 6 BUSCOs in Anophelinae are dispersed across all three chromosomes. This pattern suggests that the dispersal of ALG 6 BUSCOs in Anophelinae represents a derived state. Additional evidence comes from the Dixidae genome, in which the majority of ALG 6 BUSCOs are concentrated on a chromosome that is distinct from the Anopheles X chromosome, suggesting an independent fusion event involving ALG 6 and an autosome in this lineage. Together, these findings indicate that ALG 6 may have existed as an independent linkage group in the last common ancestor of Culicomorpha, later undergoing lineage-specific chromosomal fusions followed by progressive gene dispersal. Overall, the results support the hypothesis that ALG 6 was ancestrally an independent, sex-linked linkage group in the last common ancestor of Diptera.

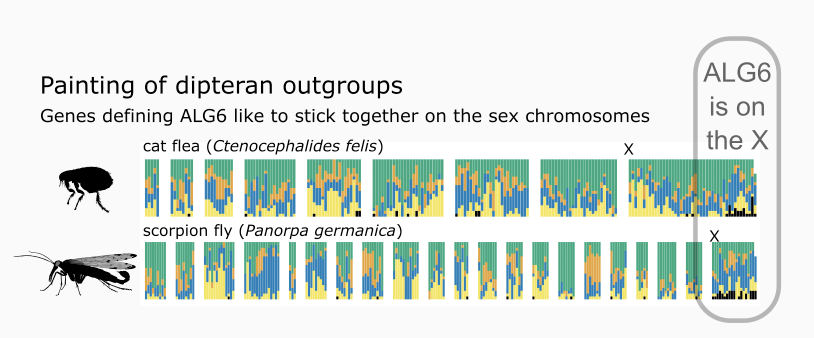

#### Fig. S10: Painting of dipteran outgroups: cat flea and a scorpion fly

Genomes of a scorpion fly *Panorpa germatica* (*44*) and a cat flea *Ctenocephalides felis* (*45*) painted by the ALGs of Diptera using colours consistent with the rest of the manuscript. The sections divided by the white gaps represent individual chromosomes and each bar represents 20 consecutive BUSCO genes painted by their origin (larger section therefore indicates more BUSCO genes on the chromosome). The outgroups show no conserved synteny with Diptera for all ancestral linkage groups except for ALG 6. Genes associated with ALG 6 are located in both cases on the sex chromosomes of the species. In the case of *P. germanica*, the genes are dispersed across the whole sex chromosome. In the case of the cat flea, the ALG 6 genes are localised at a side of the sex chromosome.

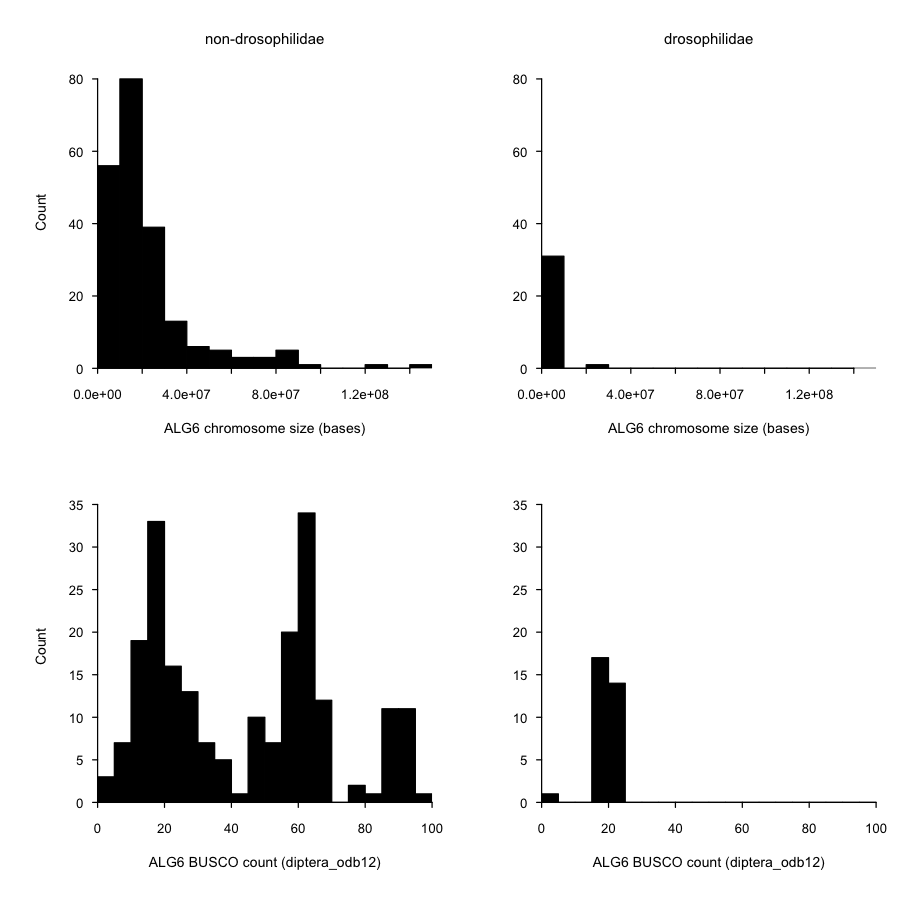

#### Fig. S11: Sizes and numbers of BUSCO of the ALG 6 chromosomes in Drosophilidae and other Diptera

Drosophilidae show much smaller ALG 6 chromosomes with substantially smaller number of BUSCO genes (typically 20-22 compared to 49-51 in other Diptera).

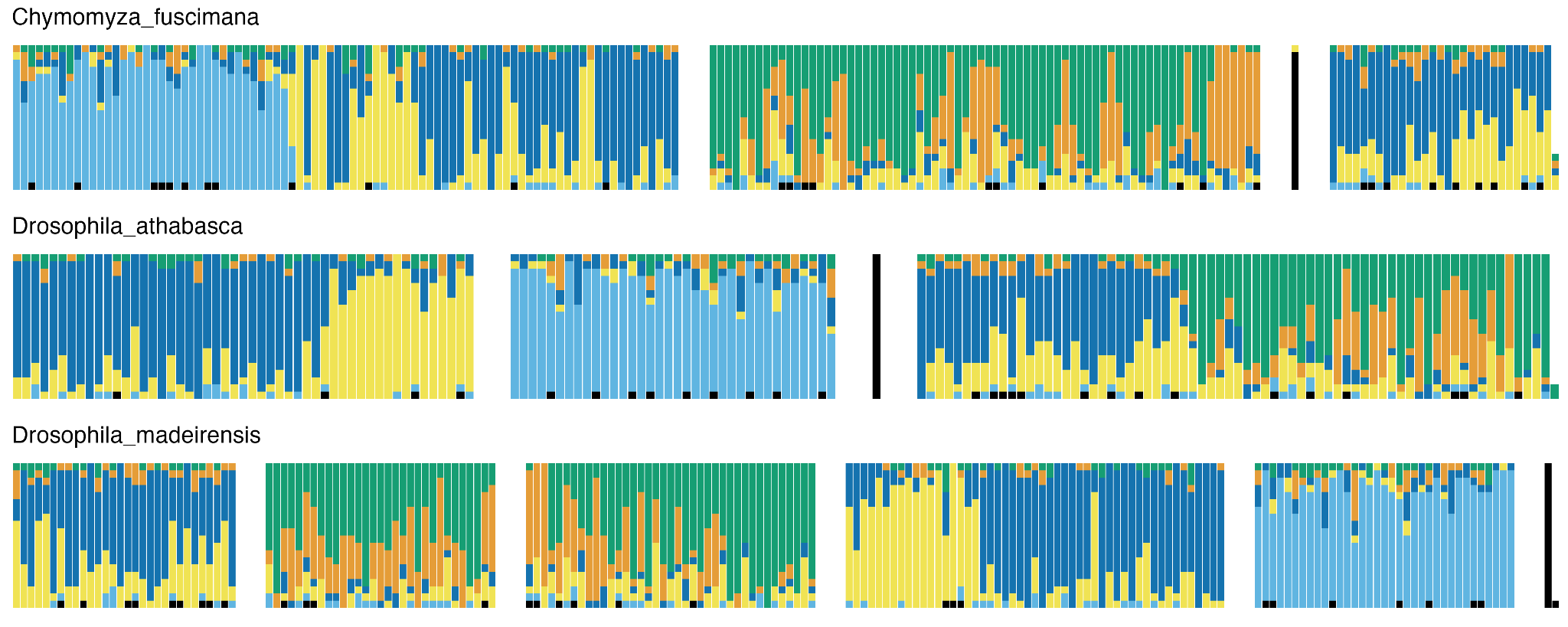

#### Fig. S12: Painting of Drosophilidae species show that the majority of ALG 6 derived BUSCOs lost from Muller element F and dispersed across the genome

Genomes of three Drosophilidae species painted by the ancestral linkage group of diptera using colours consistent with the rest of the manuscript. The sections divided by the white gaps represent individual chromosomes and each bar represents 20 consecutive genes painted by their origin. In all three species there is ‘the dot’ chromosome containing nearly exclusively ALG 6 genes. The reminder of the genes is dispersed across their genomes.

### Supplementary Tables

#### Table S1. Table of 346 genomes used in the study with associated metadata.

This table includes citations (DOI) for previously published genomes.

**Table S2: Number of BUSCO markers from diptera_odb12 per chromosome per genome.**

The alg6_derived_chromosome column marks chromosomes that have >50% of markers originating from ALG 6.

**Table S3: Summary statistics for diptera_odb12 BUSCO runs per genome.**

**Table S4: Table outlining new and known sex chromosome changes across Diptera.**

The “How do we know” column denotes the type of evidence for the sex chromosome change. GENOMES - information from known sex chromosomes; chromosomal genomes; IMPLIED - genomes without known sex chromosomes that do not feature the ancestral or potentially ancestral sex chromosome; LITERATURE - any other method (sometimes even genome based) but we did not use the data, we took it from the paper as described. References are recorded in a form of DOI.

### Table S5: Sex chromosome identities confirmed using read data.

### Supplementary Data

**Data S1:** Diptera ALG ‘paintings’ for each chromosome of each genome organised by family. Paintings show ALG assignment by colour collated into bins of 20 BUSCO genes.
