## Supplementary figures and images for "340 dipteran genomes reveal the origin of Muller elements and sex chromosomes in Diptera"

### 1_Rhiniidae.png

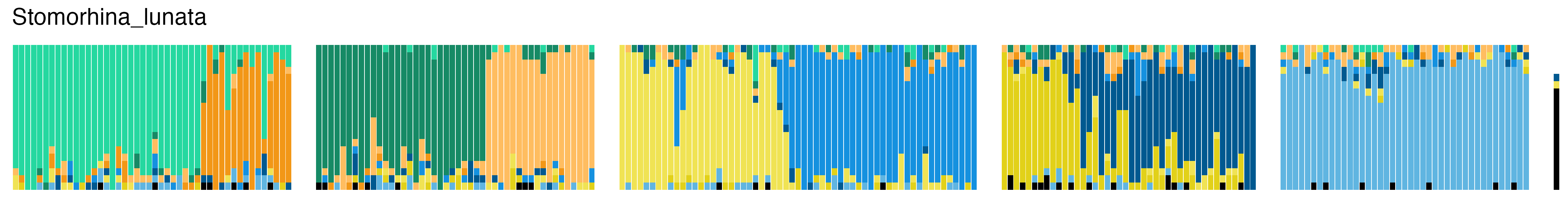

### 2_Calliphoridae.png

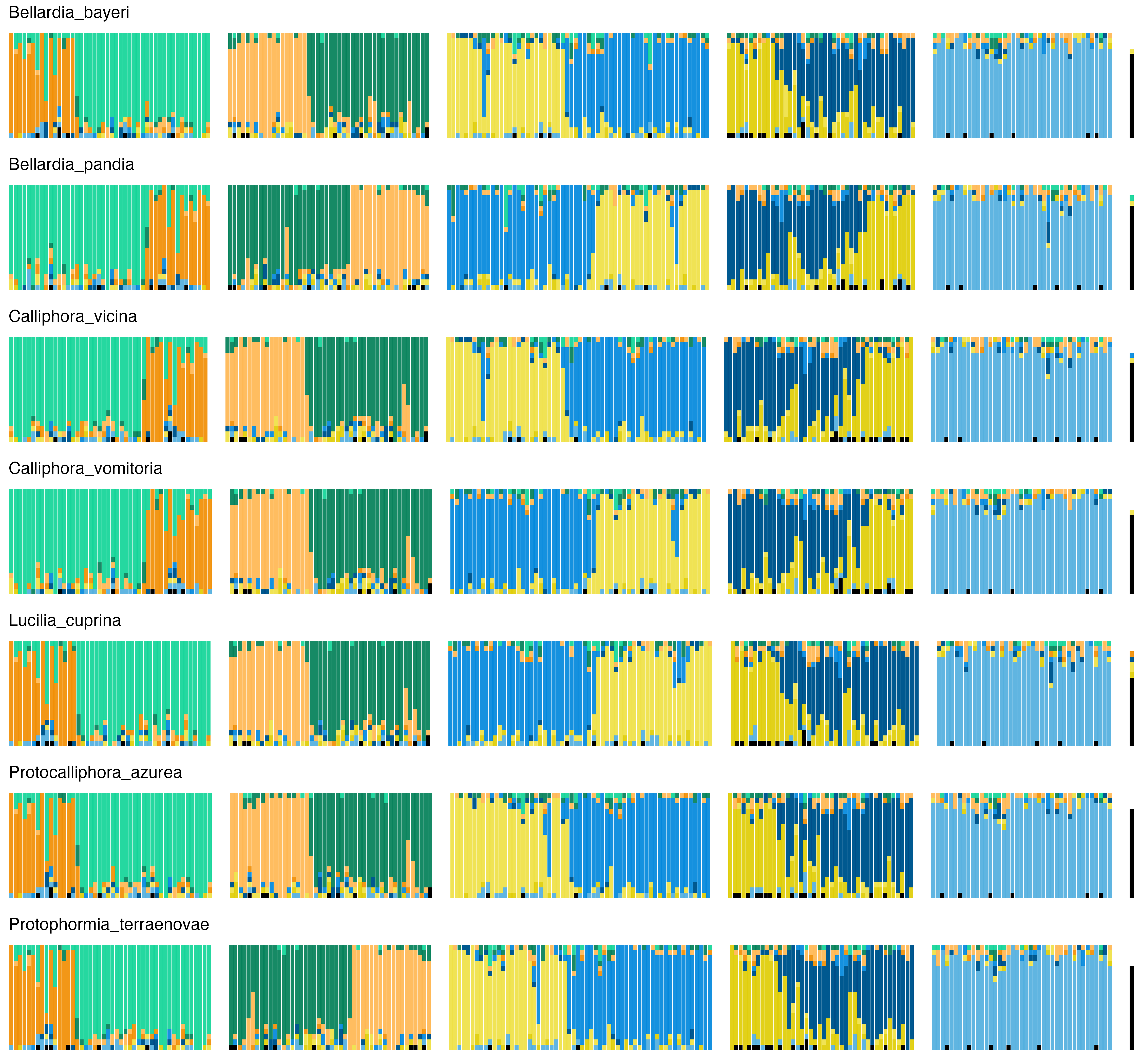

### 3_Rhinophoridae.png

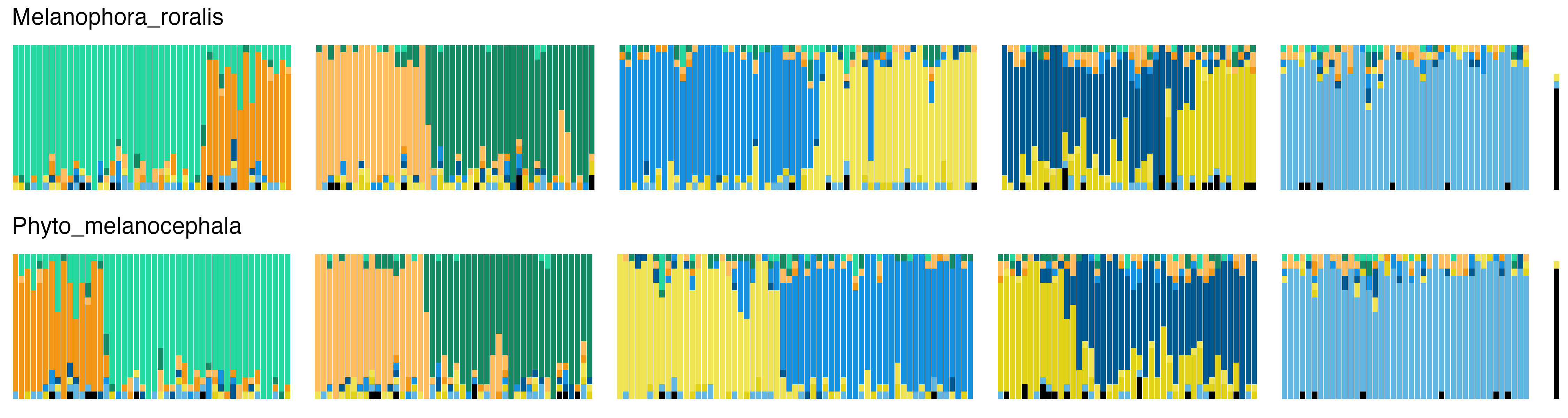

### 4_Polleniidae.png

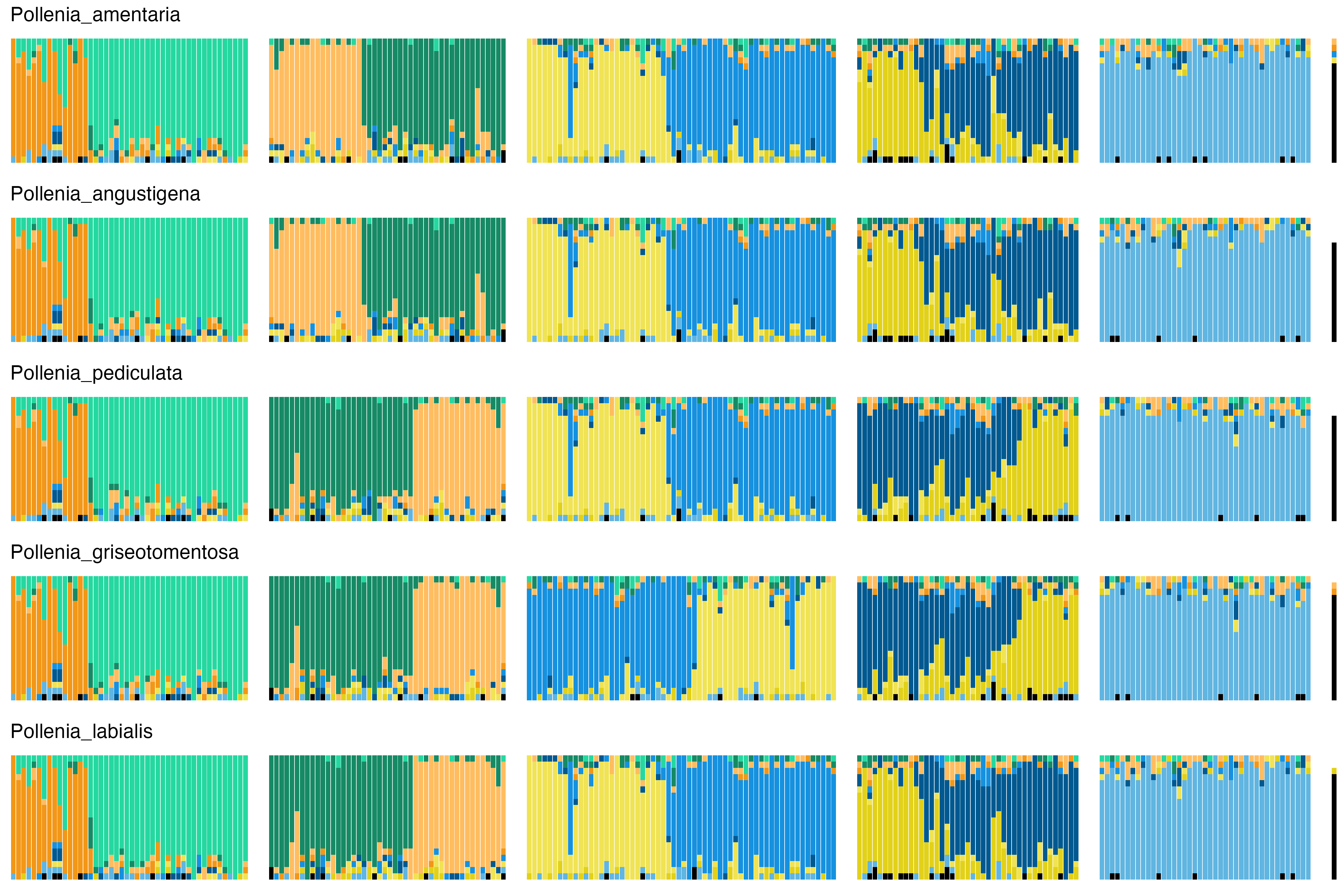

### 5_Tachinidae.png

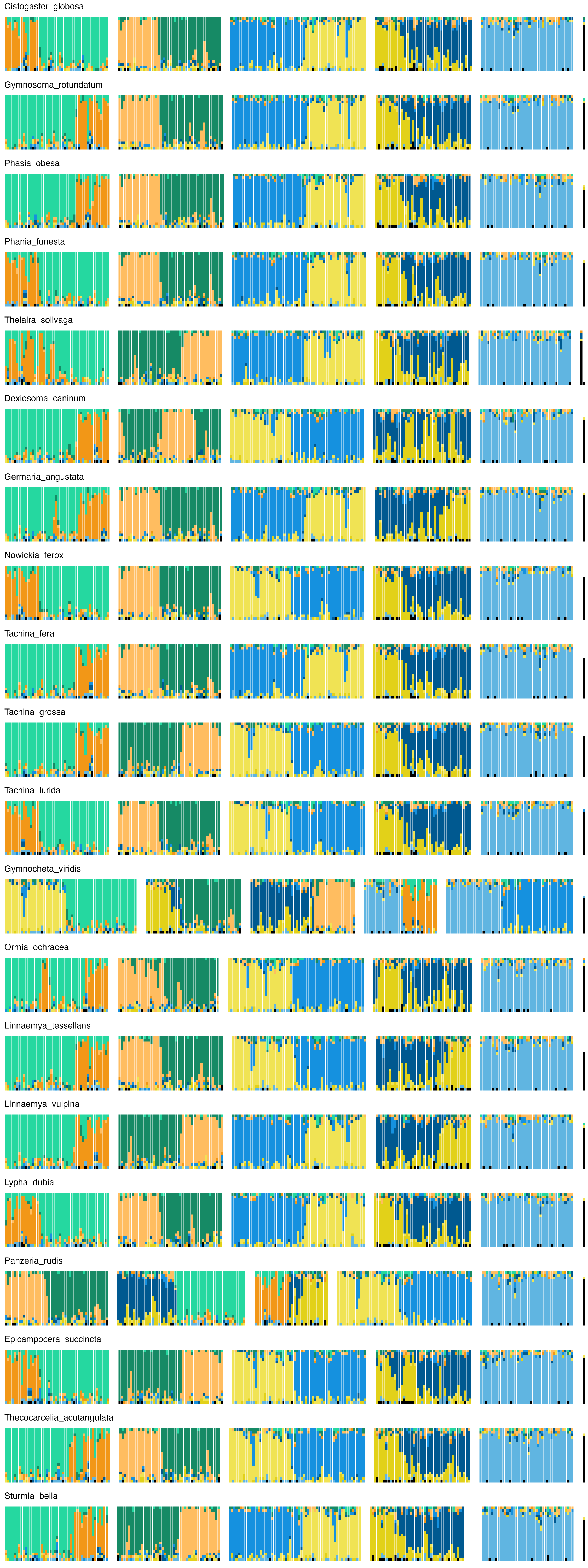

### 6_Sarcophagidae.png

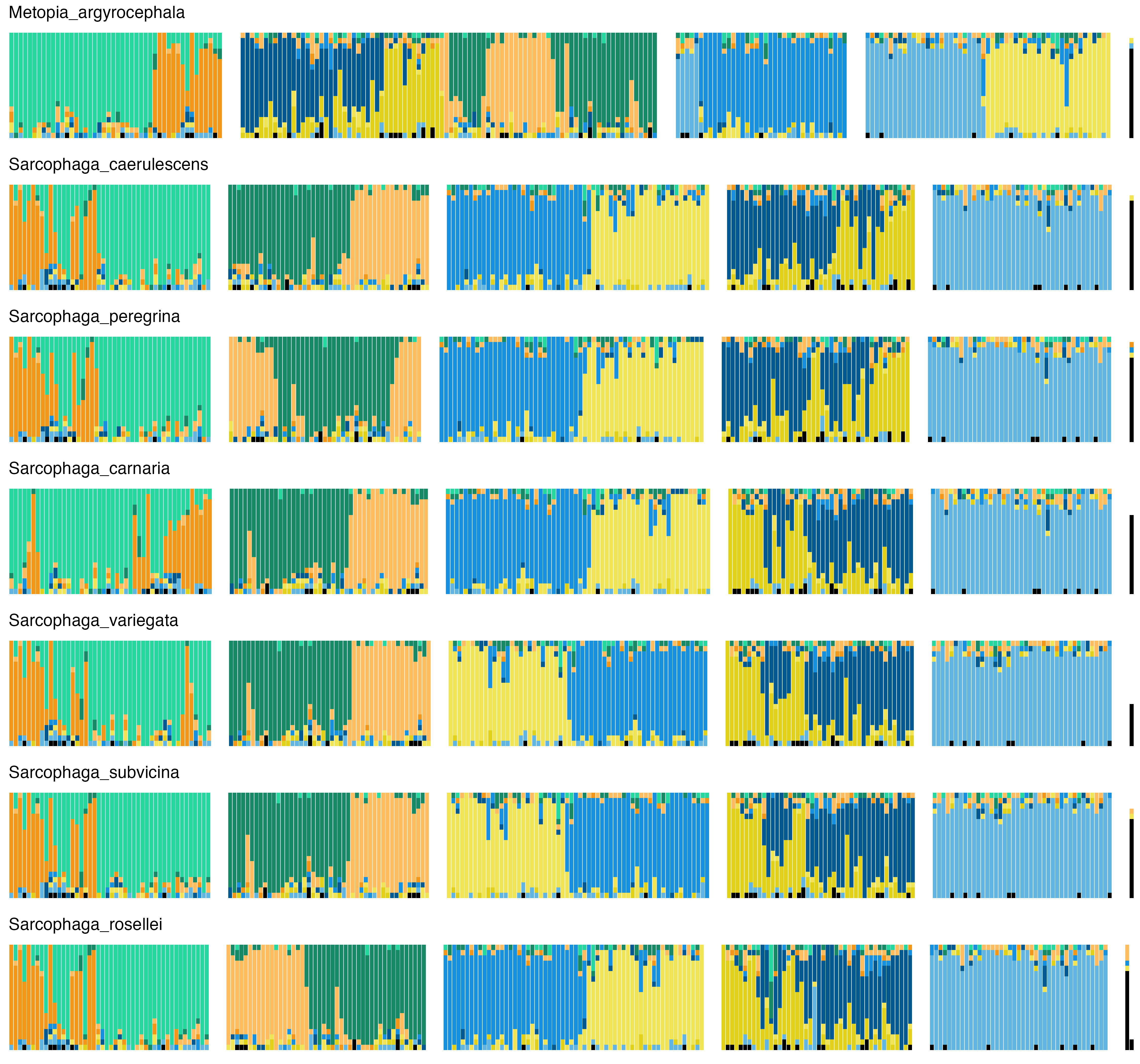

### 7_Anthomyiidae.png

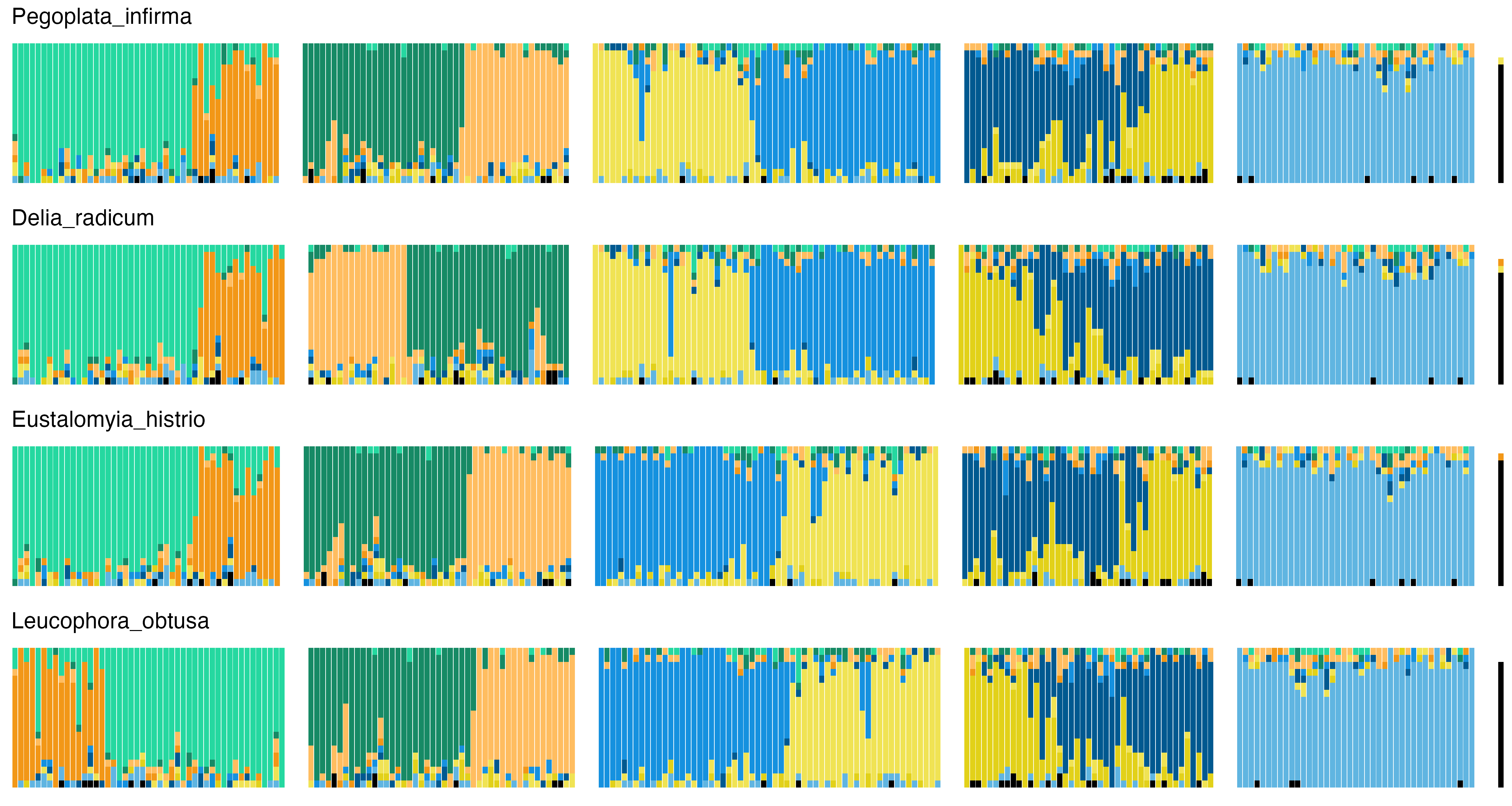

### 8_Scathophagidae.png

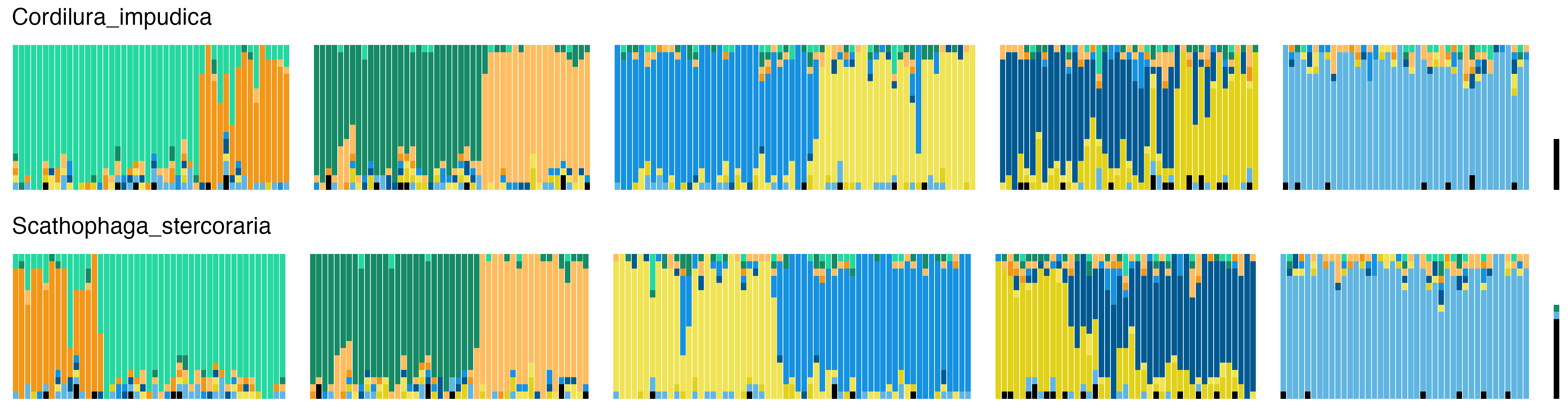

### 9_Muscidae.png

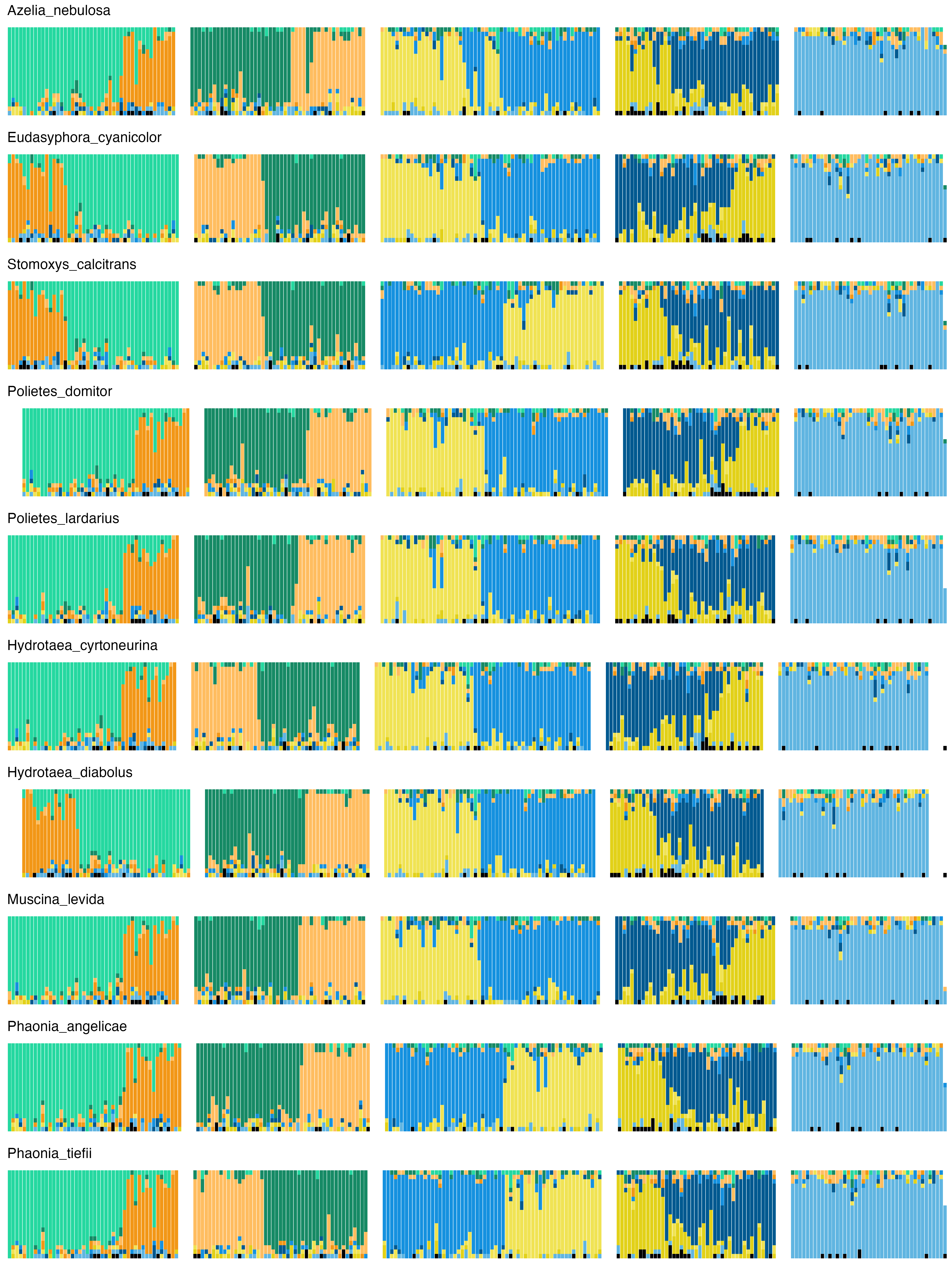

### 10_Glossinidae.png

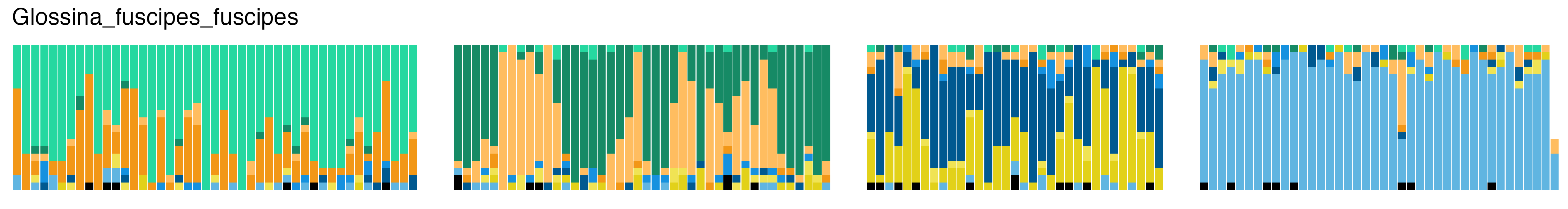

### 11_Hippoboscidae.png

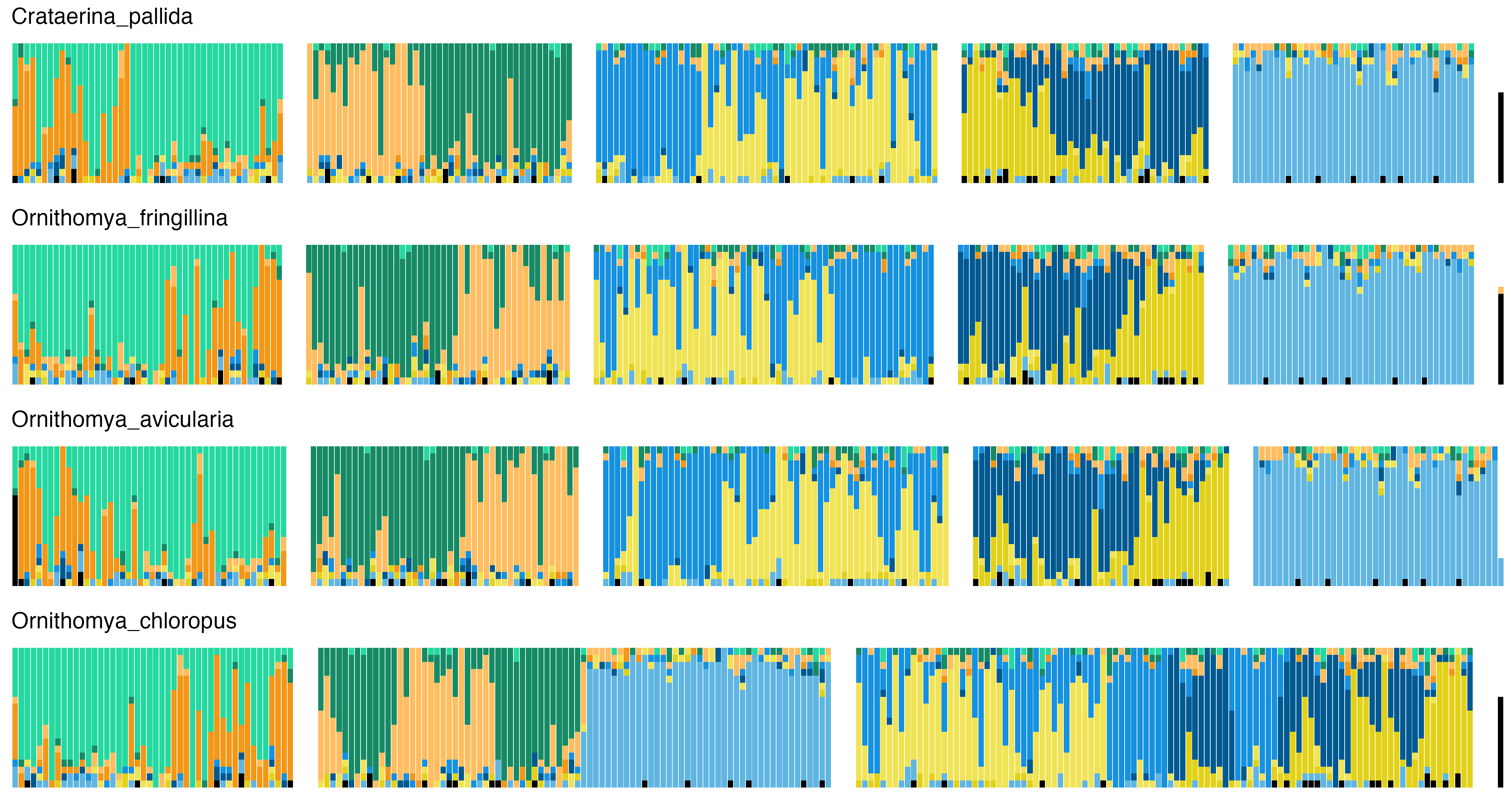

### 12_Drosophilidae.png

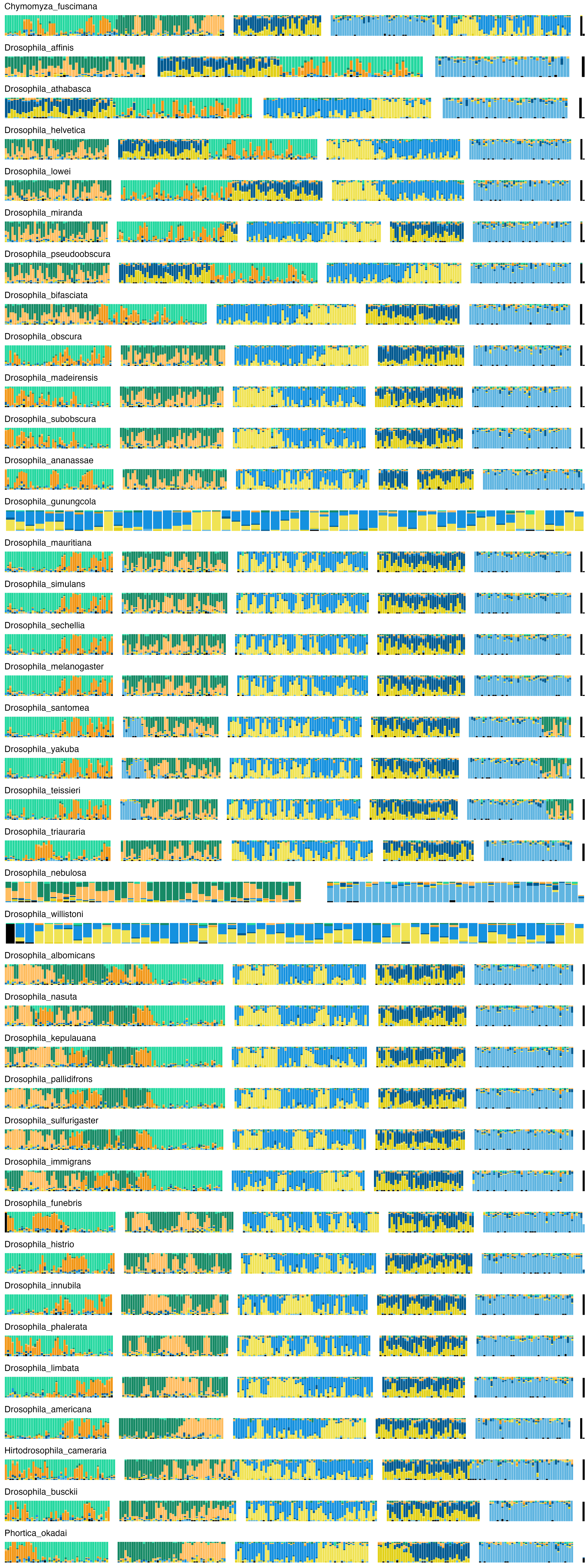

### 13_Agromyzidae.png

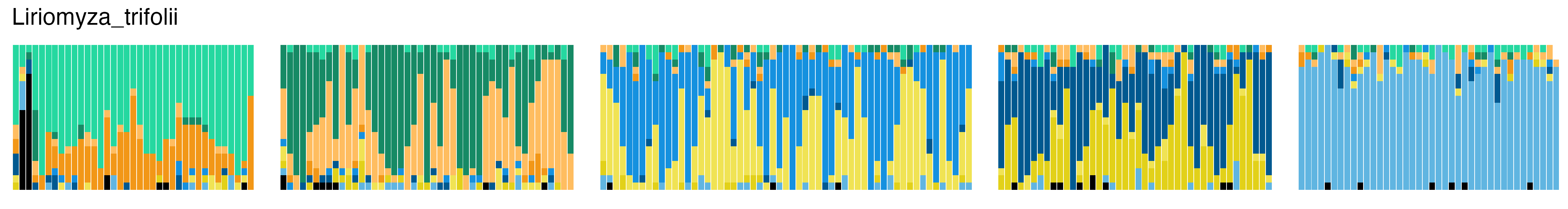

### 14_Chloropidae.png

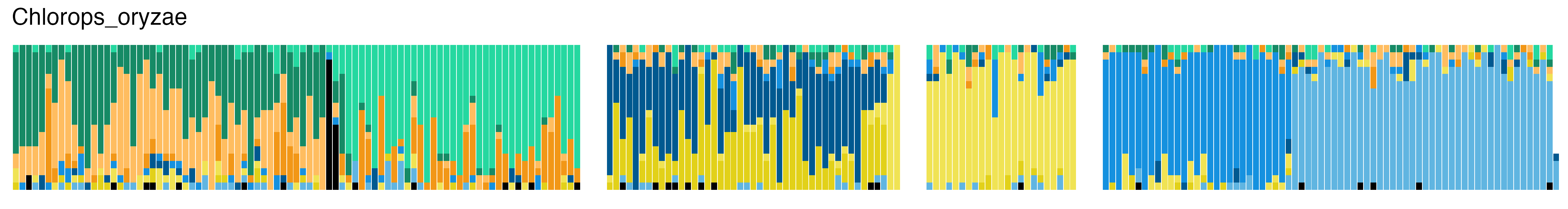

### 15_Sepsidae.png

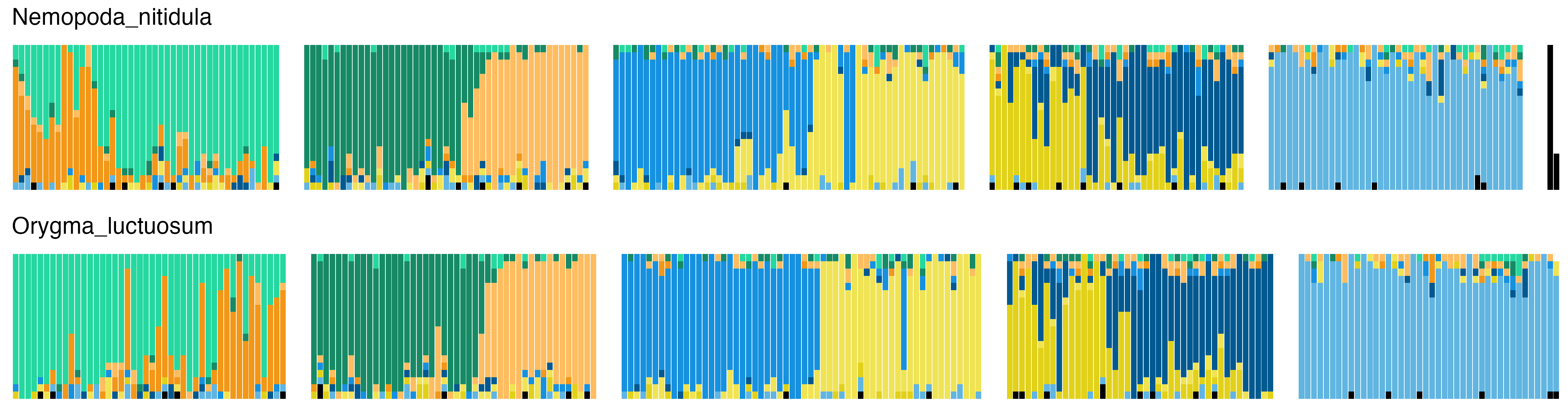

### 16_Opomyzidae.png

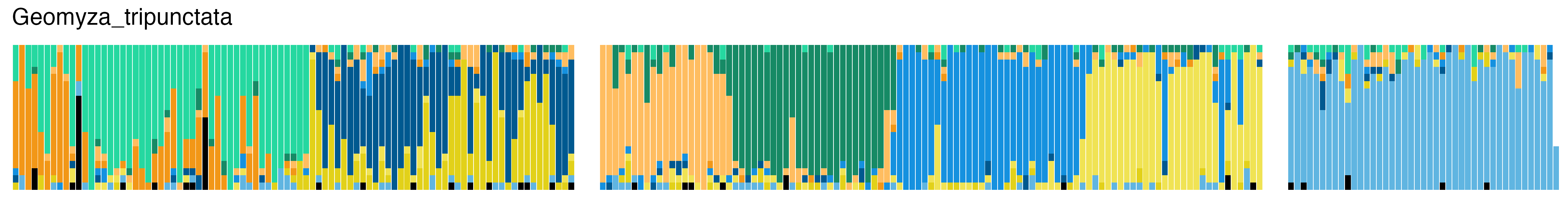

### 17_Megamerinidae.png

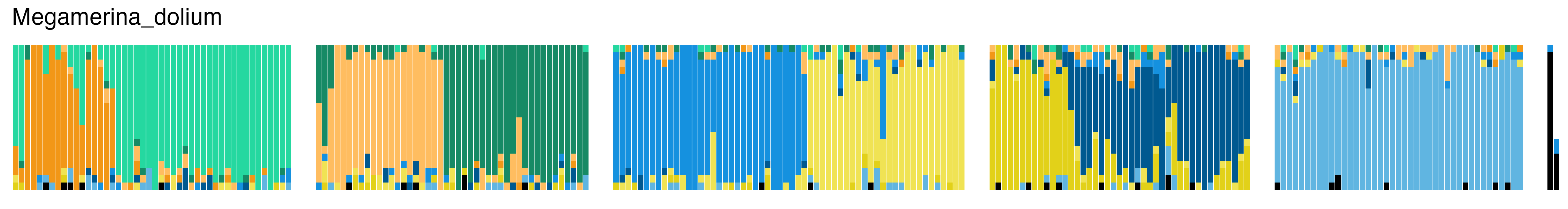

### 18_Ulidiidae.png

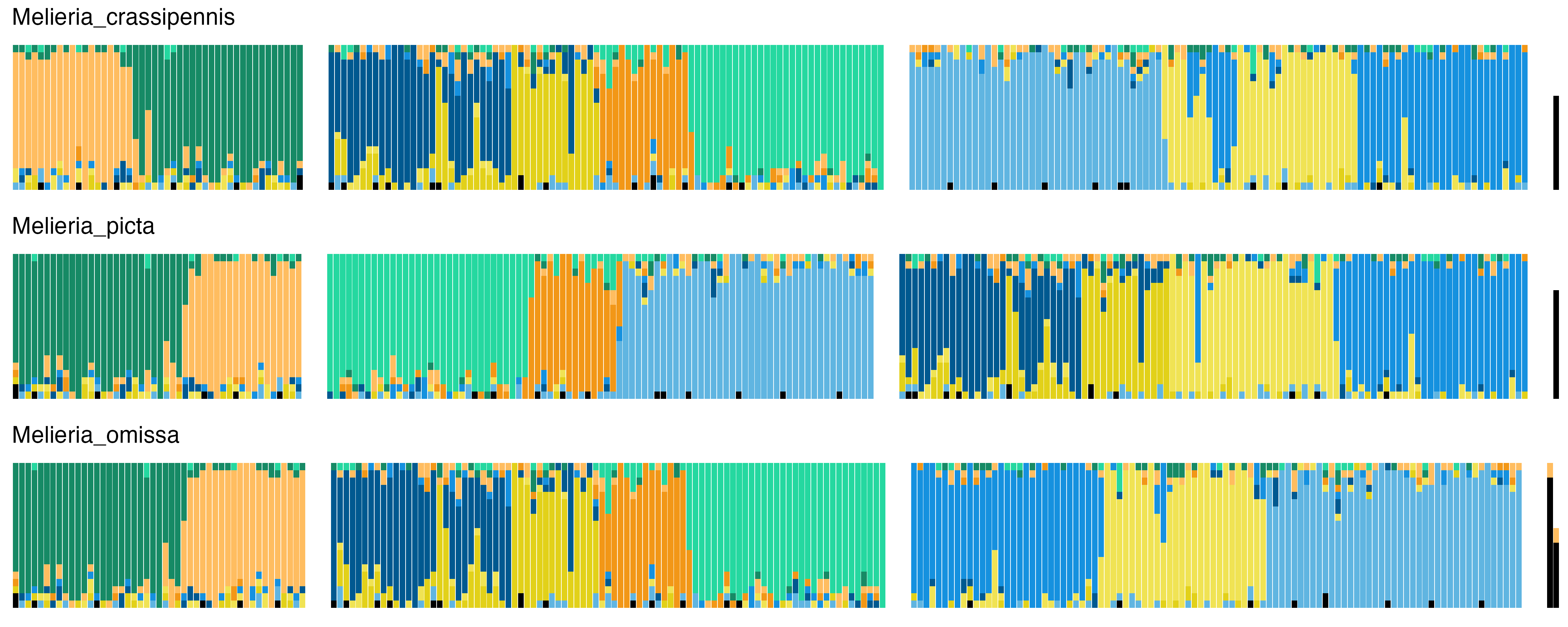

### 19_Tephritidae.png

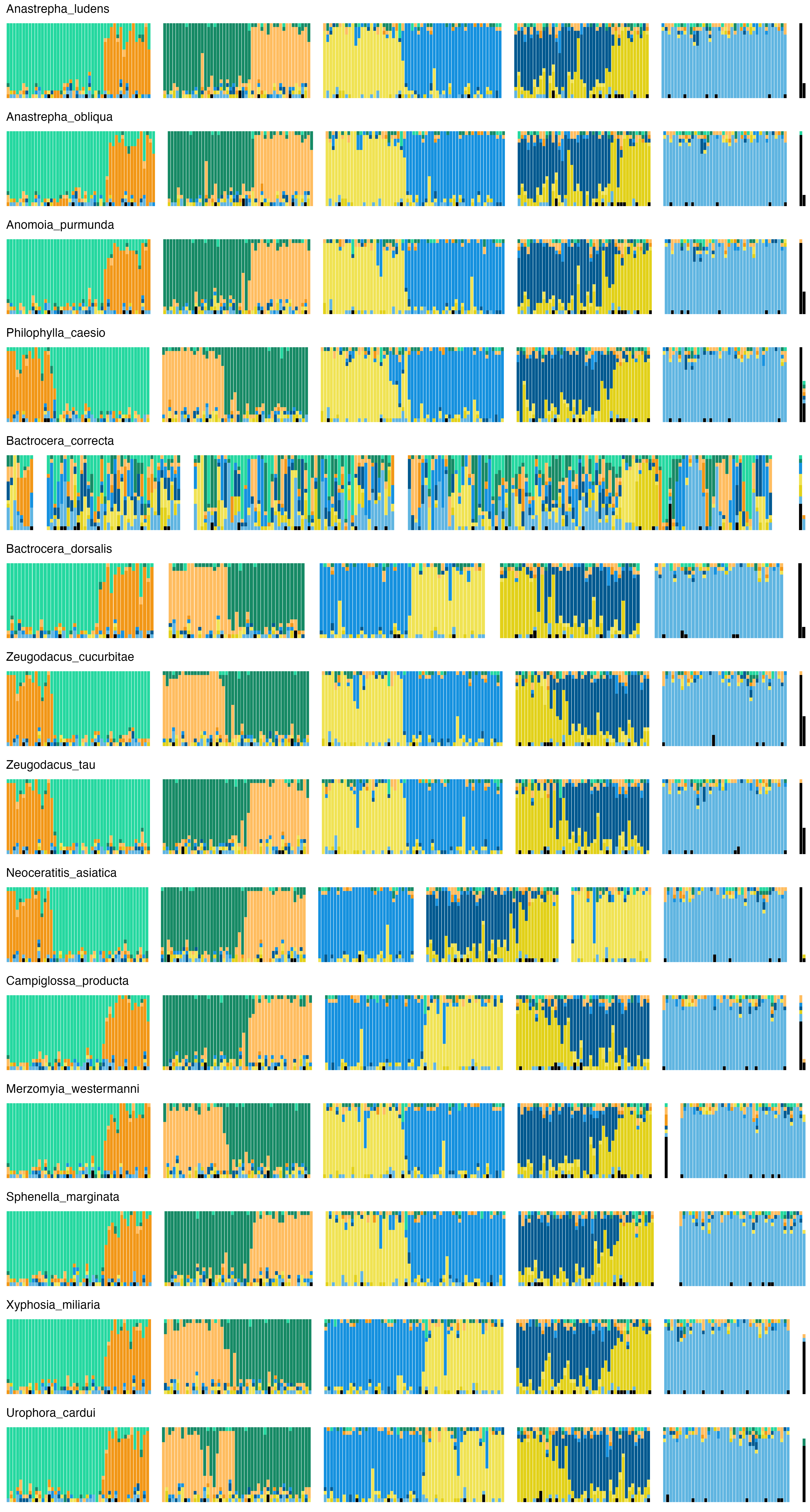

### 20_Pallopteridae.png

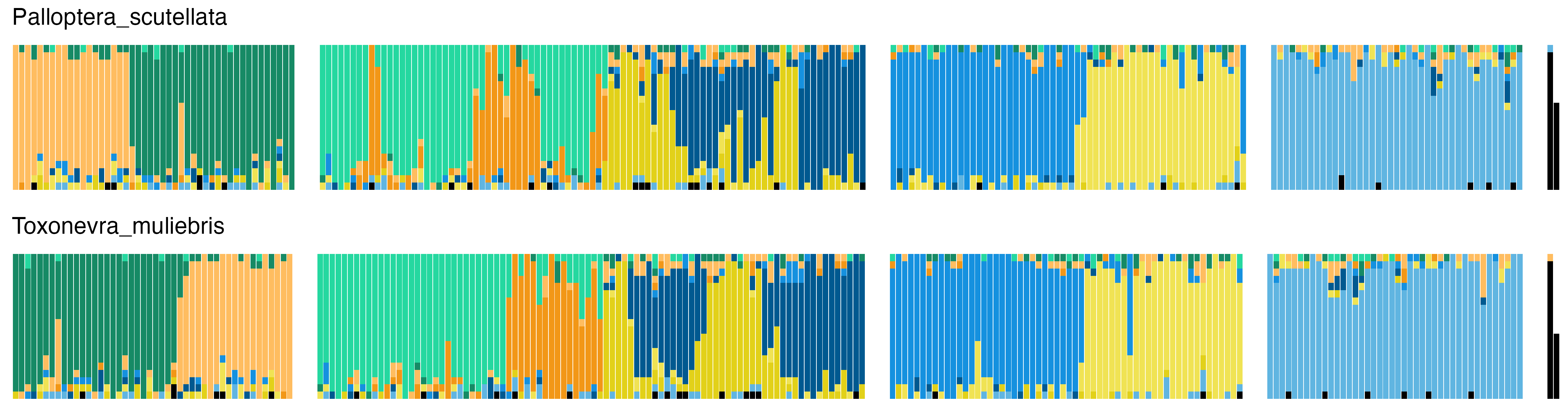

### 21_Micropezidae.png

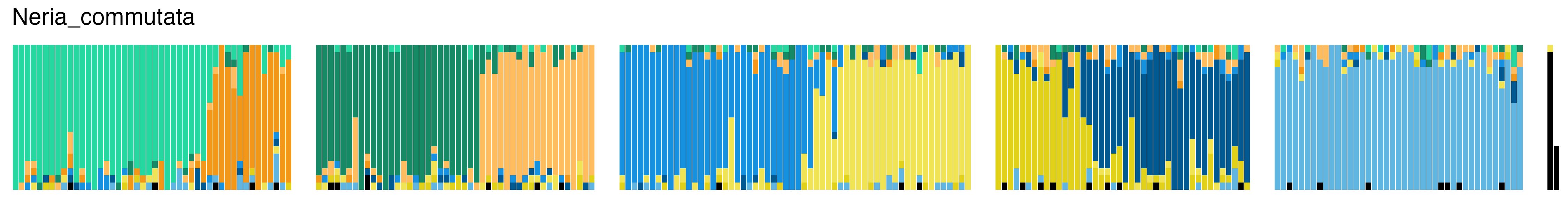

### 22_Clusiidae.png

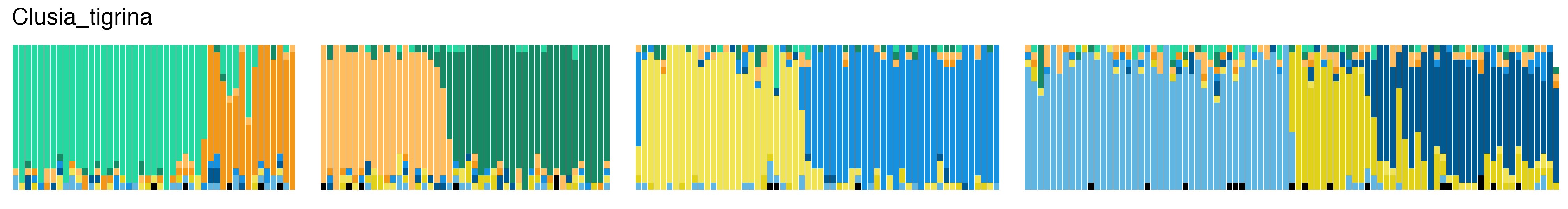

### 23_Heleomyzidae.png

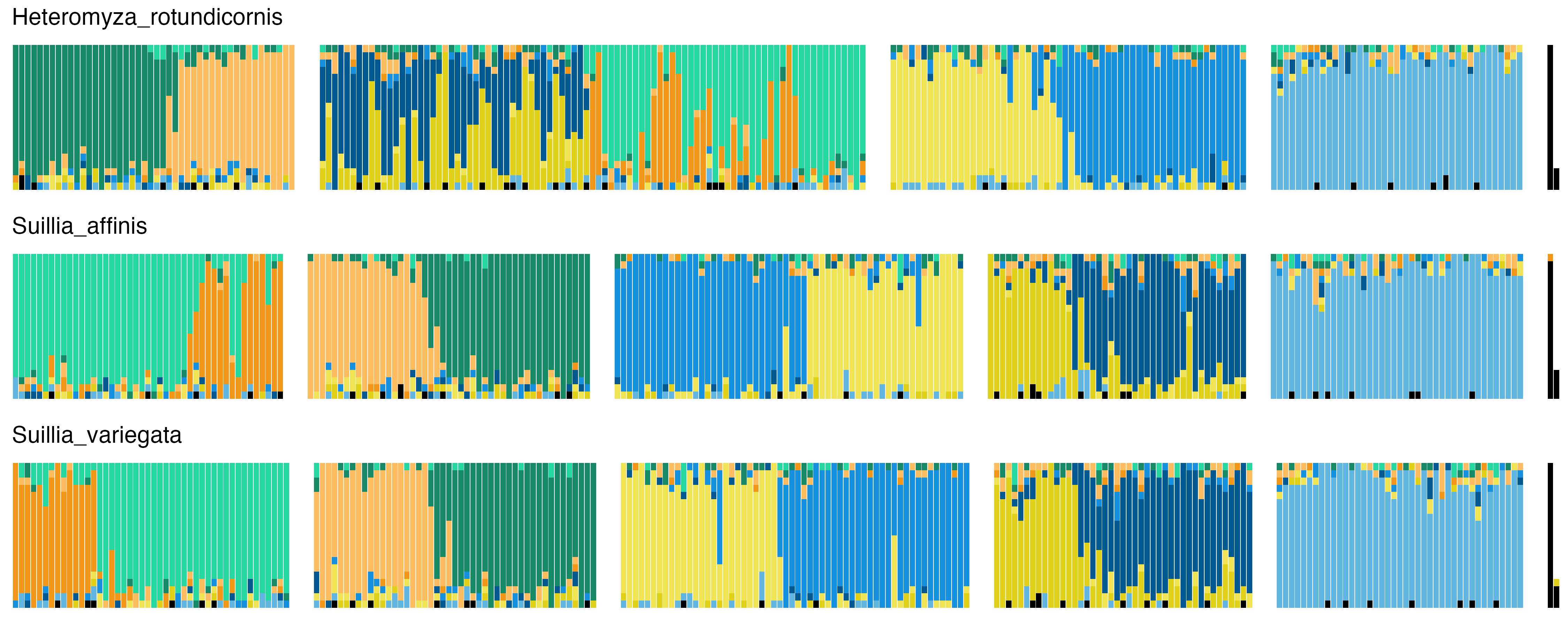

### 24_Diopsidae.png

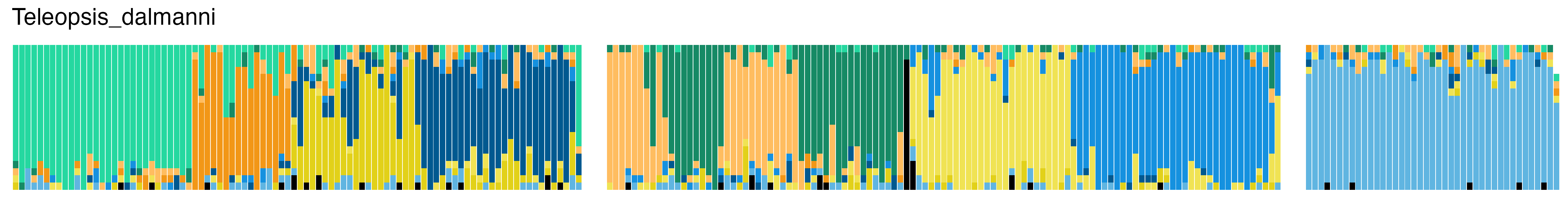

### 25_Anthomyzidae.png

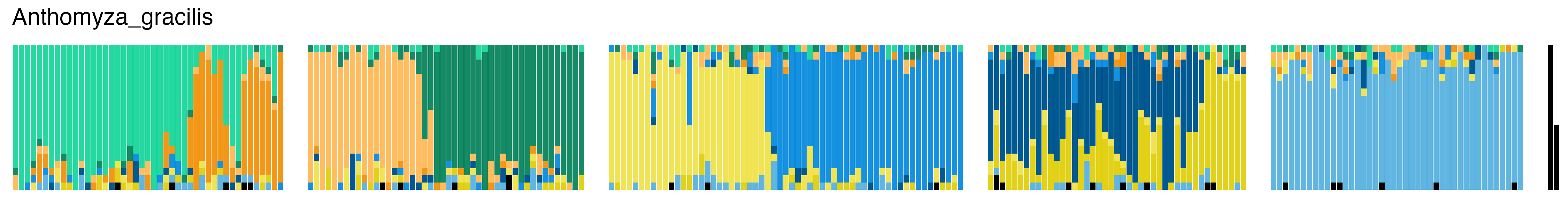

### 26_Sciomyzidae.png

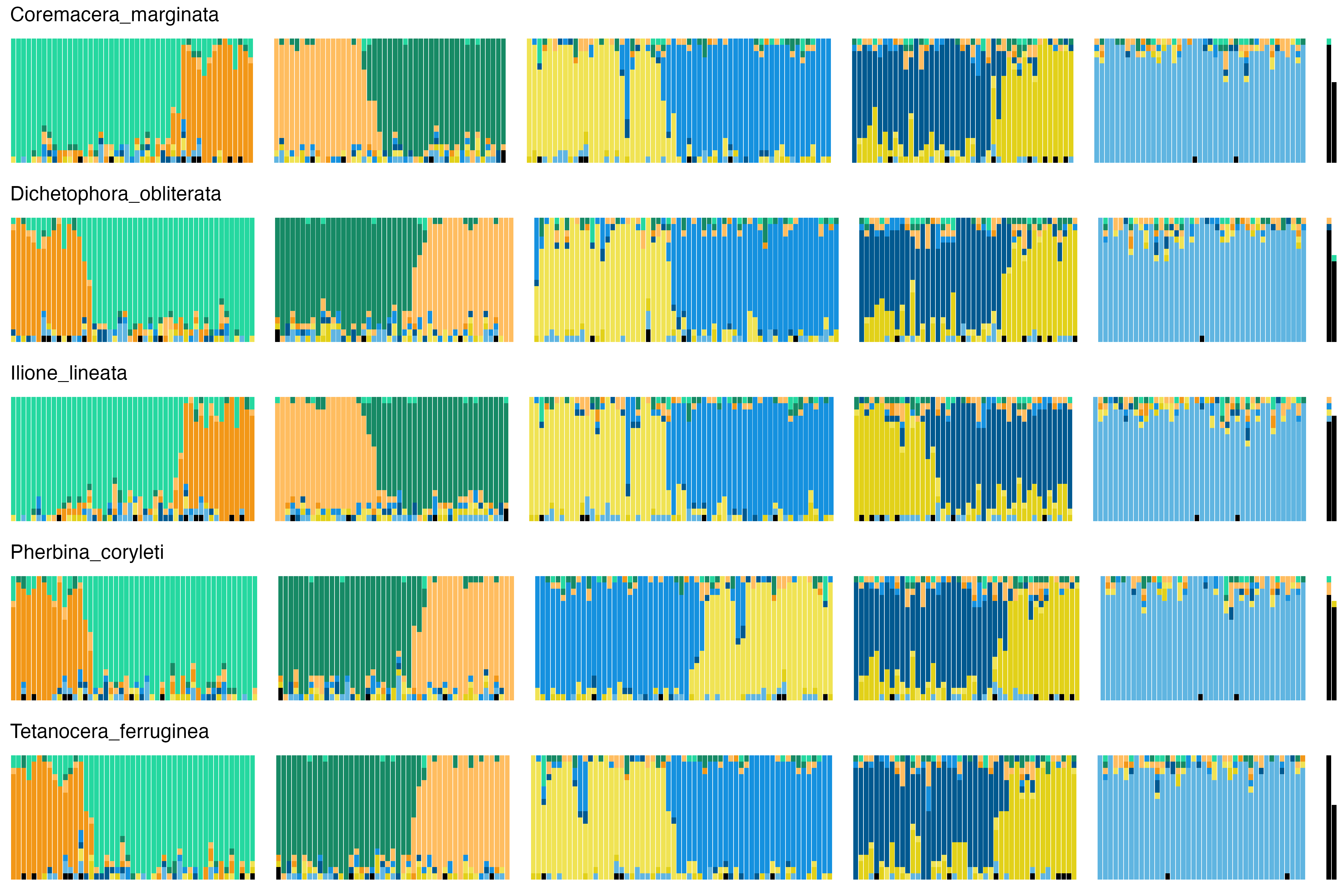

### 27_Conopidae.png

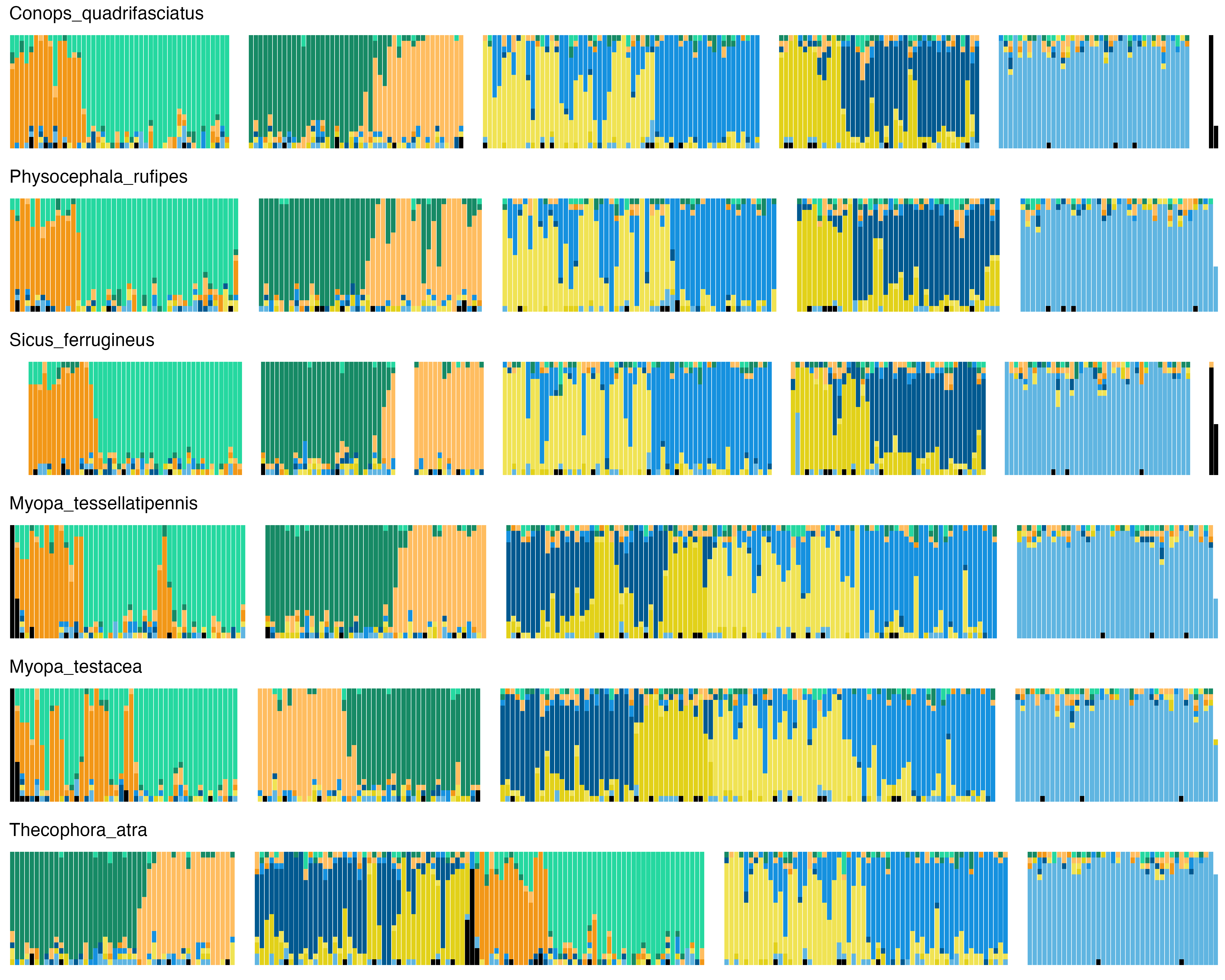

### 28_Dryomyzidae.png

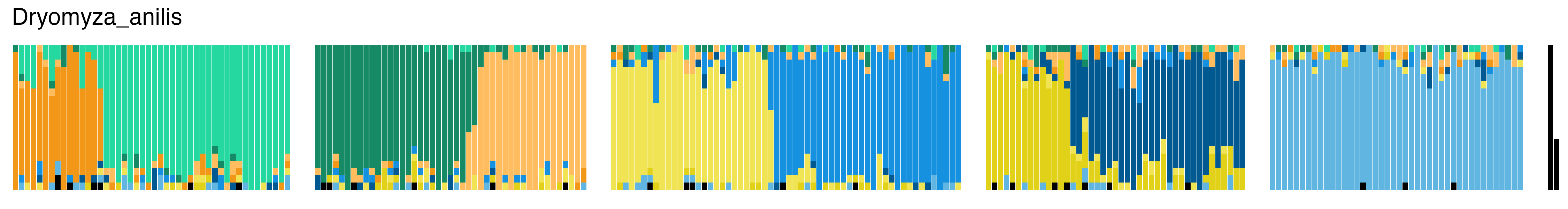

### 29_Coelopidae.png

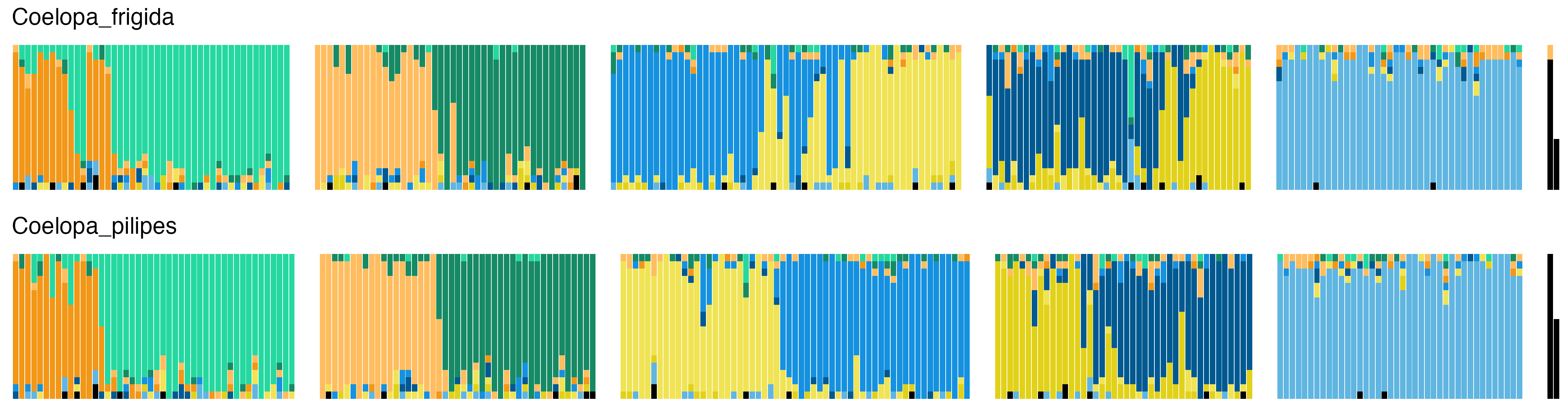

### 30_Lauxaniidae.png

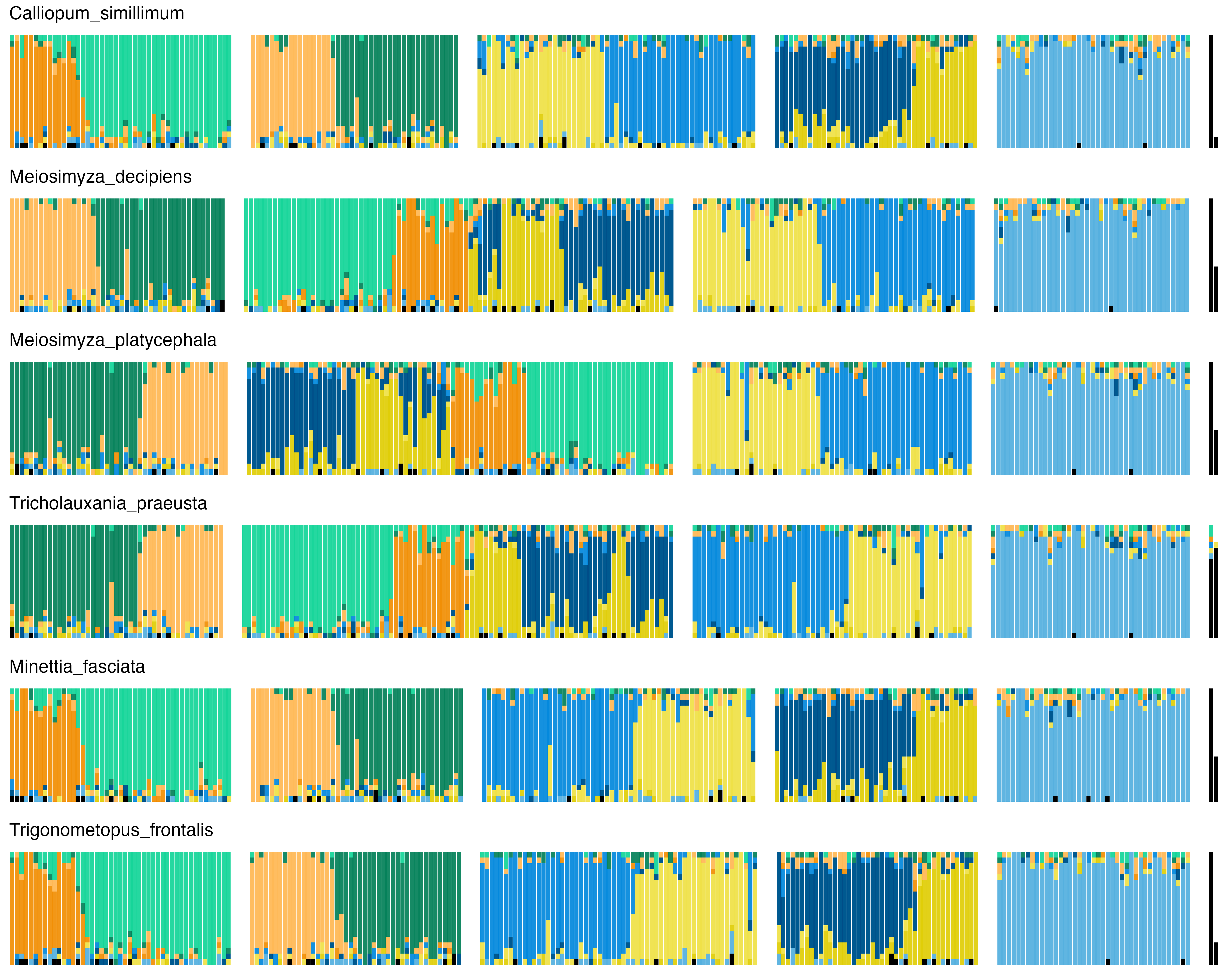

### 31_Pipunculidae.png

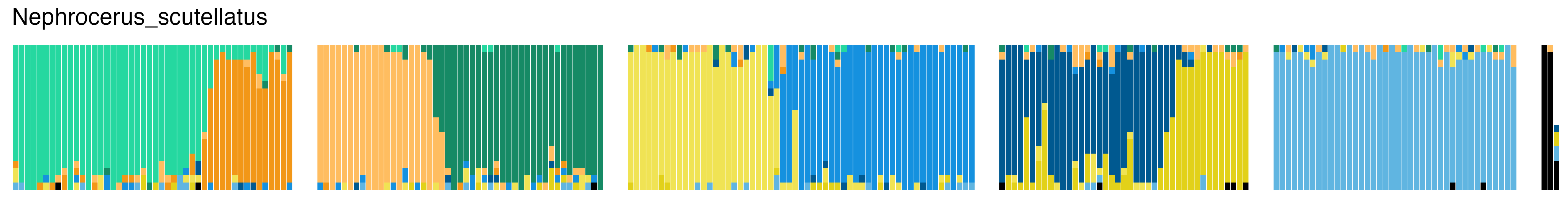

### 32_Syrphidae.png

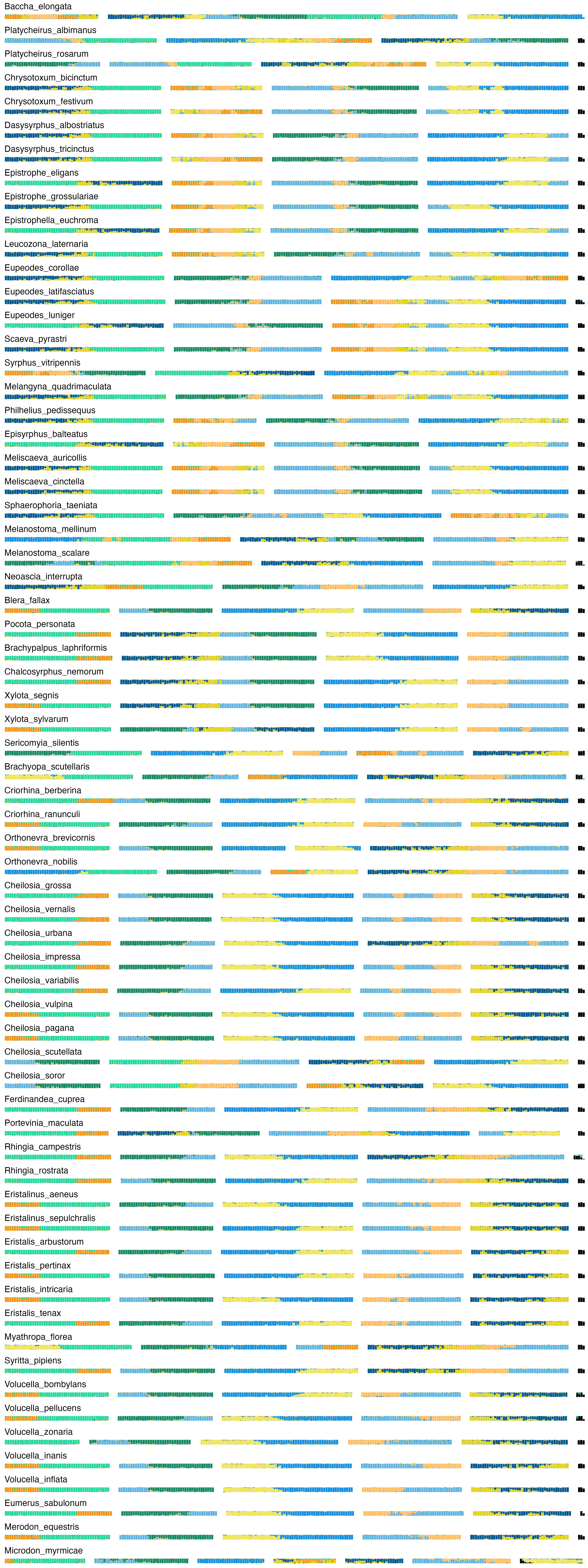

### 33_Phoridae.png

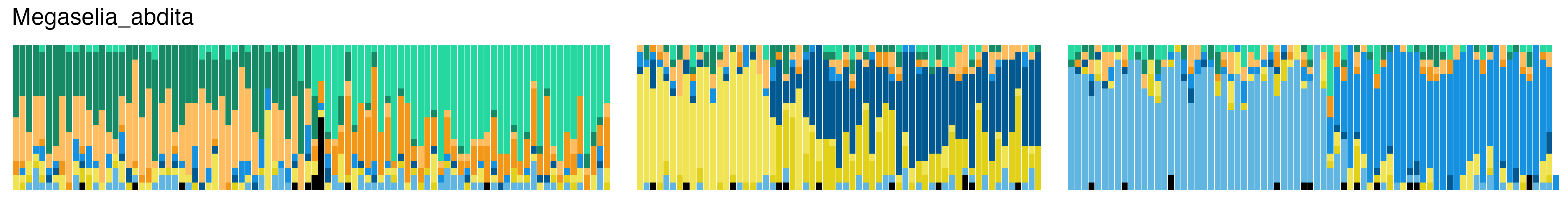

### 34_Lonchopteridae.png

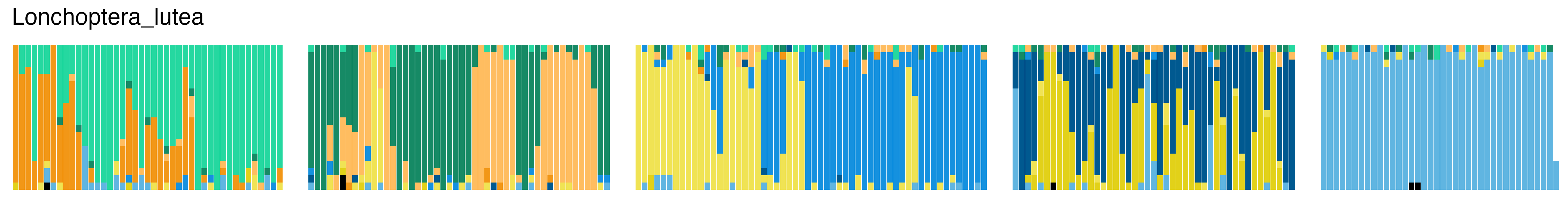

### 35_Hybotidae.png

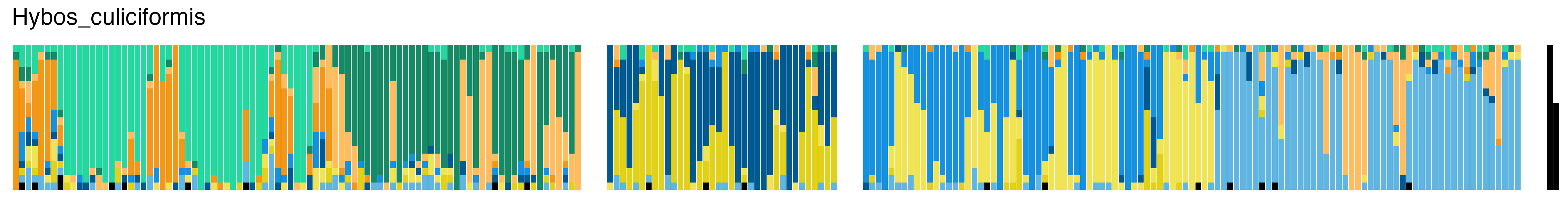

### 36_Dolichopodidae.png

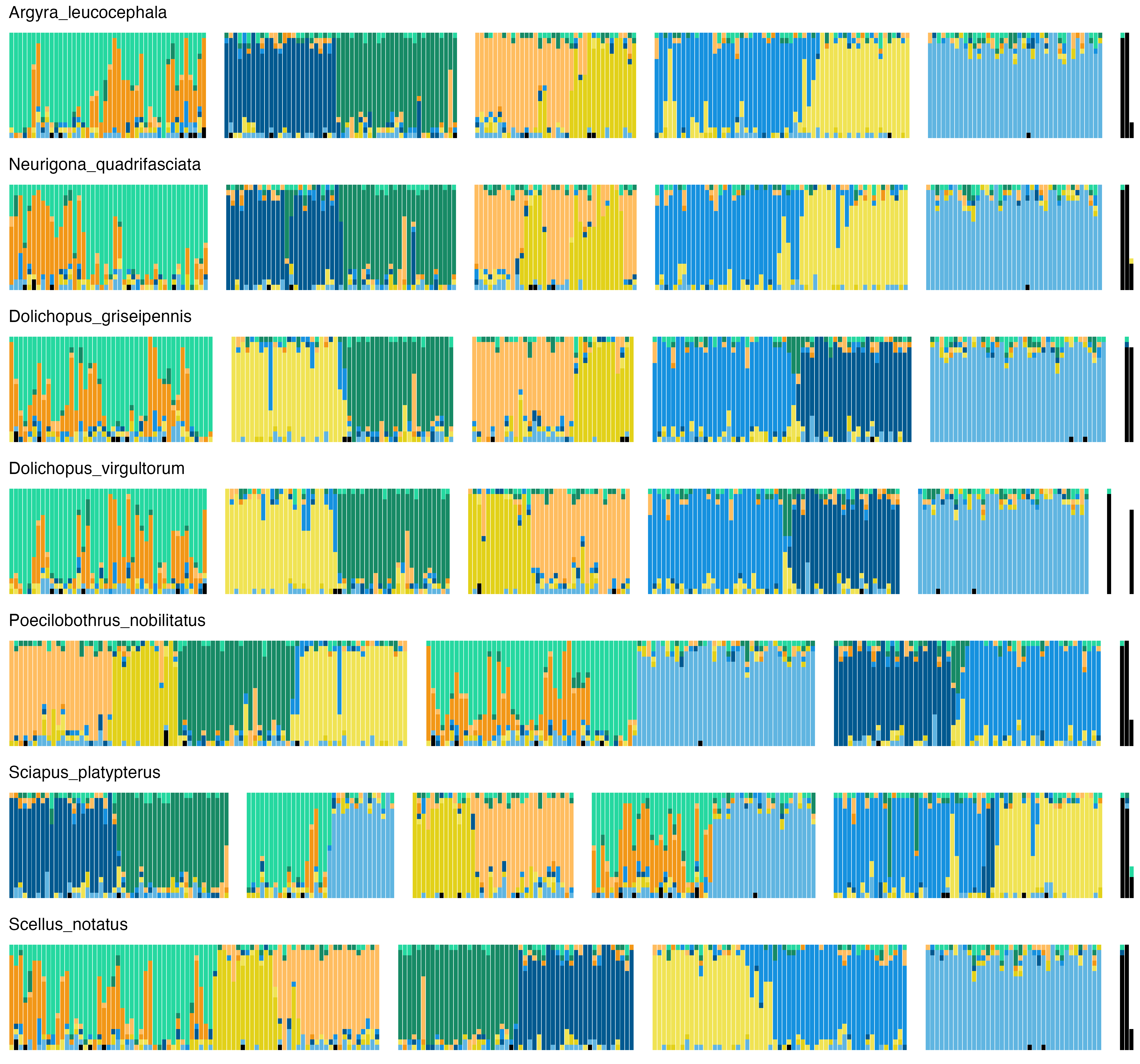

### 37_Empididae.png

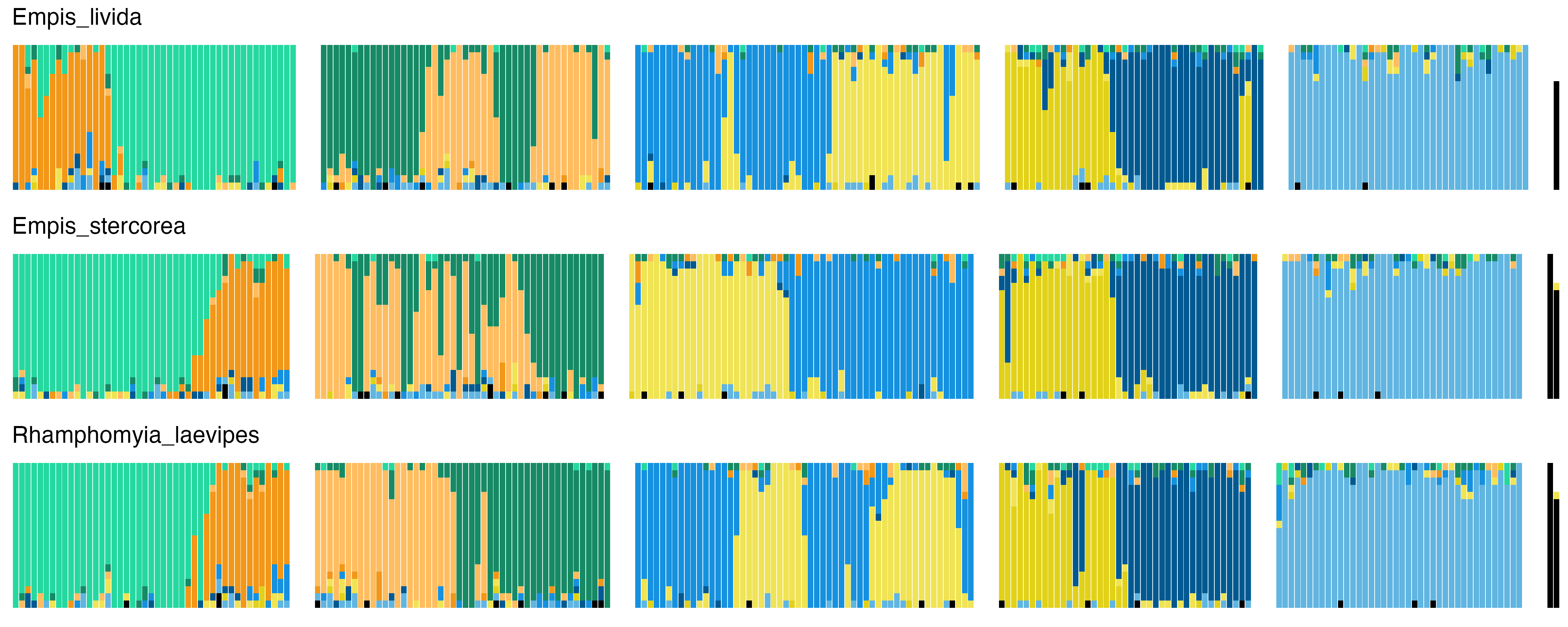

### 38_Therevidae.png

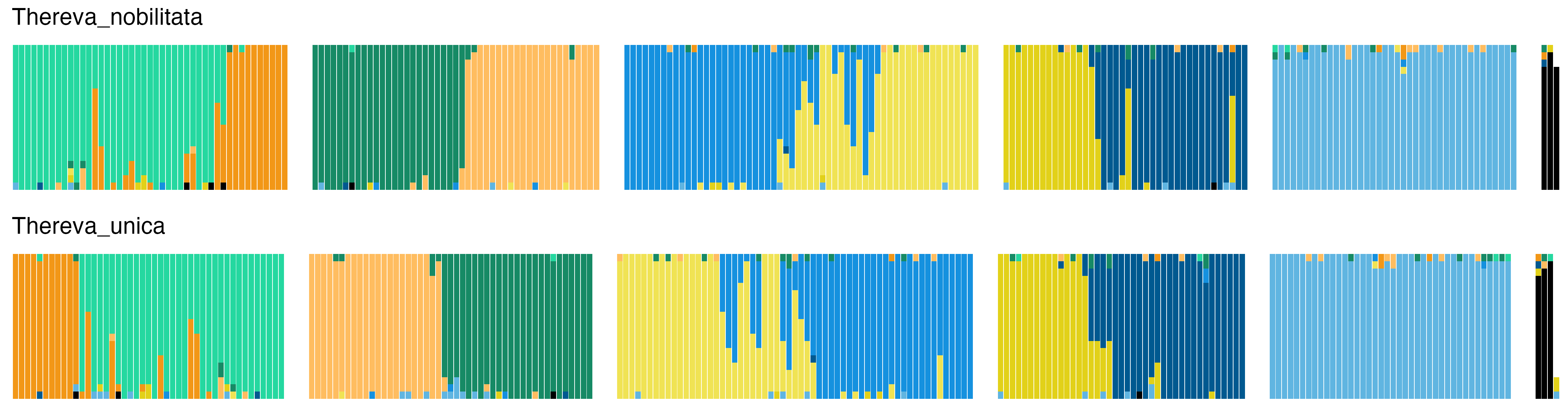

### 39_Asilidae.png

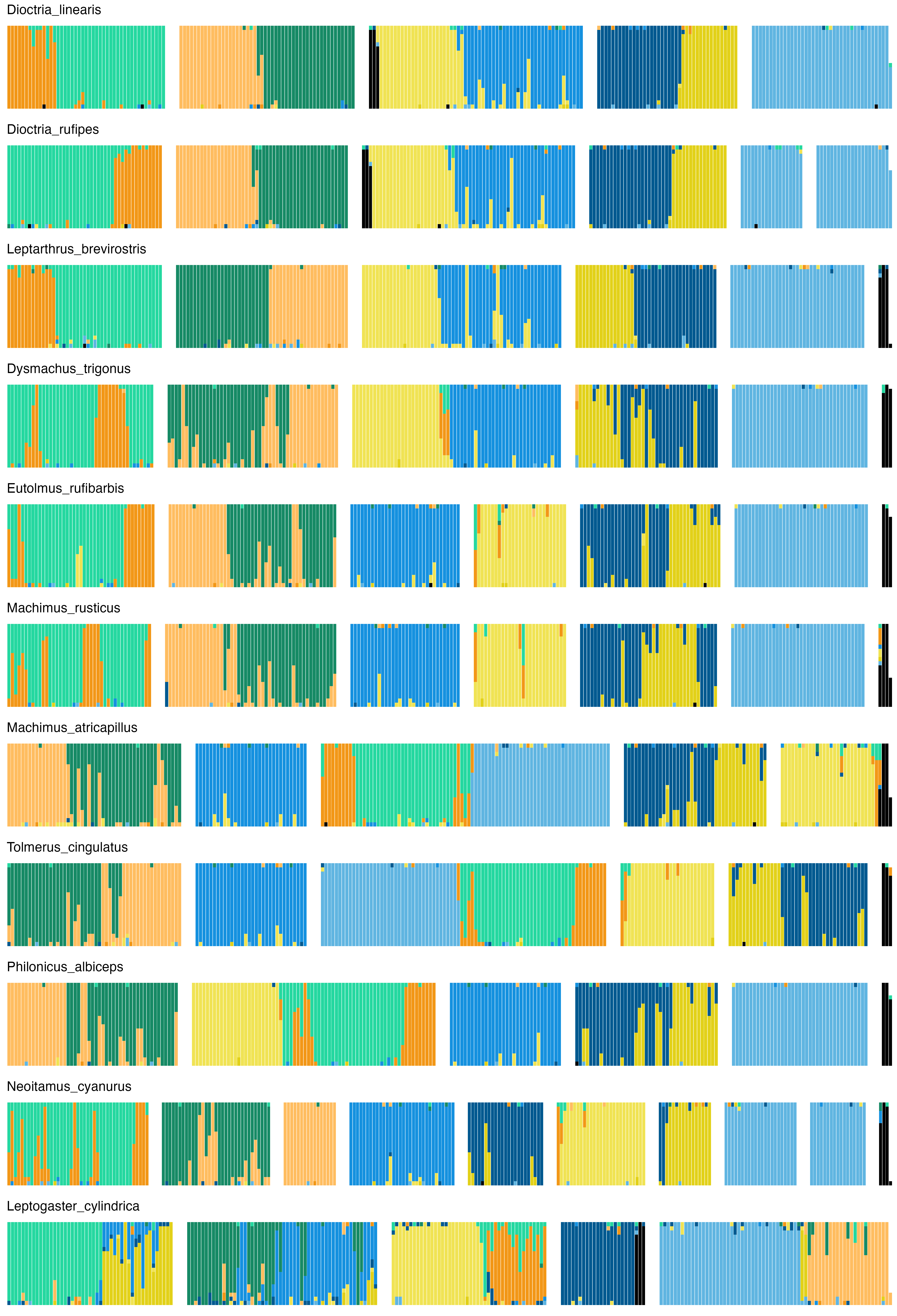

### 40_Bombyliidae.png

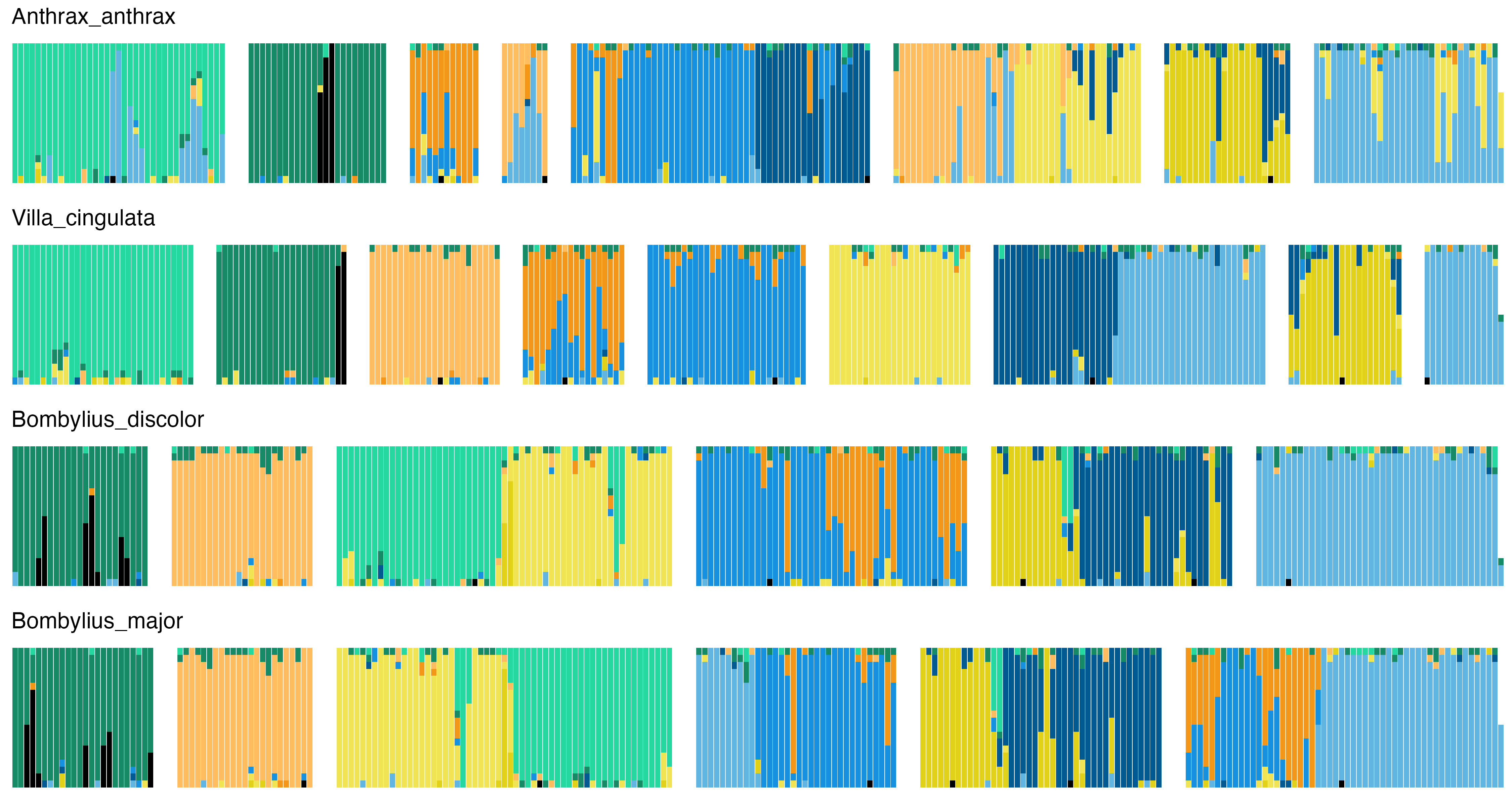

### 41_Xylophagidae.png

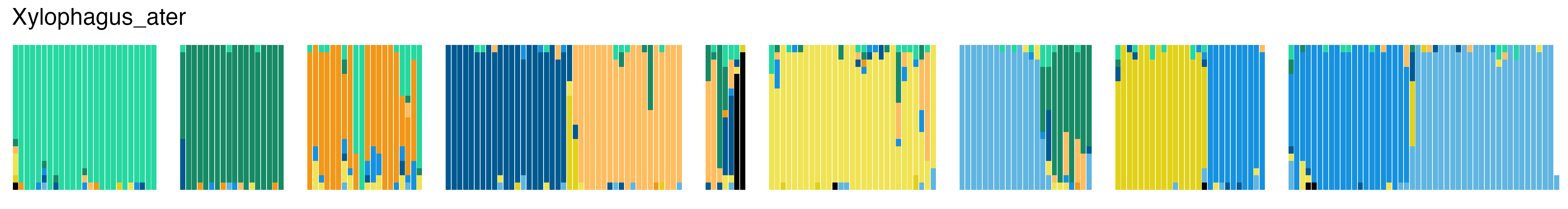

### 42_Acroceridae.png

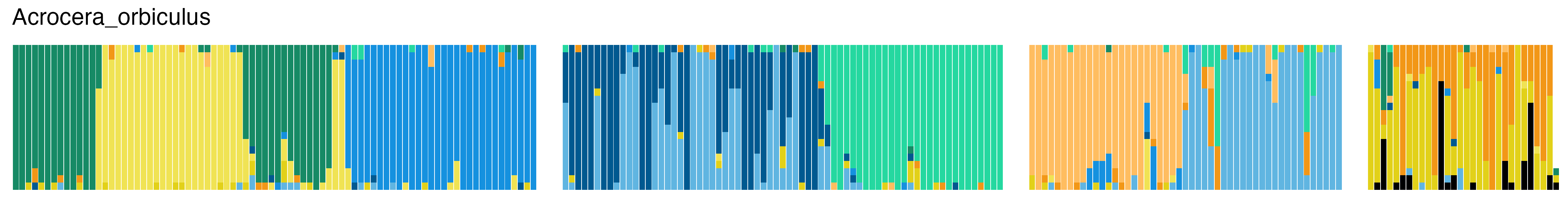

### 43_Stratiomyidae.png

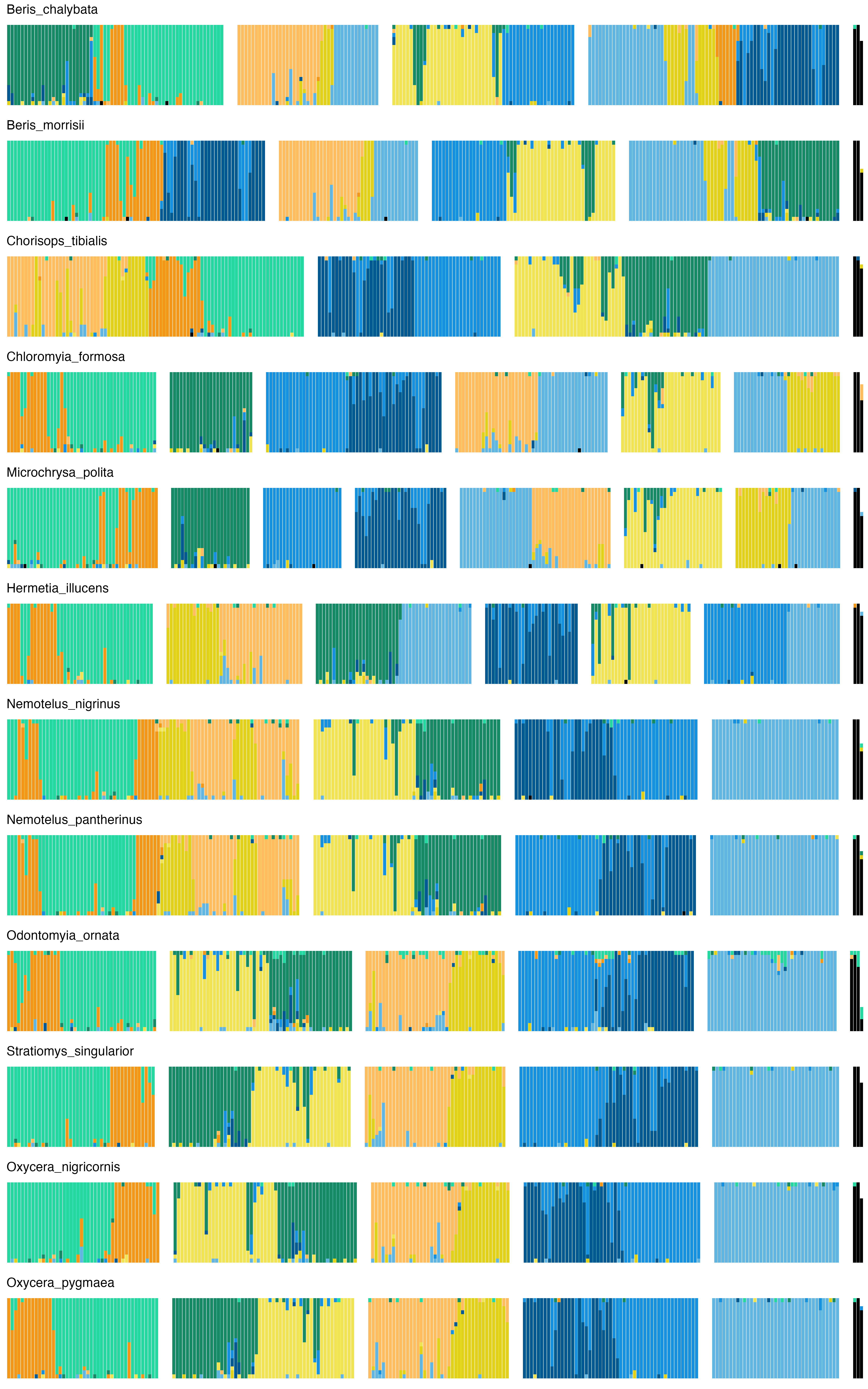

### 44_Tabanidae.png

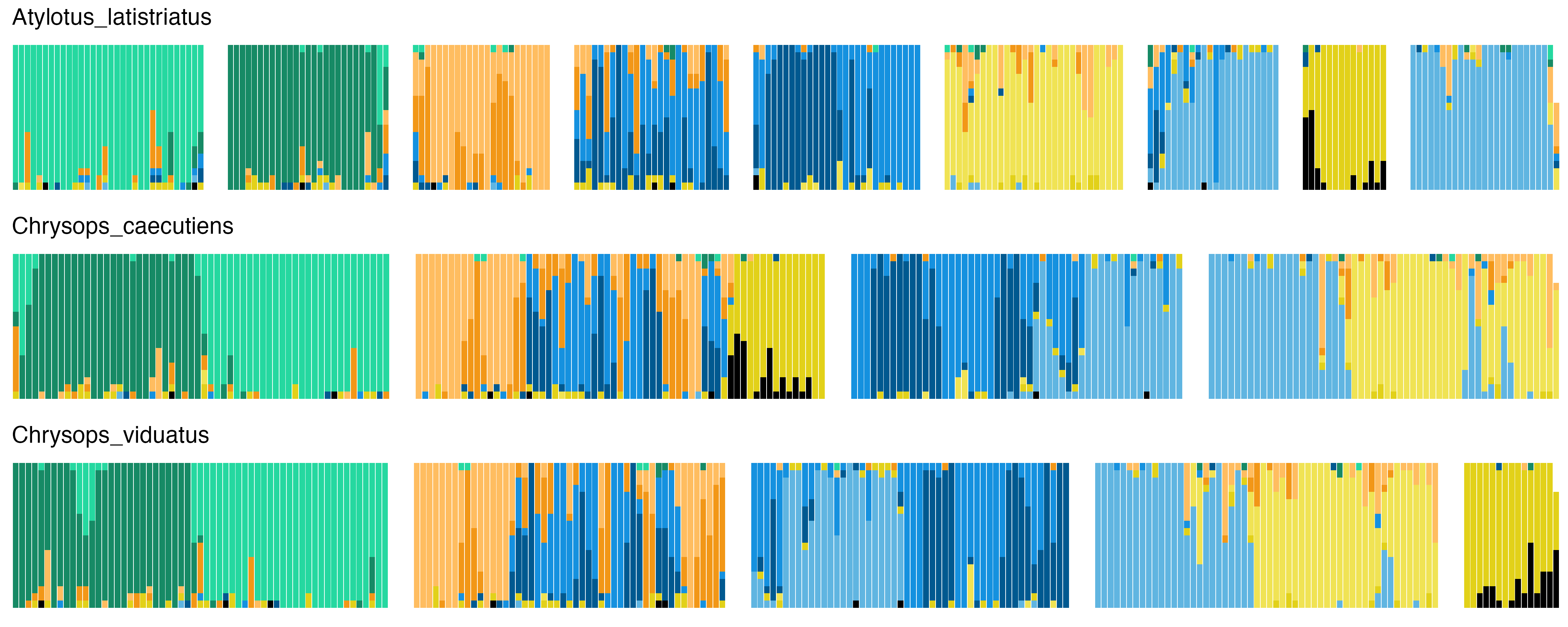

### 45_Athericidae.png

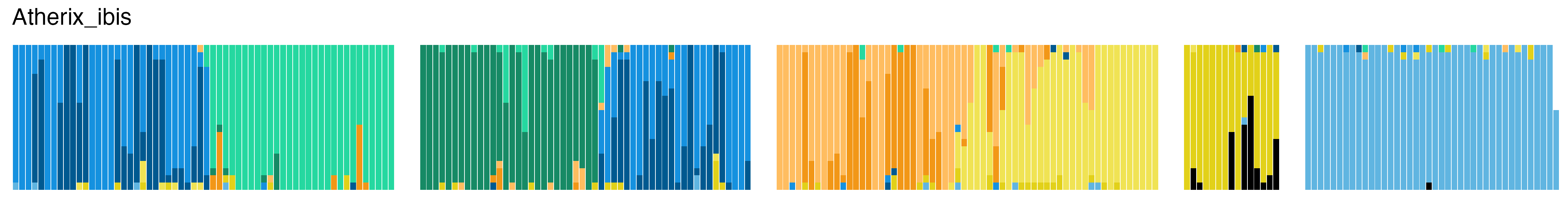

### 46_Rhagionidae.png

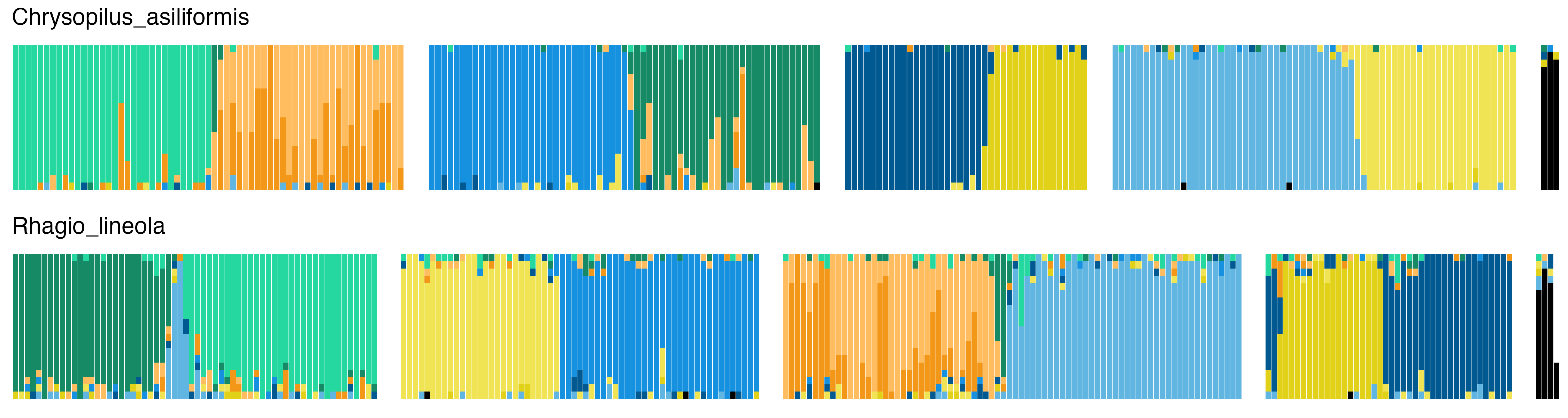

### 47_Mycetophilidae.png

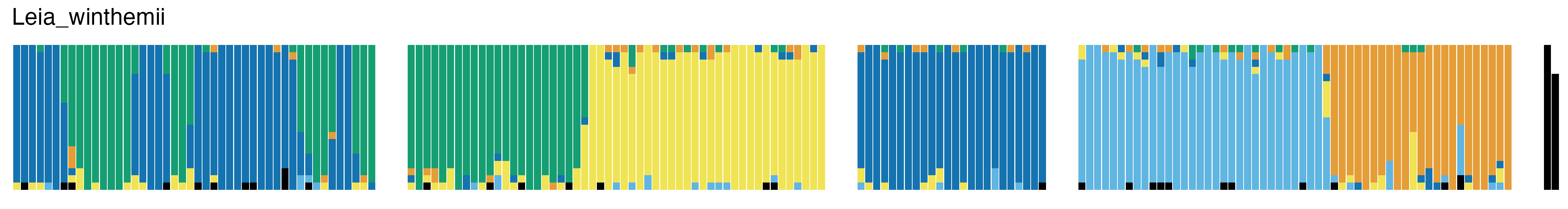

### 48_Sciaridae.png

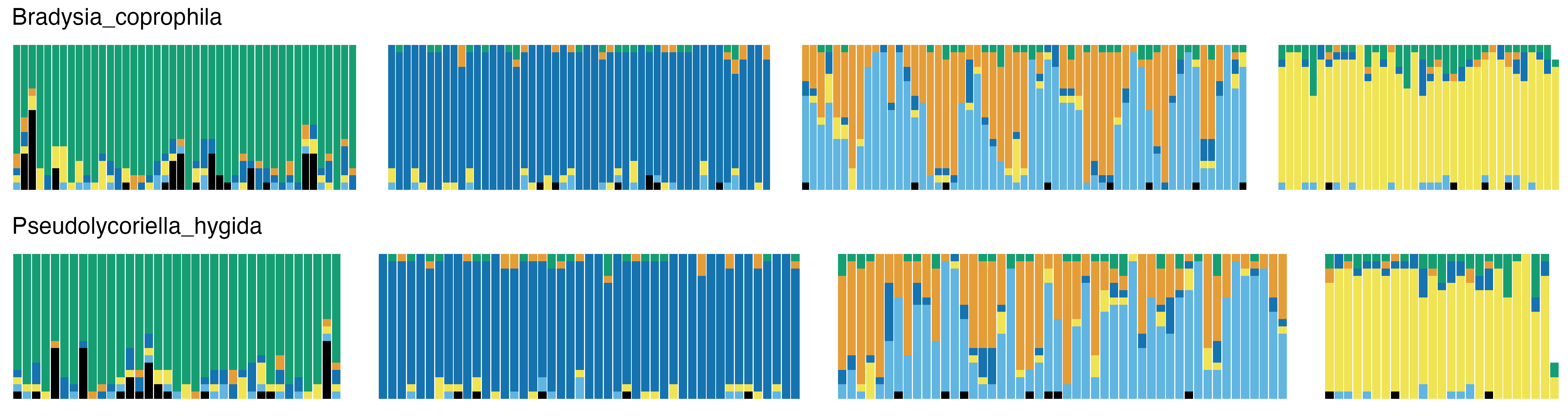

### 49_Cecidomyiidae.png

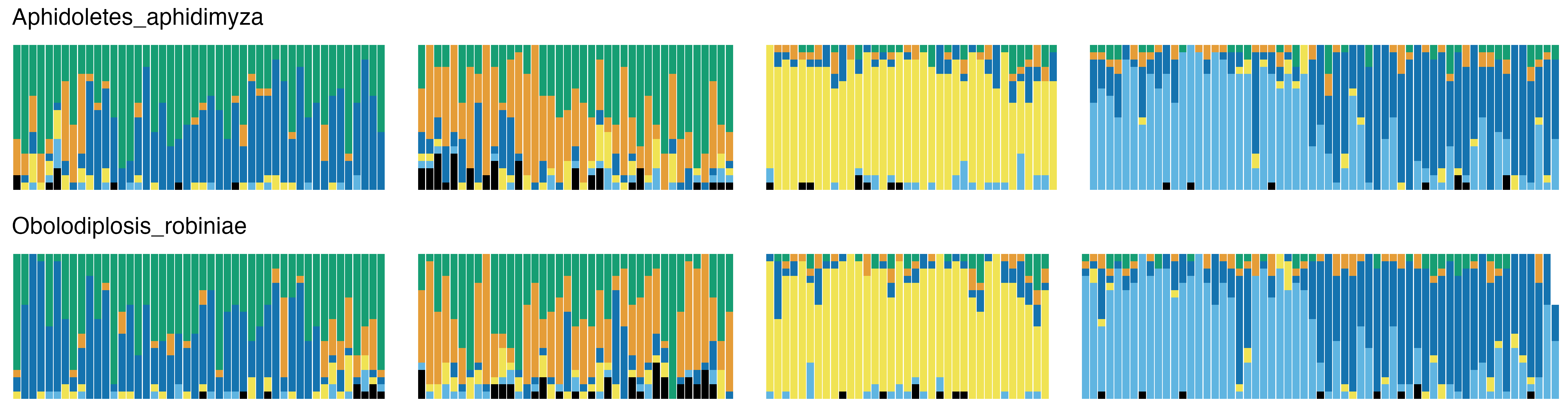

### 50_Bibionidae.png

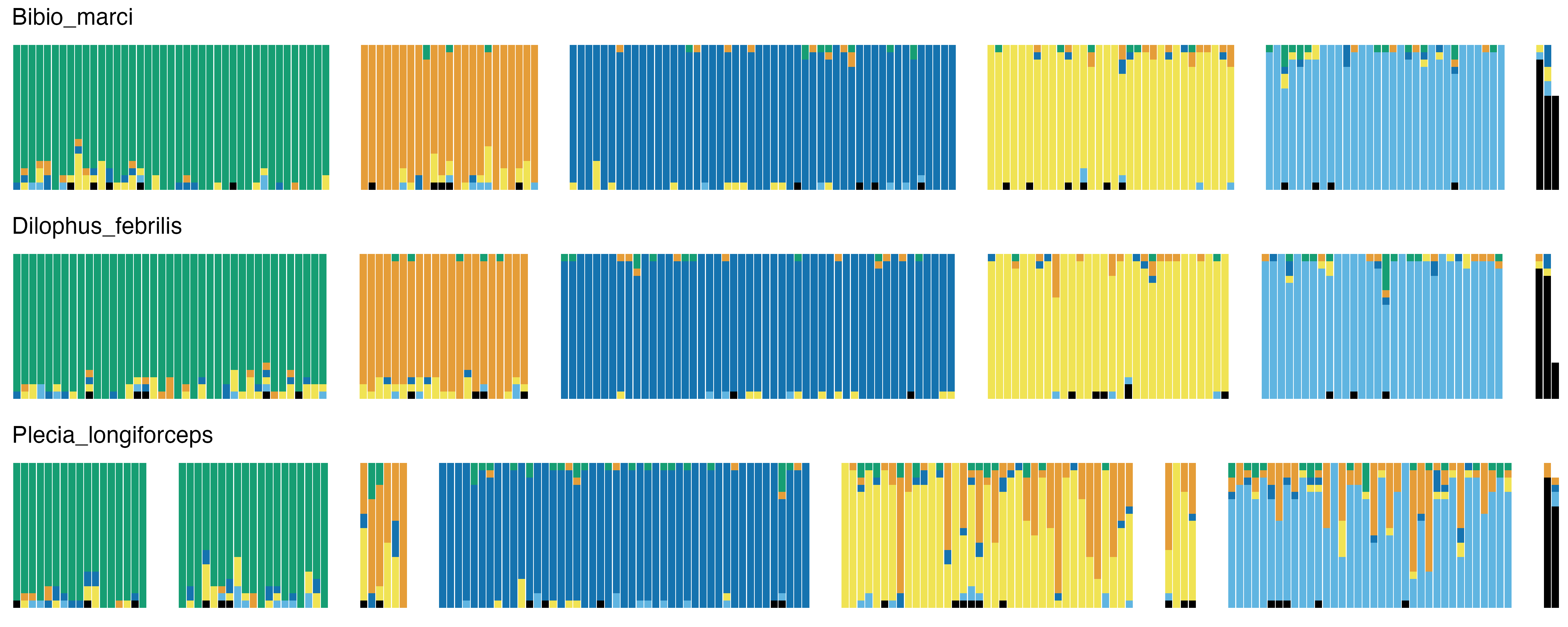

### 51_Anisopodidae.png

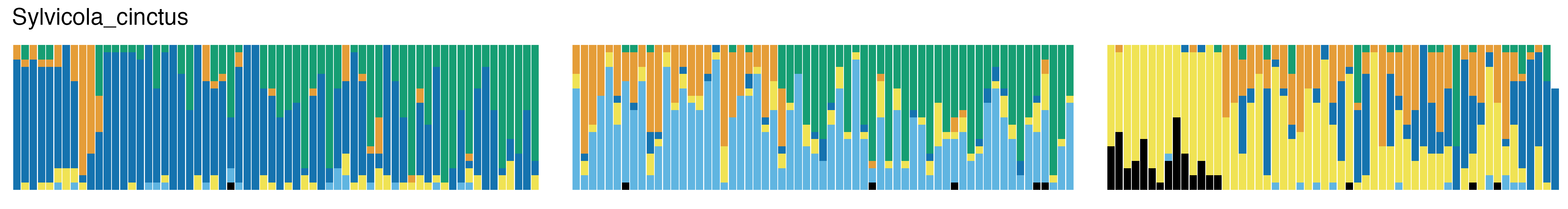

### 52_Tipulidae.png

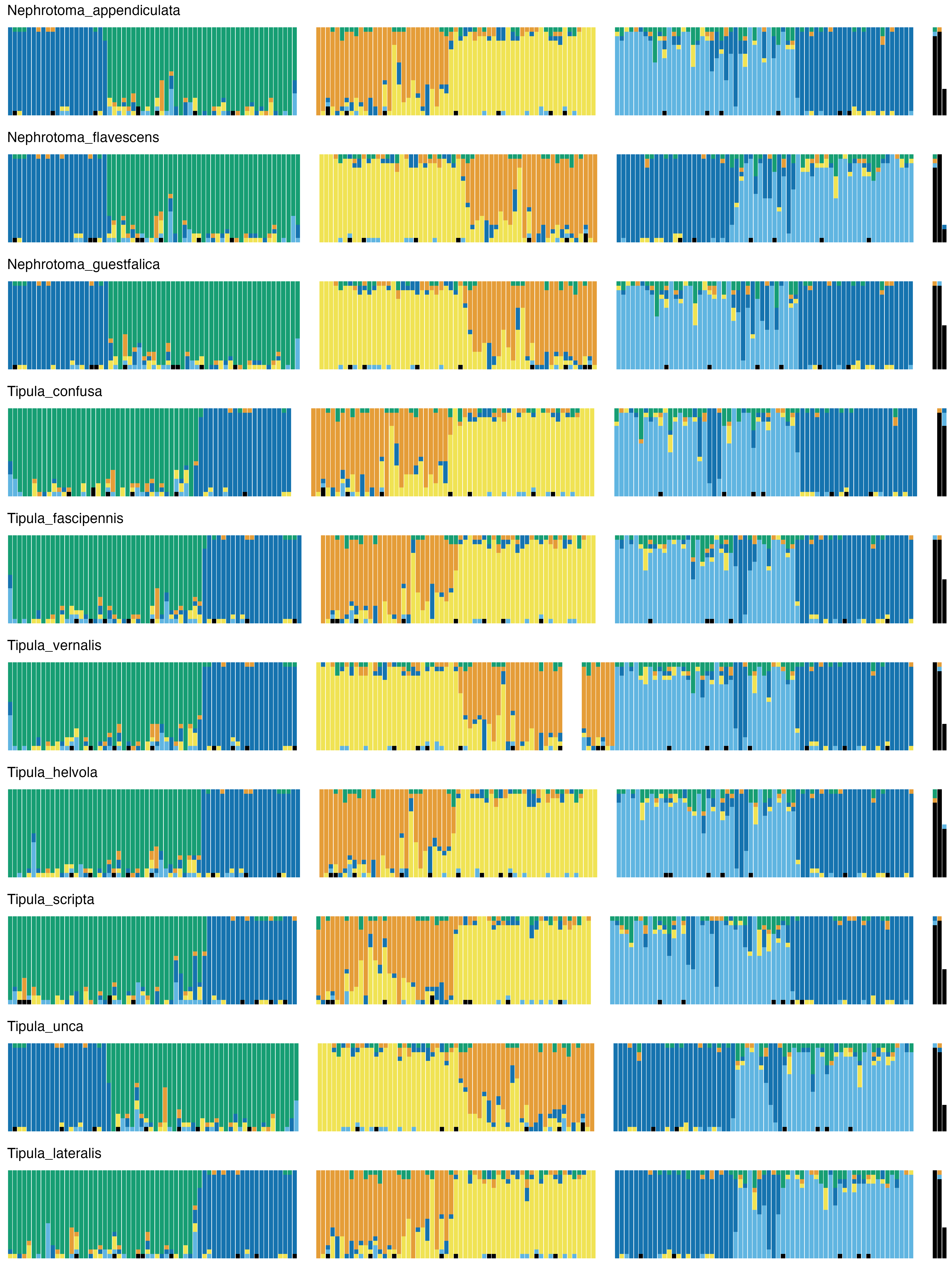

### 53_Cylindrotomidae.png

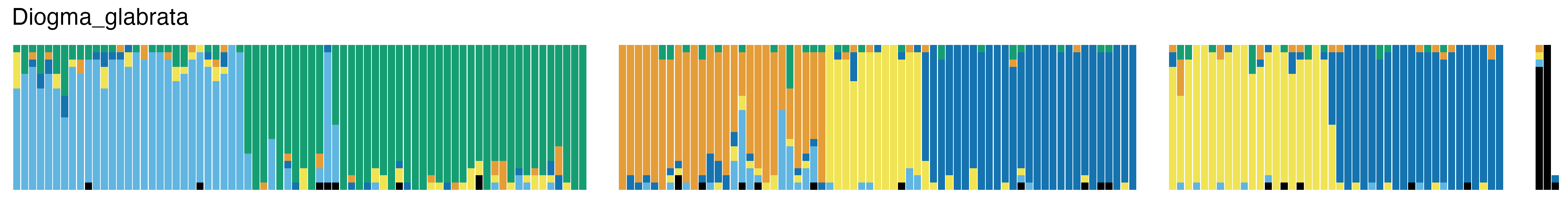

### 54_Pediciidae.png

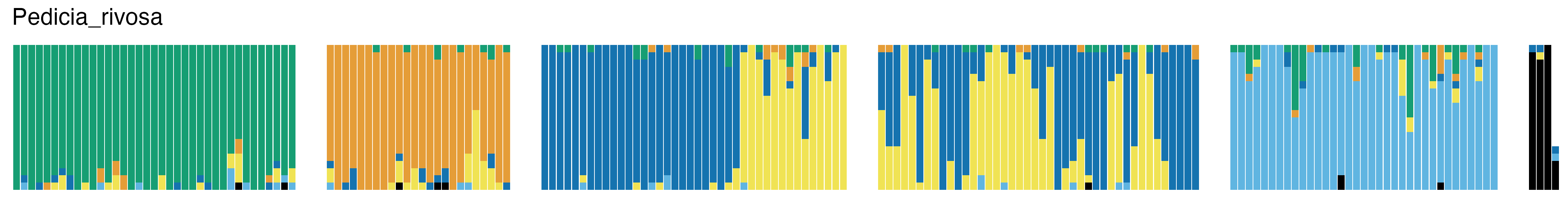

### 55_Ptychopteridae.png

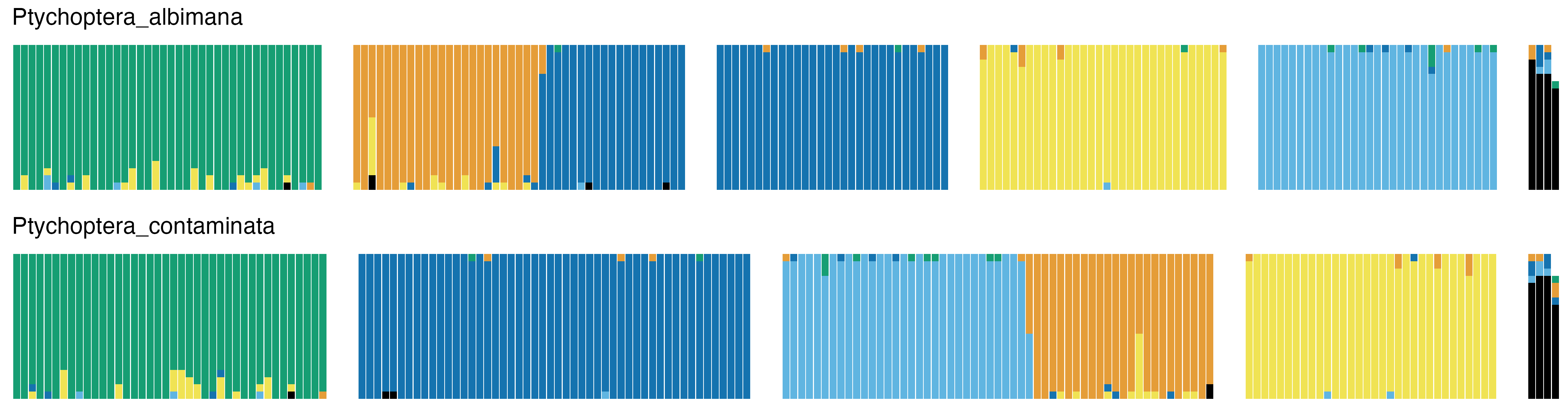

### 56_Psychodidae.png

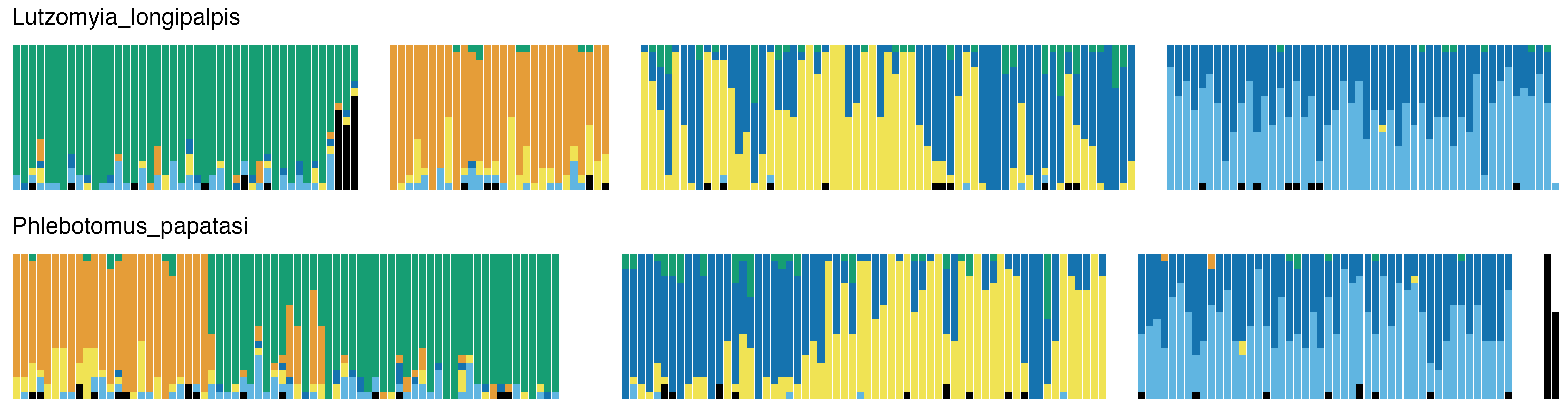

### 57_Ceratopogonidae.png

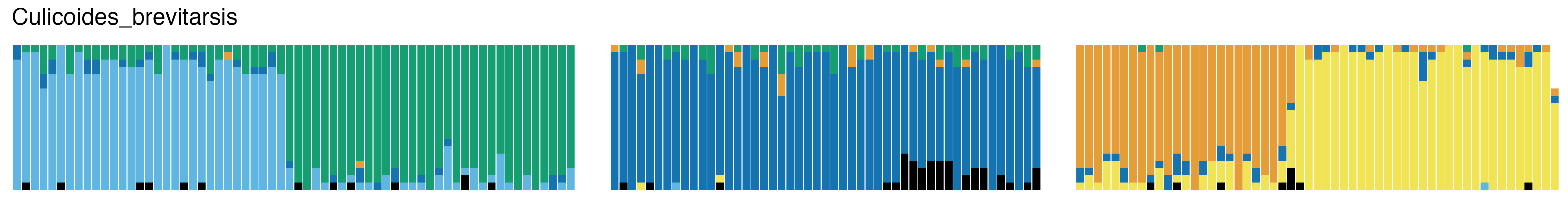

### 58_Chironomidae.png

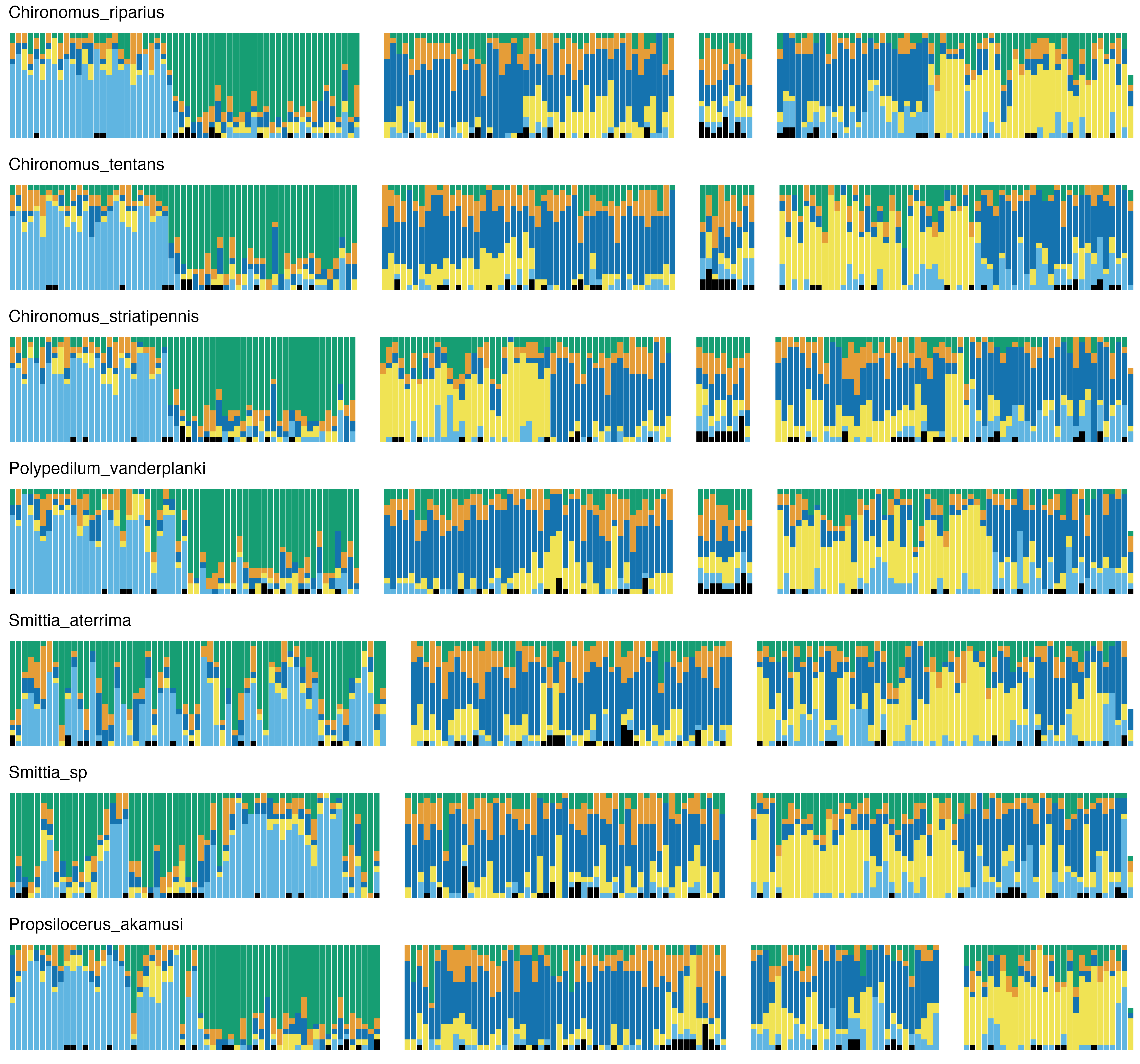

### 59_Culicidae.png

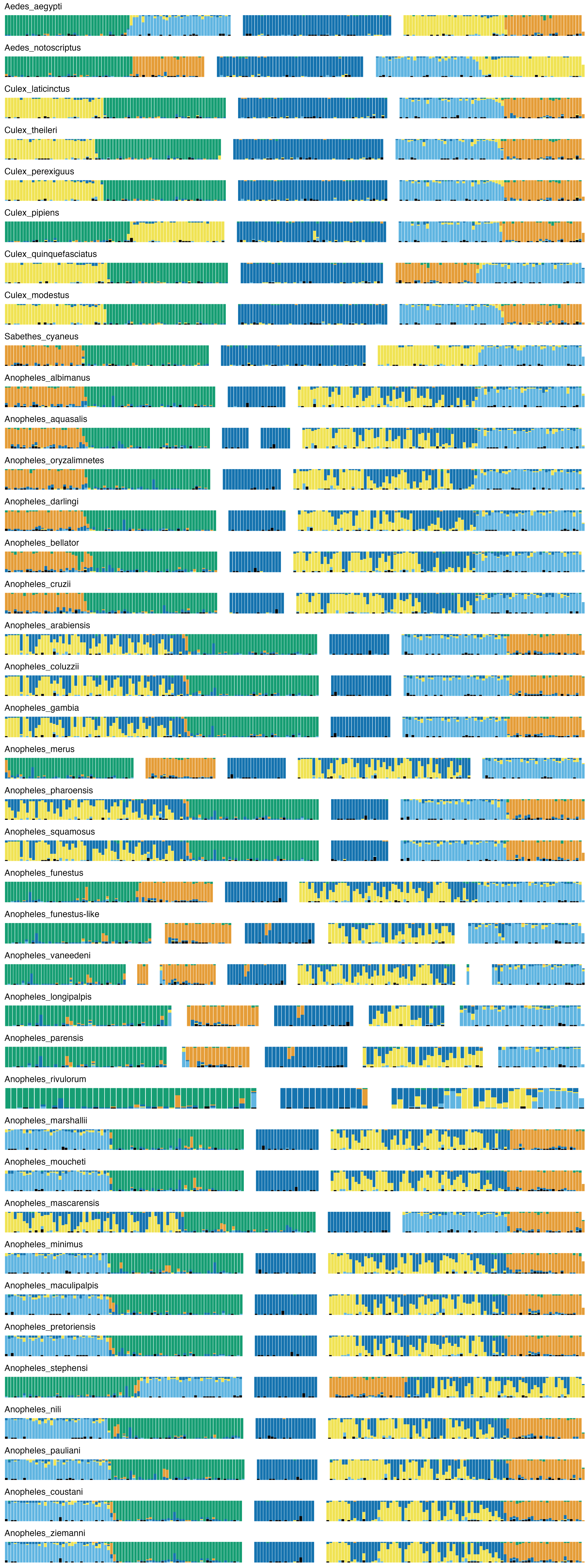

### 60_Panorpidae.png

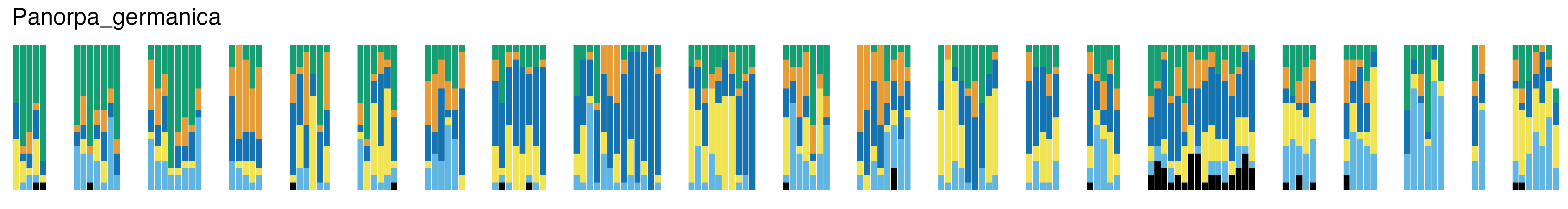
